# Microhomology-mediated end joining acts directly on replication forks to repair single-ended double strand breaks

**DOI:** 10.64898/2026.01.15.699632

**Authors:** Shibo Li, Yuqin Zhao, Youhang Li, Sameer Bikram Shah, Yanmeng Shi, Tran Nguyen, Zi Wang, Chia-Yu Chang, Anagh Ray, Te-Hsuan Bu, Salvatore Loguercio, Takayo Sasaki, Jonathan H. Sussman, Hailong Wang, David M. Gilbert, Mirit I. Aladjem, Xiaohua Wu

**Affiliations:** College of Life Sciences, Tianjin Normal University, Tianjin, 300387, China; Department of Molecular and Cellular Biology, The Scripps Research Institute, La Jolla, CA 92037, USA; Developmental Therapeutics Branch, Center for Cancer Research, National Cancer Institute, Bethesda, MD 20892, USA; Scripps Research Translational Institute, La Jolla, CA 92037, USA; San Diego Biomedical Research Institute, San Diego, CA 92121, USA; College of Life Sciences, Capital Normal University, Beijing 100037, China

**Author notes:** These authors contributed equally (Yuqin Zhao, Youhang Li, Sameer Bikram Shah, Yanmeng Shi).

**Keywords:** Microhomology-mediated end joining (MMEJ), Break-induced replication (BIR), fork-MMEJ, single-ended double-strand breaks (seDSBs), Polθ, PIF1, ATR, end resection, leading and lagging strands

## Abstract

Replication stress, intrinsic to oncogenesis, often leads to fork breakage and double-strand break (DSB) formation. Conventionally, break-induced replication (BIR) is considered the primary mechanism for repairing replication-associated single-ended DSBs (seDSBs). Here, we demonstrate that microhomology-mediated end joining (MMEJ) acts directly to repair seDSBs at broken replication forks (fork-MMEJ), preferentially on the leading strands, and functions cooperatively with BIR. We also showed that while fork-MMEJ is promoted by Polθ, it operates independently of MRE11/CtIP-mediated end resection, relies on RPA, and produces asymmetric deletion patterns, which is distinct from canonical MMEJ (cMMEJ) defined at replication-independent double-ended DSBs (deDSBs). ATR, activated as end resection proceeds, serves as a pivotal switch to suppress fork-MMEJ while promoting BIR. Combined inactivation of ATR and Polθ synergistically kills cancer cells under high replication stress with minimal toxicity to normal cells. Together, our study provides fundamental insights into the MMEJ mechanism and offers new strategies for cancer treatment.

## INTRODUCTION

Replication fork breakage can arise spontaneously during each S-phase due to replication stress^1^ or when replication encounters single strand breaks (nicks), which are among the most common endogenous lesions^2^. Fork breakage often generates single-ended double strand breaks (seDSBs), and break-induced replication (BIR) is thought to play the primary role in repairing them^3,4^. As replication stress is integral to oncogenesis^5–7^, fork breakage-induced DSBs are a major source driving cancer-related genome instability. Microhomology sequences are frequently observed at breakpoints in cancer genomes^8–12^, suggesting the potential involvement of microhomology-mediated end joining (MMEJ) in cancer-related chromosomal rearrangements^13–17^; however, the underlying mechanisms are still elusive.

Canonical microhomology-mediated end joining (cMMEJ) has been primarily studied for repairing double-ended DSBs (deDSBs)^18–21^. cMMEJ requires short-range end resection by MRE11 and CtIP to expose microhomology (MH) sequences for annealing, followed by trimming of non-homologous (NH) tails, DNA synthesis from the 3’ end of MH, and resolution by end ligation (Figure S1A)^19,20^. DNA polymerase theta (Polθ), encoded by the *POLQ* gene, plays a central role in cMMEJ^13–17^. MMEJ is inherently mutagenic and often causes deletions and insertions (indels) accompanied by 1-6 bps of microhomology at the repair junctions^18–21^.

Notably, in mammalian cells, Polθ deficiency results not only in sensitivity to ionizing radiation (IR) but also to DNA damaging agents that induce replication stress^14,22,23^. Polθ binds to replication forks, and this association increases under replication stress^23–27^. Replication stress-induced DSBs arising in S/G2, can persist into mitosis and be repaired by Polθ-mediated MMEJ^28–31^. Moreover, Polθ is involved in filling post-replicative single-stranded DNA (ssDNA) gaps and promoting microhomology-mediated gap skipping in BRCA-deficient cancer cells^24,25^. In *C. elegans*, Polθ-mediated cMMEJ participates in the post-replicational repair of broken forks following fork convergence or second round replication (Figure S1B)^22,32–35^. However, although Polθ plays a key role in the replication stress response, it remains unknown whether MMEJ can be used directly to repair seDSBs upon fork breakage.

Fork breakage at nicks on the leading versus lagging strands generates different DSB structures. Leading-strand breakage produces seDSBs through Cdc45/MCM/GINS (CMG) run-off, whereas lagging-strand breakage often converts seDSBs to deDSBs as CMG bypasses nicks^36,37^, although seDSBs are also detected on broken lagging strands^38^. Further studies revealed that Cas9 nickase (Cas9n)-induced seDSBs on lagging strands contain 3’ ssDNA overhangs, whereas those on leading strands are typically blunt-ended^38^. The difference in DSB end structures raises an intriguing question of whether repair mechanisms are modulated differently on broken leading and lagging strands. In yeast, acetylation modification is required only on the broken leading strands to facilitate HR repair^37^. Notably, while HR is involved in repairing DSBs on both strands in mammalian cells, it appears to be more strongly activated on the broken lagging strands^39^ and more dependent on BRCA1 than on the broken leading strands^40^; however, the underlying mechanism driving this difference remains unknown.

In this study, we discovered a new activity for MMEJ to directly repair seDSBs on broken forks (fork-MMEJ) (Figure S1C left), with a mechanism substantially different from cMMEJ at deDSBs. This unique fork-MMEJ is independent of end resection, operates more frequently on the broken leading strands, and functions together with BIR to repair seDSBs. We also uncovered an elegant regulatory control mechanism by ATR that governs the use of fork-MMEJ and BIR.

## RESULTS

### MMEJ is induced upon fork breakage after Cas9n cleavage

We inserted our established EGFP-MMEJ reporter^41^ (Figure 1A top) into the *AAVS1* locus in U2OS cells, a region flanked by bidirectional replication origins as revealed by nascent strand sequencing^42,43^ (Figure S2A). MMEJ induced by I-SceI in U2OS (EGFP-MMEJ-AAVS1) cells was Polθ-dependent (Figure S2B). To study MMEJ repair at broken forks, we used gRNA2 (g2)/Cas9^D10A^ to introduce a nick on the top strand of the EGFP-MMEJ reporter, located 1 bp outside of the left MH (Figure 1A top). When replication progresses from either the right or left toward the EGFP-MMEJ reporter and encounters the nick, fork breakage occurs either on the leading strand, producing seDSBs (lead-seDSBs), or on the lagging strand, generating seDSBs that are subsequently converted to deDSBs (lag-seDSBs/deDSBs)^36,38^ (Figure S1D). Thus, g2/Cas9^D10A^-induced nicks are expected to generate a mixture of lead-seDSBs and lag-seDSBs/deDSBs in the reporter cell population.

**Figure 1.**
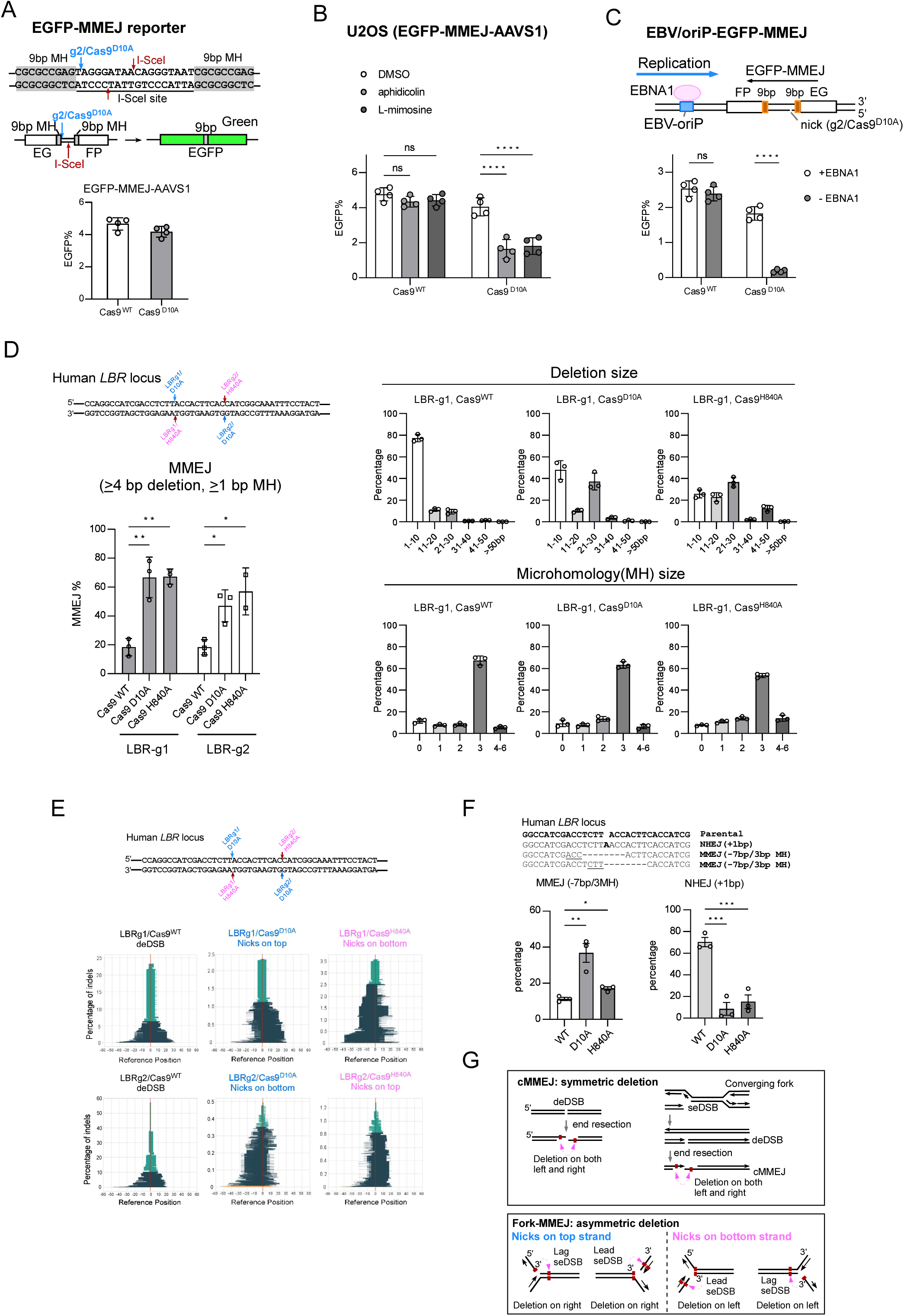
MMEJ is induced upon fork breakage after Cas9 nickase (Cas9n) cleavage. (**A**) Schematic drawing of the EGFP-MMEJ reporter with indications of microhomology (MH) and gRNA2/Cas9^D10A^ and I-SceI cleavage sites (top). MMEJ was assayed in U2OS cells with the EGFP-MMEJ reporter in *AAVS1* locus (EGFP-MMEJ-AAVS1) four days after infection with g2/Cas9^WT^ or g2/Cas9^D10A^ lentiviruses (bottom). (**B**) U2OS (EGFP-MMEJ) cells with or without aphidicolin (0.4 μM) or L-mimosine (0.5 mM) treatment were assayed for MMEJ. (**C**) Schematic drawing of the EBV/oriP-EGFP-MMEJ reporter plasmid (top). U2OS cells containing EBV/oriP-EGFP-MMEJ plasmids with or without EBNA1 expression were analyzed for MMEJ (bottom). (**D**) and (**E**) Two gRNAs (LBR-g1 and LBR-g2) targeting the human *LBR* locus were used along with Cas9^WT^, Cas9^D10A^ or Cas9^H840A^ by lentiviral infection to U2OS cells, followed by deep sequencing analysis of the cleavage sites. The percentage of MMEJ (≥4 bp deletion and ≥1 bp MH) events (left) and the distribution of deletion size and MH size (right) were determined **(D)**. The percentage of repair events with indels is plotted (y-axis), with the x-axis indicating the deletion size relative to the Cas9 cleavage sites (0 on x-axis) **(E)**. The colors represent the size of deletions: light green (1-10 bp), dark green (11-50 bp), orange (51-100 bp), and yellow (≥ 100 bp). (**F**) The frequencies of typical MMEJ (7 bp deletion with 3 bp MH) and NHEJ (1 bp insertion) repair products from **(E)** are shown. (**G**) Schematic illustration of expected deletion patterns of indels generated by cMMEJ (symmetric deletions, top) at deDSBs and fork-MMEJ (asymmetric deletions, bottom) at seDSBs on forks.

We observed robust MMEJ activity upon g2/Cas9^D10A^ cleavage of the EGFP-MMEJ-AAVS1 in U2OS cells, comparable to that at deDSBs after g2/Cas9^WT^ cleavage (Figure 1A bottom). This nick-induced MMEJ (g2/Cas9^D10A^) was suppressed by aphidicolin and L-mimosine, which inhibit replication (Figure 1B). Likewise, multiple U2OS clones with randomly integrated EGFP-MMEJ reporter in the genome exhibited similar robust g2/Cas9^D10A^-induced MMEJ (Figure S3A), which was aphidicolin sensitive (Figure S3B). Moreover, when the reporter was placed on a plasmid containing the Epstein-Barr virus (EBV) replication origin (oriP) that requires EBNA1 for replication^44–46^, g2/Cas9^D10A^-induced MMEJ depended on EBNA1 (Figure 1C). These findings support a requirement for replication in converting nicks into DSBs to activate fork-MMEJ (Figure S1C left).

To confirm DSB formation in the EGFP-MMEJ reporter after Cas9n cleavage, we modified DSBCapture^47^ using reporter-specific internal primers (Figure S4A). DSBs were detected after g2/Cas9^D10A^ cleavage of the EGFP-MMEJ reporter, whereas genomic DNA without cleavage showed almost no signals (Figure S4B). We also used light-activated Cas9 system (vfCRISPR) with caged gRNA^48^ to induce Cas9n cleavage in U2OS (EGFP-MMEJ-AAVS1) cells arrested in G1 by Palbociclib. We showed that DSBs formed only in cells progressing to S-phase (8 hours after release), but not in G1 arrested cells by droplet digital PCR (ddPCR) (Figure S4C), supporting the requirement of replication for converting nicks into DSBs.

### MMEJ at broken forks shows asymmetric features

To assess MMEJ to repair broken forks at genomic loci, we used Cas9n to cleave the *LBR* locus in human cells, where MMEJ was analyzed after Cas9^WT^ cleavage^49^. We used two gRNAs, LBR-g1 and LBR-g2, targeting opposite DNA strands at the *LBR* locus (Figure 1D top). Deep sequencing analysis revealed that indels with MMEJ features (≥ 1bp MH and ≥ 4 bp deletion) were enriched following Cas9^D10A^ and Cas9^H840A^ cleavage compared to Cas9^WT^ (Figure 1D left). The size of deletion and MH of resulting indels was greater after cleavage by Cas9^D10A^ and Cas9^H840A^ than those by Cas9^WT^ (Figure 1D right, S5A and S5B).

Intriguingly, while indels induced by Cas9^WT^ exhibited a symmetric pattern, those from Cas9^D10A^ and Cas9^H840A^ cleavage showed an asymmetric pattern—nicking the top strand leads to deletions extending rightward, whereas nicking the bottom strand resulted in deletions extending leftward (Figure 1E). The symmetric deletion pattern of cMMEJ after Cas9^WT^ cleavage is comparable with bidirectional end resection observed at deDSBs (Figure 1G top). Cas9n-induced indels showing asymmetric deletion patterns were diminished when replication was inhibited by aphidicolin (Figure S5C). Therefore, MMEJ at broken forks induced by Cas9n exhibits distinct features compared to cMMEJ at deDSBs induced by Cas9^WT^.

As reported, LBR-g1/Cas9^WT^ cleavage yielded a prominent +1 bp (1 bp insertion) NHEJ product and two major -7 bp (7 bp deletions with 3 bp MH) cMMEJ products^49^. In contrast, we observed that Cas9^D10A^ and Cas9^H840A^ cleavage produced more frequently the -7 bp MMEJ products, whereas the +1 bp NHEJ products were markedly low (Figure 1F), indicating that fork-MMEJ, rather than NHEJ, is utilized to repair broken forks.

### MMEJ induced after Cas9 and Cas9n exhibits different genetic dependence

Like cMMEJ at deDSBs^18,50–53^, Cas9n-induced MMEJ was dependent on Polθ and LIG3 (Figure 2A and S6A). However, contrary to the role of RPA in suppressing cMMEJ^19,51^, expressing RPA2 shRNA significantly reduced fork-MMEJ (Figure 2B and S6B). We tagged RPA2 with dTAG (FKBP12^F36V^)^54^, and found that fork-MMEJ was reduced, while cMMEJ was increased, after inducing RPA2-dTAG degradation in U2OS (EGFP-MMEJ-AAVS1) cells with endogenous RPA2 depleted by shRNAs (Figure S6C). These data suggest that RPA is required for fork-MMEJ. Additionally, in stark contrast to cMMEJ at deDSBs, which relies on MRE11-and CtIP-mediated short end resection to expose MHs^41,55–57^, fork-MMEJ (g2/Cas9^D10A^) was independent of MRE11 and CtIP, and instead, depleting MRE11 and CtIP resulted in an increase in fork-MMEJ (Figure 2C and S6D). Moreover, like cMMEJ, which does not require extensive end resection^41^, fork-MMEJ was not decreased; rather, it was moderately increased after depleting BLM, DNA2 and EXO1 (Figure S6E). We propose that at broken forks, instead of end resection, DNA unwinding by helicase activities may be involved in exposing MHs. We term MMEJ that is used to directly repair fork-associated DSBs as fork-MMEJ. Our study suggests that fork-MMEJ exhibits characteristics that are distinct from that of cMMEJ.

**Figure 2.**
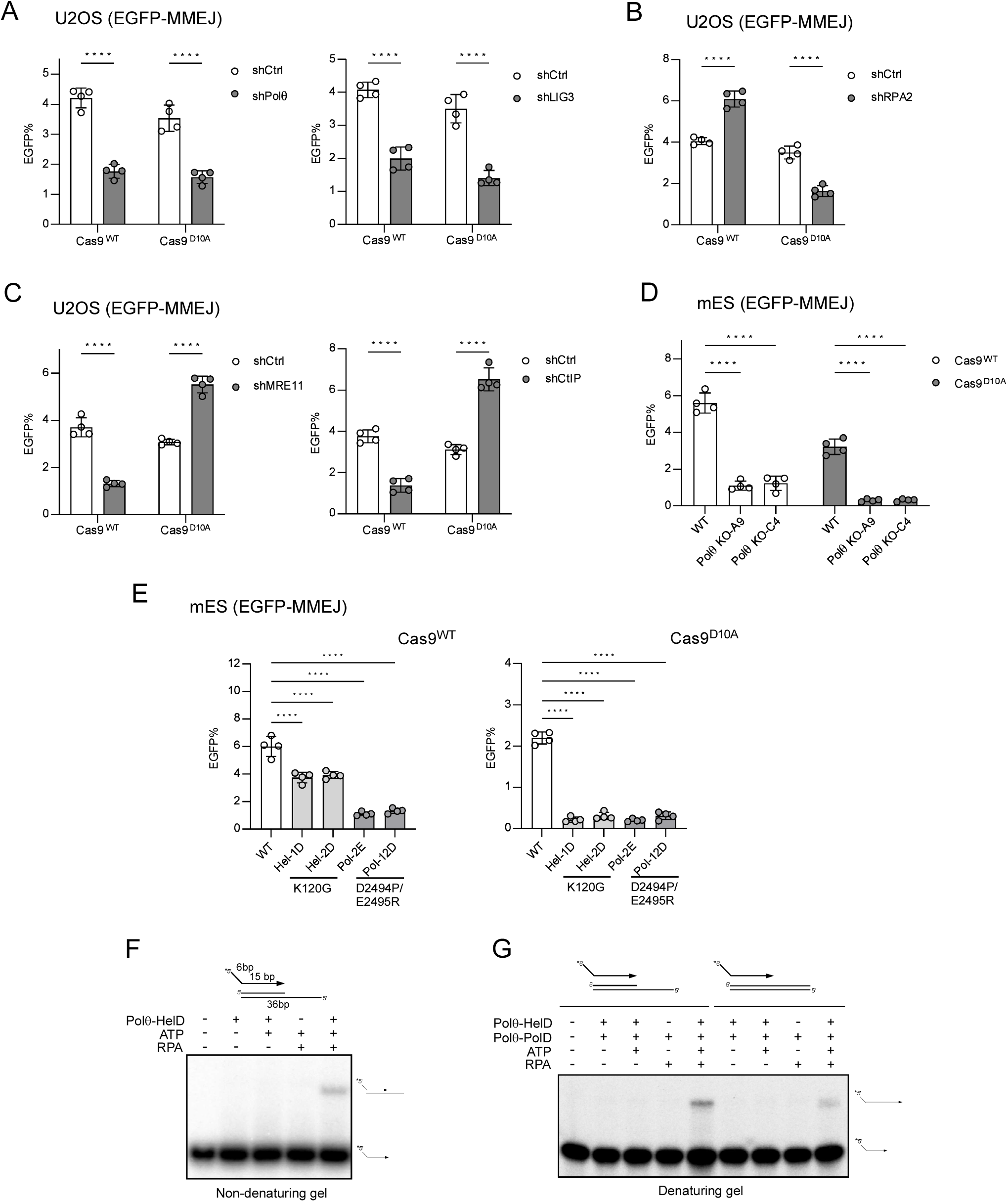
MMEJ induced by Cas9 and Cas9n exhibits distinct genetic dependence. (**A-C**) MMEJ was assayed in U2OS (EGFP-MMEJ) cells expressing shRNAs for Polθ (**A**), LIG3 (**A**), RPA2 (**B**), MRE11 (**C**) or CtIP (**C**) after g2/Cas9^WT^ or g2/Cas9^D10A^ cleavage. (**D**) MMEJ was assayed in WT or *POLQ*-KO mES (EGFP-MMEJ) cells four days after transfection of plasmids encoding g2/Cas9^WT^ or g2/Cas9^D10A^. (**E**) mES (EGFP-MMEJ) cells with *POLQ* WT allele or knock-in (KI) alleles containing mutations of K120G or D2494P/E2495R were assayed for MMEJ after g2/Cas9^WT^ or g2/Cas9^D10A^ cleavage. (**F**) *In vitro* biochemical assay showing Polθ-HelD activity in unwinding dsDNA to facilitate strand exchange with ssDNA carrying 15 bp homology. ^32^P-5’-labeled ssDNA was mixed with the indicated dsDNA substrate and incubated with Polθ-HelD, ATP and/or RPA, and the reaction products were resolved on a non-denaturing gel. (**G**) *In vitro* biochemical assay showing strand displacement DNA synthesis by Polθ-HelD and Polθ-PolD. ^32^P-5’-labeled ssDNA was mixed with the indicated dsDNA substrates and incubated with Polθ-HelD, ATP and/or RPA, followed by DNA extension with Polθ-PolD with 0.1 mM dNTPs, and the reaction products were resolved on a denaturing gel.

We introduced the EGFP-MMEJ reporter to mouse embryonic stem cells (mES) cells at the *ROSA* locus, where replication also proceeds bidirectionally towards *ROSA*^58^ (Figure S7A) and showed that *POLQ*-KO led to a strong reduction in both fork-MMEJ and cMMEJ (Figure 2D and S7B). To test the role of the polymerase and helicase activities of Polθ in fork-MMEJ, we generated knock-in mutants of *POLQ*-K120G (helicase mutant) and D2494P/E2495R (polymerase mutant) in the Polθ helicase domain (HelD) and polymerase domain (PolD), respectively^53^. cMMEJ (g2/Cas9^WT^) was drastically reduced in the polymerase mutant D2494P/E2495R, comparable to that in *POLQ*-KO, whereas the helicase mutant K120G exhibited 30-40% reduction in cMMEJ (Figure 2E left), consistent with the involvement of the Polθ helicase activity for removing RPA from ssDNA to promote cMMEJ^19^. However, after g2/Cas9^D10A^ cleavage, both D2494P/E2495R and K120G mutants showed a severe defect in fork-MMEJ, comparable to *POLQ*-KO (Figure 2E right). Thus, fork-MMEJ relies more on Polθ helicase activity compared to cMMEJ. We speculate that Polθ-HelD is involved in DNA unwinding to expose MHs for fork-MMEJ at seDSBs (Figure 7G right), in addition to its role in antagonizing RPA^19^.

It has been described that Polθ-HelD exhibits activity to unwind DNA *in vitro*, which is facilitated by RPA (Figure S8A)^59^. We further showed that Polθ-HelD unwound DNA duplex, allowing ssDNA to anneal the template in an RPA-facilitated and ATP-dependent manner (Figure 2F and S8B). Polθ-PolD could then utilize the 3’ end of the newly annealed DNA strand to start DNA synthesis, even under conditions requiring strand displacement synthesis, which also required ATP and RPA (Figure 2G and S8C).

To examine whether Polθ is recruited to seDSBs in cells, we performed chromatin immunoprecipitation (ChIP) analysis after Cas9^D10A^ cleavage in U2OS (EGFP-MMEJ) cells. Polθ was recruited to seDSBs as efficiently as to deDSBs using γH2AX ChIP to mark DSB formation (Figure S8E), supporting the role of Polθ in fork-MMEJ.

### Fork-MMEJ is strongly suppressed by 2 bp or longer NH tails 3’ to the MH

To examine whether MH size influences MMEJ frequency, we generated a set of MMEJ reporters using 2-6 bp MHs for annealing and found that longer MHs promoted higher frequencies of both c-MMEJ and fork-MMEJ (Figure S9A). Notably, fork-MMEJ remains relatively more efficient as MH length decreases compared to c-MMEJ.

While nicking the top strand of the EGFP-MMEJ reporter at the *AAVS1* locus in U2OS cells with g2/Cas9^D10A^ induced a substantial level of fork-MMEJ (Figure 1A), surprisingly, nicking the bottom strand with g2/Cas9^H840A^ failed to do so (Figure 3A left). Similar results were obtained when we used g2/Cas9^D10A^ and g2/Cas9^DH840A^ to cleave the EGFP-MMEJ reporter inserted at the *ROSA* locus in mES cells (Figure 3A right). Depending on the replication direction, nicking the top or bottom DNA strands would result in fork breakage on leading or lagging strands. Using the EGFP-MMEJ reporter, cleaving with g2/Cas9^D10A^ (nick top) or g2/Cas9^H840A^ (nick bottom), four possible fork breakage outcomes could be generated and classified into four types: Type I, II, III, and IV (Figure 3B). In a cell population, we anticipate that nicking the EGFP-MMEJ reporter by g2/Cas9^D10A^ (top strand) and g2/Cas9^H840A^ (bottom strand) would result in mixed Type I and II and mixed Type III and IV, respectively (Figure 3A and 3B). We speculate that Type II and III breaks on the leading strands generate seDSBs (lead-seDSBs), while replication bypass could occur at Type I and IV breaks on the lagging strands, converting seDSBs to deDSBs (lag-seDSBs/deDSBs) (Figure 3B and S1D), based on previous studies of fork breakage outcomes^36^.

**Figure 3.**
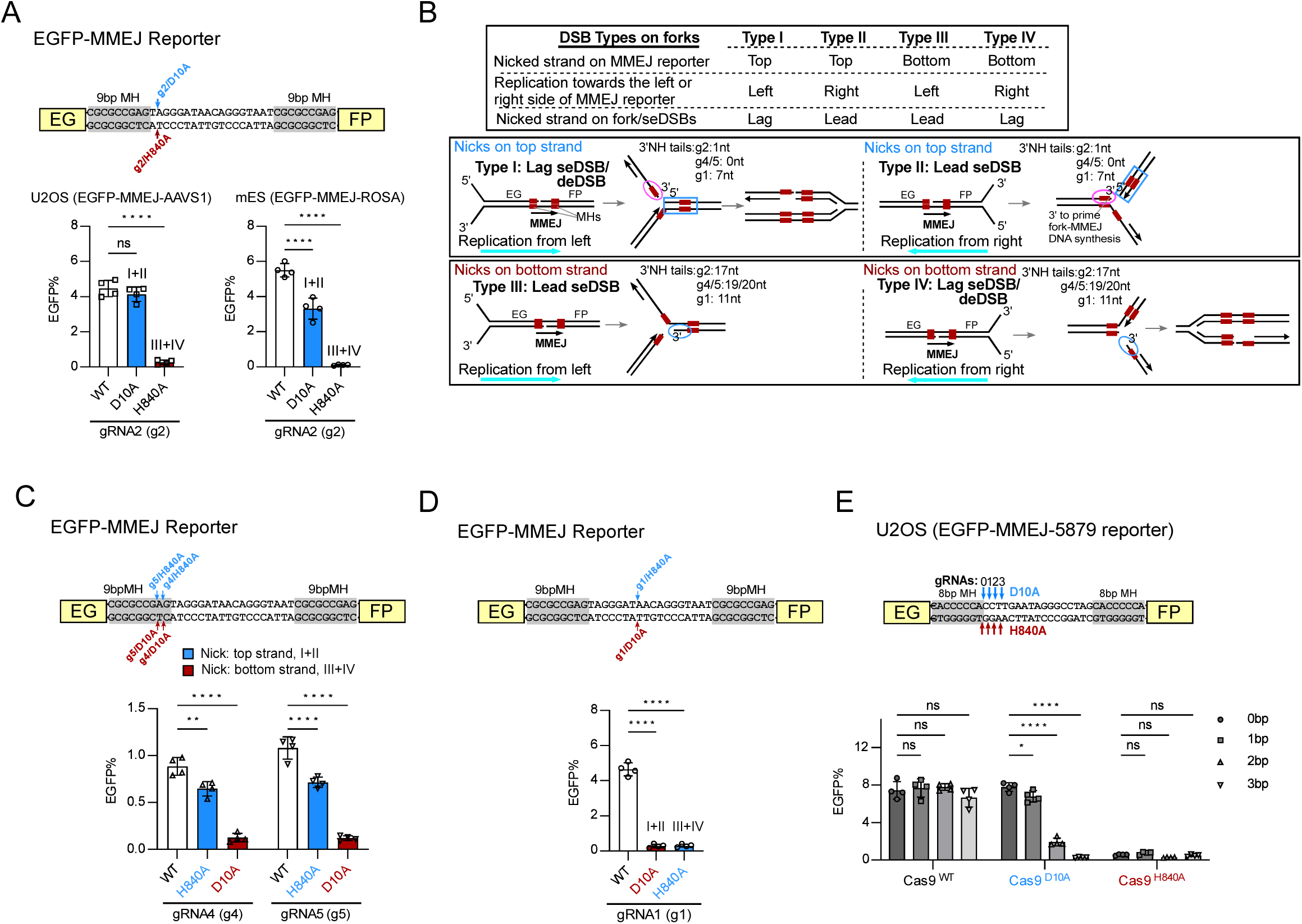
Fork-MMEJ is suppressed by 2 bp or longer NH tails at the 3’ ends of the parental broken strands. (**A**) The microhomology (MH) and gRNA (g2) cleavage site in the EGFP-MMEJ reporter are indicated (top). MMEJ was assayed in U2OS (EGFP-MMEJ-AAVS1) cells and in mES cells with the EGFP-MMEJ reporter inserted in the *ROSA26* locus (EGFP-MMEJ-ROSA) after g2/Cas9^WT^ or g2/Cas9^D10A^ cleavage. (**B**) Schematic drawing of four types of seDSB induced by nicking the top or bottom strand of the EGFP-MMEJ reporter. Lead-seDSBs or lag-seDSBs are indicated. The 3’ ends of the parental broken strands, which are expected to prime fork-MMEJ DNA synthesis, are circles. The length of 3’ non-homologous (NH) tails was indicated for different cleavage sites. (**C, D**) MMEJ was assayed in U2OS (EGFP-MMEJ-AAVS1) cells using the indicated gRNAs with Cas9 and Cas9n. (**E**) U2OS cells carrying the EGFP-MMEJ reporter designed for generating 0-3 bp non-homologous (NH) tails were assayed for MMEJ using the indicated gRNAs with Cas9 and Cas9n.

Besides g2, which cuts 1 bp outside of the left MH, we also generated nicks inside of the left MH with gRNA g4 or g5 (1 bp or 2 bp inside of left MH) in U2OS (EGFP-MMEJ-AAVS1) (Figure 3C). g4 and g5 with Cas9^H840A^ (nick top, I+II) but not with Cas9^D10A^ (nick bottom, III+IV) induced efficient fork-MMEJ. Collectively, we showed that fork-MMEJ can be efficiently induced at Type I and/or II, but not at Type III and IV DSBs on broken forks. One common feature of Type I and II seDSBs after cleavage by g2, g4 or g5 is that the 3’ ends priming for fork-MMEJ DNA synthesis on the broken parental strands contain 0 bp (g4/g5) or 1 bp (g2) NH tails (Figure 3B, pink circles), whereas 3’ ends of Type III and IV seDSBs retain 19/20 bp (g4/g5) or 17 bp (g2) NH tails (Figure 3B, blue circles). We speculate that long 3’ NH tails outside of MH at seDSBs on the broken parental strands would block fork-MMEJ DNA synthesis (Figure S10A, Type III/IV). To further test this, we used g1/Cas9^H840A^ to create Type I/II seDSBs with 7 bp 3’ NH tails (nick top, Figure 3D top and 3B) and found that fork-MMEJ was inefficient at the level comparable to that observed after g1/Cas9^D10A^ cleavage (Figure 3D bottom), which nicked the bottom strand to create Type III/IV breaks with 11 bp 3’ NH tails (Figure 3B). This supports the notion that long 3’ NH tails may inhibit fork-MMEJ.

To investigate further, we designed a new EGFP-MMEJ-5879 reporter where 4 gRNAs generate no (0 bp), 1 bp, 2 bp or 3 bp NH tails to the left MH (Figure 3E top). Cas9^WT^ induced similar levels of cMMEJ at these gRNA sites (Figure 3E bottom), indicating comparable cutting and cMMEJ efficiency regardless of the 3’ NH tail length. Interestingly, however, Cas9^D10A^ induced efficient fork-MMEJ (nick top, Type I+II) when nicks were at the 0 bp site or 1 bp outside of MH (higher efficiency at 0 bp), but not at seDSBs with 2 bp and 3 bp NH tails (Figure 3E bottom). Thus, 3’ NH tails (≥2 bp) outside of MH on the parental strands of broken forks, block fork-MMEJ at seDSBs (Figure S10A), but not cMMEJ at deDSBs. We propose that while cMMEJ involves a trimming process to remove NH tails on both DSB ends (Figure S1A), at broken forks, Polθ may directly initiate DNA synthesis using the 3’ broken end on the parental strand without end trimming, allowing up to 1 bp NH tail (Figure S10A, S10B and Figure 7G step 3). In support of this, previous biochemical analysis showed that Polθ can tolerate one unpaired base at the 3’ ends for DNA synthesis *in vitro*^60^. We further demonstrated that while 1-bp NH tail could be tolerated with reduced efficiency, a 2-bp NH tail completely blocked DNA synthesis (Figure S9B).

### Fork-MMEJ is used more frequently at seDSBs on the leading strands induced by Cas9n

To examine whether fork-MMEJ acts differently on broken leading versus lagging strands, we inserted the EGFP-MMEJ reporter at *Igh* locus in mES cells. In non-B cells, including mES cells, origins are silenced in the ∼400 kb temporal transition region (TTR) at the *Igh* locus, resulting in unidirectional replication by a single fork^61^, which was confirmed by our high-resolution Repli-seq analysis in mES cells^62^ (Figure S11A). We inserted the EGFP-MMEJ reporter in both orientations at *Igh* TTR (*Igh*-MMEJ-For and *Igh*-MMEJ-Rev, Figure 4A), without influencing replication direction in that region (Figure S11B). As replication is unidirectional at *Igh* TTR, we can distinguish each of the four types of seDSBs after g2/Cas9^D10A^ and g2/Cas9^H840A^ cleavage (Figure 4B). Notably, expression of g2/Cas9^WT^ induced similar levels of cMMEJ in both *Igh*-MMEJ-For and *Igh*-MMEJ-Rev reporters, comparable to that in the *ROSA*-MMEJ reporter (Figure 4C left), suggesting that replication direction relative to the MMEJ inserted orientation does not influence cMMEJ at deDSBs. However, fork-MMEJ was more efficiently induced at Type II seDSBs on leading strands (*Igh*-MMEJ-Rev, g2/Cas9^D10A^, lead-seDSBs) compared with Type I seDSBs/deDSBs on lagging strands (*Igh*-MMEJ-For, g2/Cas9^D10A^, lag-seDSBs/deDSBs) (Figure 4C right). As expected, fork-MMEJ was inefficient at Type III seDSBs and IV seDSBs/deDSBs (*ROSA*-MMEJ, *Igh*-MMEJ-For and *Igh*-MMEJ-Rev, g2/Cas9^H840A^), which contain long NH tails at the 3’ ends on the parental strands (Figure 4C right and 4B). We also used another gRNA, g6, which cleaves in the middle of the right MH, and obtained similar results that only Type II lead seDSBs (*Igh*-MMEJ-For, g6/Cas9^D10A^) can be repaired by fork-MMEJ (Figure 4D and S12). These results suggest that fork-MMEJ is more frequently utilized at seDSBs on broken leading strands (lead-seDSBs) compared to broken lagging stands (lag-seDSBs/deDSBs) after Cas9n cleavage.

**Figure 4.**
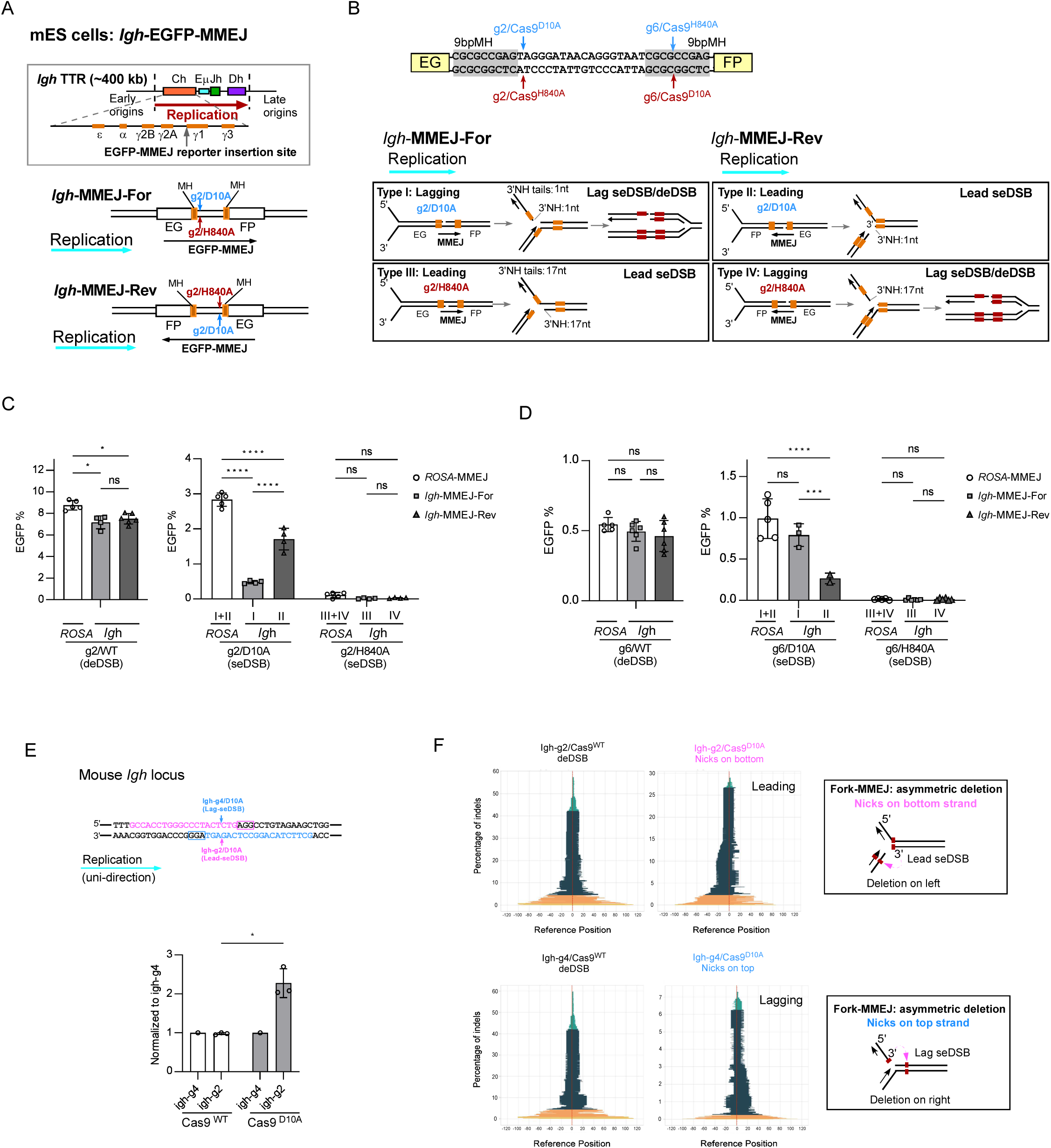
MMEJ is preferentially used on the leading strands to repair seDSBs induced by Cas9n. (**A**) Schematic drawing of the EGFP-MMEJ reporter inserted in the *Igh* TTR locus in mES cells in two different orientations (forward: *Igh*-MMEJ-For and reversed: *Igh*-MMEJ-Rev). (**B**) The four types of seDSB generated after cleavage by gRNA2 along with Cas9^D10A^ or Cas9^H840A^ are illustrated. Unidirectional replication at the *Igh* locus in reference to the orientation of the MMEJ reporter is indicated. (**C**, **D**) MMEJ was assayed in mES (*Igh*-MMEJ-For) and (*Igh*-MMEJ-Rev) cells and mES (EGFP-MMEJ-ROSA) cells after expressing gRNA2 (**C**) or gRNA6 (**D**) along with Cas9^WT^, Cas9^D10A^ or Cas9^H840A^. (**E**) The *Igh* locus in mES cells was cleaved using gRNA Igh-g2 and Igh-g4 with Cas9^WT^ and Cas9^D10A^, followed by deep sequencing analysis of the cleavage sites. The normalized frequency of indels with MMEJ features (≥4 bp deletion and ≥1 bp MH) was calculated by setting the frequency of Igh-g4/Cas9^WT^ and Igh-g4/Cas9^D10A^ to 1, respectively. (**F**) The percentage of repair events with indels (y-axis) corresponding to the indicated deletion sizes (x-axis) is plotted after cleavage of the *Igh* locus in mES cells using Igh-g2 and Igh-g4 with Cas9^WT^ and Cas9^D10A^, with deletion sizes color-coded as described in Figure 1E. Schematic illustrations of the expected deletion patterns of indels are shown on the right.

In *Xenopus* nuclear extracts, Cas9n-induced lag seDSBs contain ∼70 bp 3’ ssDNA overhangs while lead seDSBs carry blunt ends or short 5′ overhangs of up to 3 bp^38^. We speculate that pre-existing 3’ ssDNA overhangs on the broken lagging strands (Figure S1D) may preferentially channel seDSB repair to BIR, thereby reducing the use of fork-MMEJ (Figure S18). In addition, as replication at *Igh* TTR is unidirectional without converging forks^61^, efficient fork-MMEJ detected at Type II seDSBs support the model that fork-MMEJ can directly access to seDSBs to repair broken forks preferentially on the leading strands without involving fork convergence (Figure S1C, fork-MMEJ).

We also analyzed indel formation directly at the *Igh* TTR locus in mES cells, using a pair of gRNAs, Igh-g2/Cas9^D10A^ and Igh-g4/Cas9^D10A^, which cleave the bottom (leading strand) and top (lagging strand) strands, respectively (Figure 4E, top). While the frequency of indel formation with MMEJ features (≥ 1bp MH and ≥ 4 bp deletion) at deDSBs was comparable following cleavage with Igh-g2/Cas9^WT^ and Igh-g4/Cas9^WT^, the frequency at lead-seDSBs induced by Igh-g2/Cas9^D10A^ was significantly higher than that at lag-seDSBs/deDSBs generated by Igh-g4/Cas9^D10A^ (Figure 4E, bottom). In addition, we analyzed genomic loci in U2OS cells where replication proceeds unidirectionally (Figure S13A), using pairs of gRNAs/Cas9^D10A^ to cleave either the leading or lagging strands (Figure S13B left), and the frequency of indels with MMEJ features (≥ 1bp MH and ≥ 4 bp deletion) at those loci was also markedly higher on the leading strands than on the lagging strands (Figure S13B right). These findings are consistent with our MMEJ reporter results (Figure 4C and 4D, Type II), indicating that fork-MMEJ operates preferentially at lead-seDSBs.

Moreover, while cleavage at the *Igh* locus in mES cells with Igh-g2/Cas9^WT^ and Igh-g4/Cas9^WT^ (deDSBs) generated symmetric deletions (Figure 4F left), expressions of Igh-g2/Cas9^D10A^ and Igh-g4/Cas9^D10A^ resulted in asymmetric deletions (Figure 4F middle) as observed at the *LBR* locus in U2OS cells (Figure 1E). As expected, Cas9n-induced indels on the leading strands at the *Igh* locus were reduced when replication was inhibited by aphidicolin (Figure S5D). Collectively, these data are in line with our working model that the 3’ end of the broken parental strand searches for internal MH on the other end and directly primes DNA synthesis without end trimming (Figure S10B).

### Fork-MMEJ functions together with BIR to repair seDSBs

MMEJ is error prone^18,19^, but one advantage of MMEJ over HR/BIR is that MMEJ could act fast; MMEJ only needs short (cMMEJ)^41^ or no (fork-MMEJ) end resection (this study), while HR/BIR requires long end resection, which is time consuming. To determine the contribution of MMEJ and BIR to repair broken forks, we established an EGFP-MMEJ/mCherry-BIR reporter, simultaneously monitoring MMEJ and BIR. We inserted the BIR donor cassette EG-T2A-mCherry upstream of the EGFP-MMEJ cassette (Figure 5A top). When MMEJ is used, green cells are produced. To initiate BIR, the EG in the EGFP-MMEJ cassette invades the EG homology in the EG-T2A-mCherry BIR donor cassette on its sister chromatid, followed by replicating 2 kb through mCherry and reaching the FP homology in the EGFP-MMEJ cassette to complete BIR/LTGC. As a result, the CMV promoter is placed in front of the EG-T2A-mCherry cassette, producing red cells (Figure 5A bottom, BIR/LTGC).

**Figure 5.**
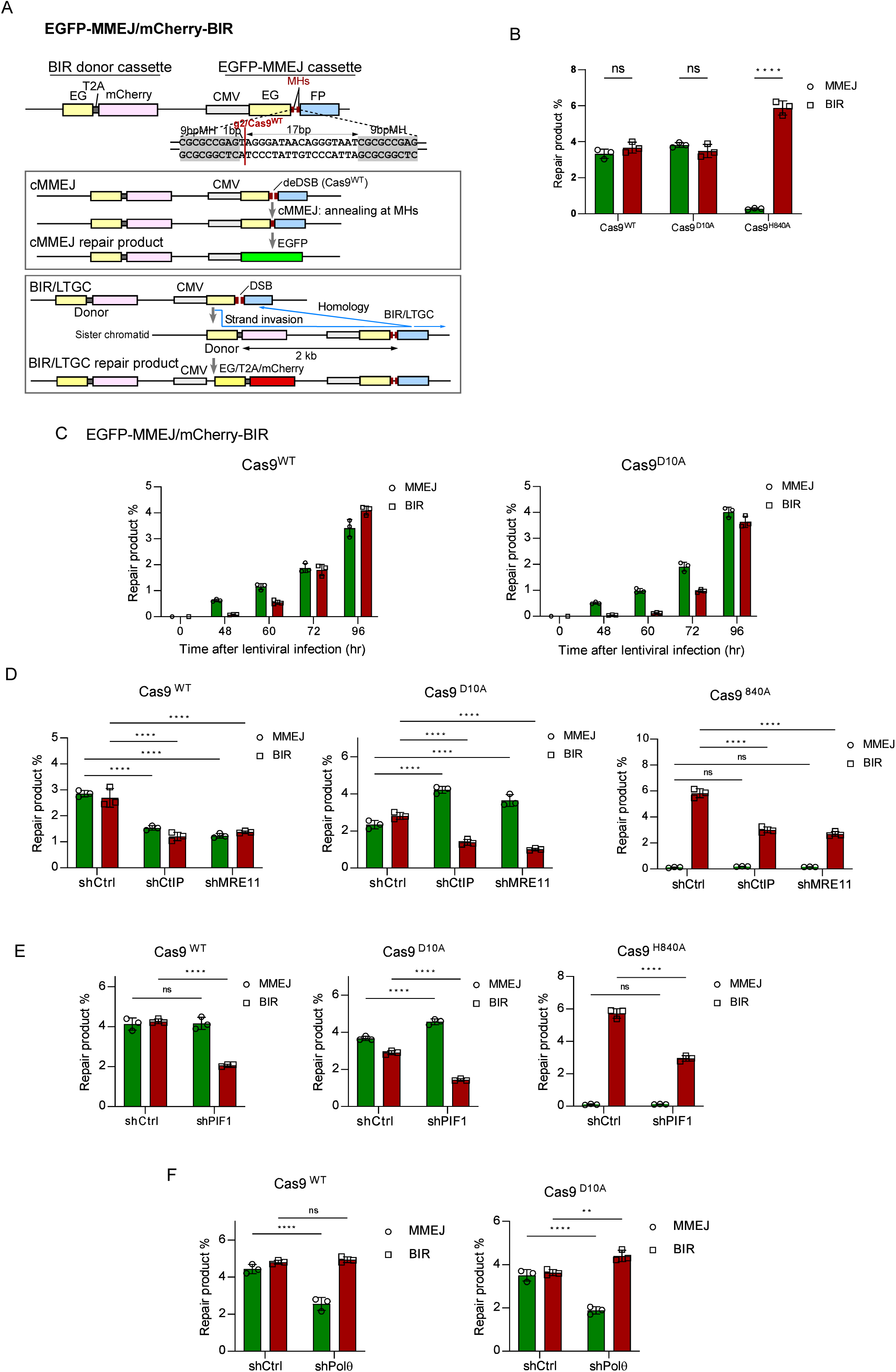
Fork-MMEJ functions together with BIR to repair seDSBs on broken forks. (**A**) Schematic drawings of the EGFP-MMEJ/mCherry-BIR reporter and the repair products by cMMEJ and BIR after Cas9^WT^ cleavage. The gRNA2 cleavage site is indicated. (**B**) MMEJ or BIR was scored in U2OS (EGFP-MMEJ/mCherry-BIR) cells by FACS to determine EGFP or mCherry positive cells after cleavage by gRNA2 with Cas9^WT^ and Cas9^D10A^. (**C**) Time course experiments were performed in U2OS (EGFP-MMEJ/mCherry-BIR) cells to analyze MMEJ and BIR after cleavage by gRNA2 with Cas9^WT^ and Cas9^D10A^. (**D**-**F**) MMEJ and BIR was assayed in U2OS (EGFP-MMEJ/mCherry-BIR) cells expressing shRNAs for CtIP (**D**), MRE11 (**D**), PIF1(**E**) or Polθ (**F**) after expressing gRNA2 with Cas9 and Cas9n.

We inserted the EGFP-MMEJ/mCherry-BIR reporter into the *AAVS1* locus in U2OS cells. cMMEJ/fork-MMEJ (green) and BIR (red) were used concurrently after g2/Cas9^WT^ and g2/Cas9^D10A^ cleavage (Figure 5B). However, g2/Cas9^H840A^ induced substantial BIR, but not fork-MMEJ (Figure 5B), indicating that g2/Cas9^H840A^ effectively generated nicks but long NH tails blocked fork-MMEJ as described (Figure 3A: g2/Cas9^H840A^ and Figure 3B: Type III/IV with 17 bp tail). Time-course experiments showed that cMMEJ (g2/Cas9^WT^) and fork-MMEJ (g2/Cas9^D10A^) were initiated prior to BIR, with substantial BIR observed at later time points (Figure 5C), supporting the quick launch of MMEJ over BIR at both deDSBs and seDSBs.

Consistent with the role of MRE11 and CtIP in suppression of fork-MMEJ (Figure 2C), depletion of MRE11 or CtIP impaired BIR but led to a significant increase in fork-MMEJ after g2/Cas9^D10A^ cleavage in U2OS (MMEJ/BIR) cells, whereas both cMMEJ and BIR were decreased at deDSBs after g2/Cas9^WT^ cleavage (Figure 5D and S14A). We propose that while fork-MMEJ can be launched prior to end resection to repair seDSBs on broken forks, the onset of end resection acts to suppress fork-MMEJ and promotes BIR (Figure 7G left and S18).

To assess whether fork-MMEJ/cMMEJ and BIR can complement the loss of one another, we depleted PIF1 and Polθ by shRNAs in U2OS (MMEJ/BIR) cells. Inactivation of PIF1 resulted in a defect in BIR after cleavage by both g2/Cas9^WT^ and g2/Cas9^D10A^, but with almost no effect on cMMEJ (g2/Cas9^WT^) and a small increase in fork-MMEJ (g2/Cas9^D10A^) (Figure 5E and S14B left). Conversely, depletion of Polθ resulted in a strong defect in cMMEJ and fork-MMEJ, but a small (g2/Cas9^D10A^) or no (g2/Cas9^WT^) increase in BIR (Figure 5F and S14B right). These data suggest that despite being used simultaneously, MMEJ and BIR are not completely interchangeable. Consistent with the importance of both fork-MMEJ and BIR in repairing seDSBs at broken forks, we observed that while depletion of Polθ or PIF1 alone caused HU sensitivity (Figure 6A) and impaired replication restart efficiency and speed (Figure 6B)^23,63^, simultaneous inactivation of both led to more severe defects (Figure 6A, 6B, S14C and S14D).

**Figure 6.**
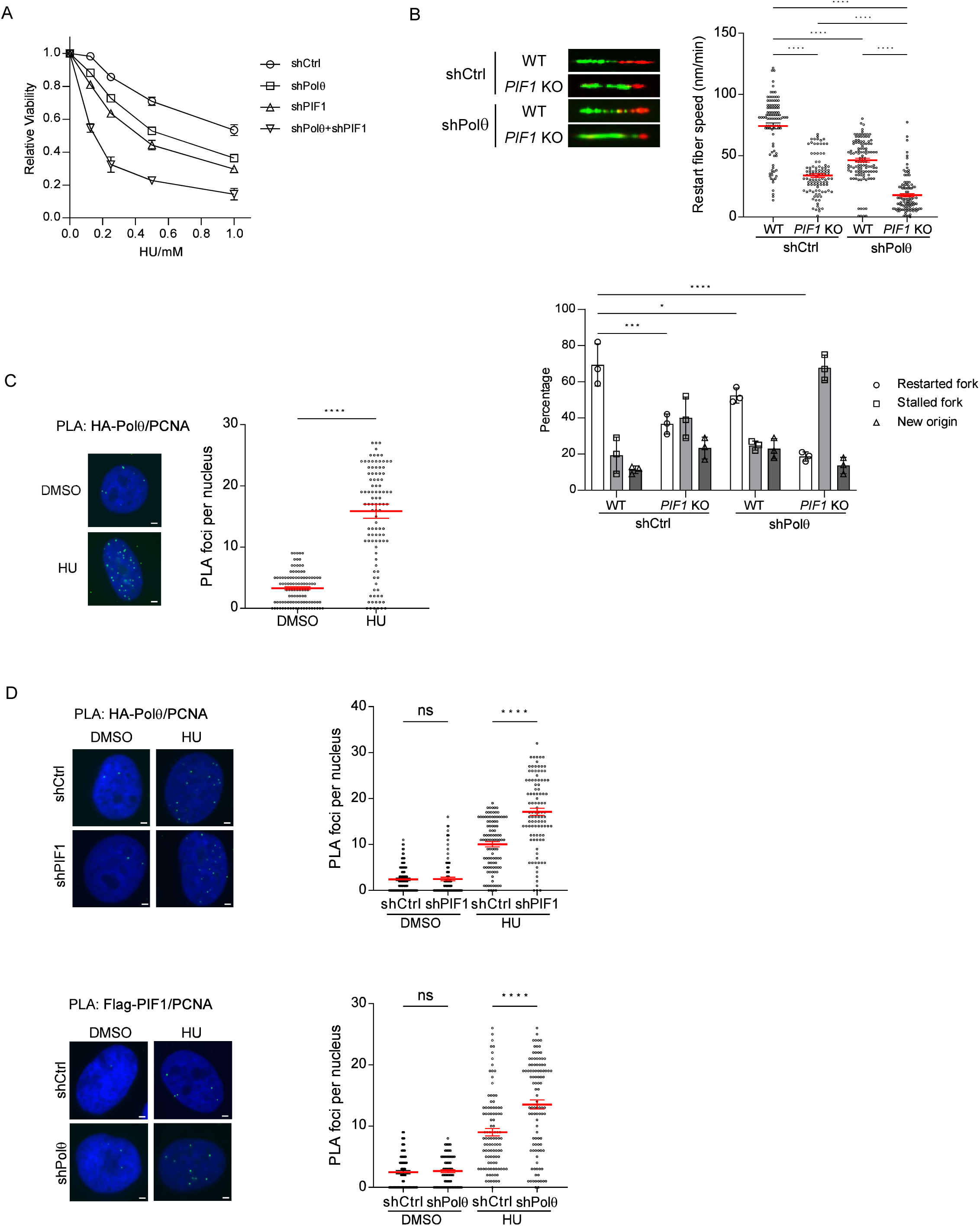
Both MMEJ and BIR are important for replication restart. (**A**) U2OS cells expressing shRNAs for Polθ, PIF1 or both were treated with HU (72 hours), and cell viability was determined. (**B**) DNA fiber analysis was performed in WT or *PIF1*-KO U2OS cells expressing Polθ shRNA or shCtrl. Examples of restarted forks are shown at the top left. The speed of restarted forks (top right) and the percentage of restarted and stalled forks, and new origin firing (bottom) are shown. (**C**) PLA of HA-Polθ and PCNA was performed in U2OS cells with or without treatment of HU (2 mM, 24 hours). (**D**) PLA was performed to show colocalization of HA-Polθ and PCNA with or without depletion of endogenous PIF1 (left) and colocalization of Flag-PIF1 and PCNA with or without depletion of endogenous Polθ (right) before and after HU (2 mM, 24 hours) treatment. Scale bar, 2 μm.

**Figure 7.**
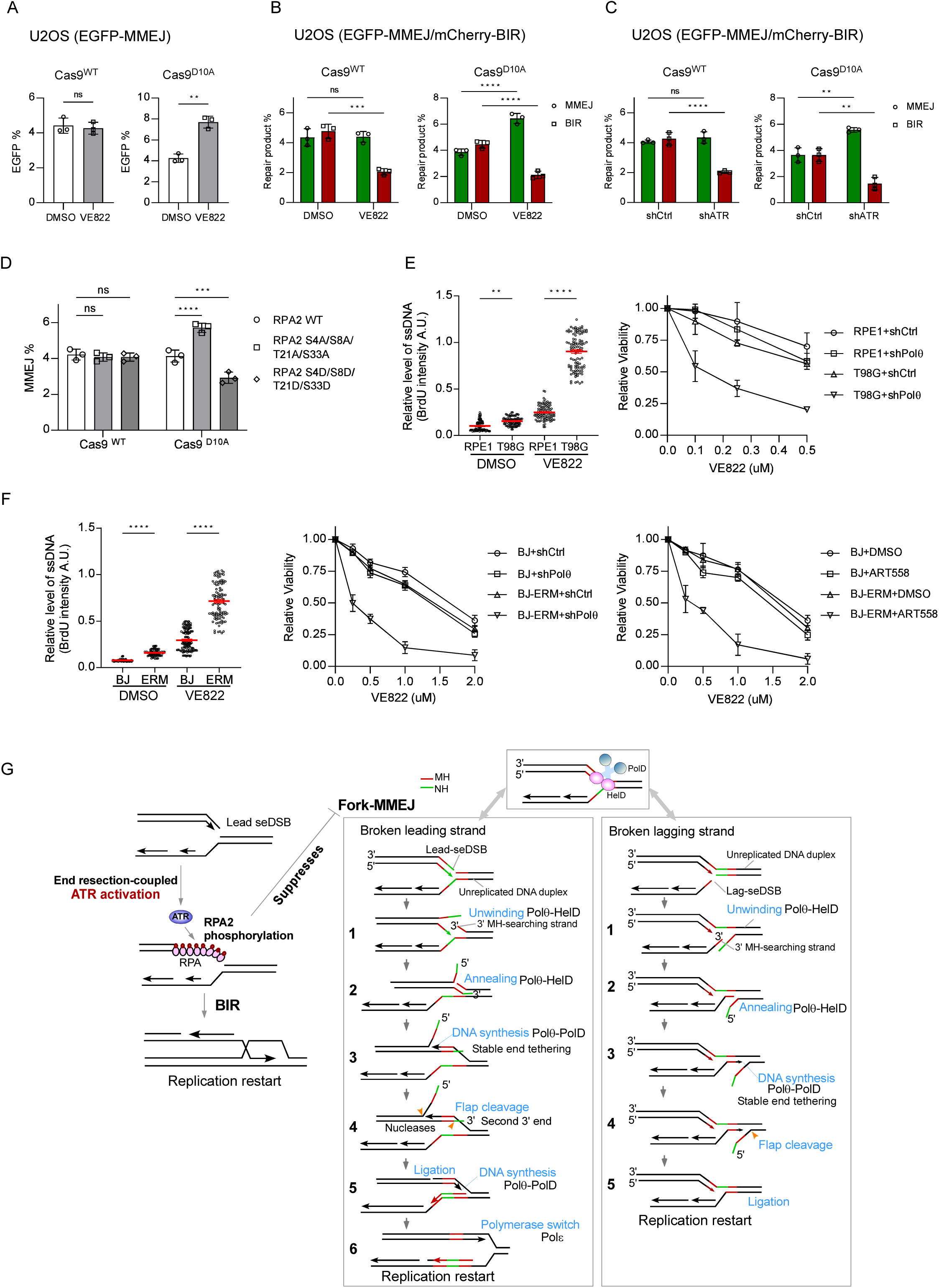
ATR, activated by end resection, specifically inhibits fork-MMEJ. (**A**) U2OS (EGFP-MMEJ) cells treated with or without VE822 (0.5 μM), or (**B**) U2OS (EGFP-MMEJ/mCherry-BIR) cells treated with or without VE822 (0.5 μM), or (**C**) expressing ATR shRNA or shCtrl, were assayed for MMEJ after expressing gRNA2 with Cas9^WT^ or Cas9^D10A^. (**D**) U2OS (EGFP-MMEJ) cells, expressing RPA2-WT or 4A and 4D mutants with endogenous RPA2 silenced by shRNA, were assayed for MMEJ after expressing gRNA2 with Cas9^WT^ and Cas9^D10A^. (**E**) RPE1 or T98G cells treated with VE822 (10 μM, 2 hours) or DMSO were assayed for the level of ssDNA by BrdU staining (left). Cell viability was determined for T98G or RPE1 cells expressing Polθ shRNA with treatment of VE822 (72 hours, right). (**F**) Normal BJ cells and transformed BJ cells expression E1A, RAS-V12 and MDM2 (ERM) were assayed for the level of ssDNA by BrdU staining (left), and cell viability after expressing Polθ shRNA or vector (middle) or in the presence of DMSO or ART558 (5 μM, right) after treatment with VE822 (72 hours). (**G**) Proposed models for the control of switching from fork-MMEJ to BIR by ATR activation induced upon end resection (left) and for the fork-MMEJ mechanism at seDSBs upon fork breakage (see main text for details).

PCNA has been shown to interact with both Polθ and PIF1 in a manner dependent on PCNA ubiquitination^64,65^. *In situ* proximity ligation assay (PLA) revealed that the interaction between Polθ and PCNA increased after HU treatment (Figure 6C), which was further enhanced after PIF1 depletion (Figure 6D top and Figure S14B left). Conversely, Polθ depletion also led to an increase in PIF1 interaction with PCNA after HU treatment (Figure 6D bottom and Figure S14B right). Furthermore, the recruitment of Polθ and PIF1 to DSB sites marked by γH2AX after HU treatment was dependent on PCNA (Figure S15A and S15B) and PCNA ubiquitination (Figure S15C). These data suggest that both Polθ and PIF1 are recruited to DSB sites by PCNA on broken forks, enabling the use of both fork-MMEJ and BIR.

### ATR inhibits fork-MMEJ and promotes the shift of fork-MMEJ to BIR in conjunction with end resection

Although it is well established that ATR is important for HR^66^, its role in MMEJ has remained unclear. Strikingly, we observed that inhibition of ATR activity with the ATR inhibitor (ATRi) VE822 led to a drastic increase in fork-MMEJ, but not cMMEJ, in U2OS (EGFP-MMEJ-AAVS1) cells (Figure 7A). Moreover, inhibition of ATR activity by ATRi EV822 or ATR shRNA in U2OS (MMEJ/BIR) cells revealed that, while BIR was reduced at both deDSBs (Cas9^WT^) and seDSBs (Cas9^D10A^), cMMEJ (Cas9^WT^) was unaffected, whereas fork-MMEJ (Cas9^D10A^) was significantly increased (Figure 7B, 7C and S16A). This suggests that ATR is required for BIR at both deDSBs and seDSBs, but specifically suppresses fork-MMEJ upon fork breakage, without significantly affecting cMMEJ at deDSBs.

We showed that RPA suppresses cMMEJ, but is required for fork-MMEJ (Figure 2B). Notably, RPA2 is progressively phosphorylated by ATR following end resection at replication-associated, but not replication-independent DSBs^67^. We mutated damage-induced phosphorylation sites S4 and S8 [DNAPK dependent^68–70^ and ATR partially dependent^67^] along with T21 and S33 [ATR dependent^67^] on PRA2. Strikingly, in the RPA2-S4A/S8A/T21A/S33A mutant (RPA2-4A) with endogenous RPA2 depleted, fork-MMEJ but not cMMEJ was significantly increased (Figure 7D and S16B). Conversely, the phosphomimetic RPA2-S4D/S8D/T21D/S33D mutant (RPA2-4D) was defective in fork-MMEJ but not cMMEJ (Figure 7D and S16B). We propose that as end resection progresses at seDSBs on broken forks, ATR is activated to phosphorylate RPA2, which impairs RPA activity required for fork-MMEJ, thereby suppressing fork-MMEJ (Figure 7G left).

To test whether MRE11-mediated end resection, ATR activation, and RPA phosphorylation act in the same pathway to suppress fork-MMEJ, we treated cells expressing MRE11 shRNA or RPA2-4A (+RPA2 shRNA) with ATRi. ATR inhibition markedly increased fork-MMEJ in control cells, but caused almost no further increase in MRE11-depleted or RPA2-4A mutant cells (Figure S17A). We previously showed that ∼200 bp ssDNA overhangs caused RPA2 phosphorylation in human nuclear extracts^71^. We further showed that ∼50 bp ssDNA overhangs were sufficient to trigger ATR-mediated RPA2 phosphorylation, with stronger induction by longer overhangs (Figure S17B). Collectively, these data suggest that ∼50 bp ssDNA overhangs can induce substantial ATR-mediated RPA2 phosphorylation, which in turn suppresses fork-MMEJ.

### Inhibition of Polθ and ATR shows a synergistic effect to eradicate cancer cells under replication stress

ATR is important for fork protection^5,72^, thereby preventing fork breakage and DSB formation^73^. Consistent with ATR deficiency reducing BIR and increasing fork-MMEJ, co-inhibition of ATR and Polθ synergistically killed T98G tumor cells but, interestingly, not the untransformed RPE1 cell line (Figure 7E right and S16C). As described, ATR inhibition leads to ssDNA accumulation, but only excessive ssDNA levels result in DSB formation^73^. We speculate that in normal cells, where replication stress is low, ATRi-induced ssDNA and DSBs remain minimal and tolerable, but in tumor cells with high replication stress, ATRi drives ssDNA beyond the threshold, generating DSBs^73^ (Figure S19). Consistently, T98G, but not RPE1 cells, exhibited high replication stress, as indicated by ssDNA accumulation after short ATRi exposure^73^ (Figure 7E left). Similarly, isogenic tumor cells derived from primary fibroblast BJ cells by expressing E1A, H-RAS-V12 (RAS) and MDM2 oncogenes (ERM), but not normal BJ cells, displayed high replication stress (Figure 7F, left). In contrast to normal BJ cells, while ATRi alone (VE822) caused a small increase of cell death in BJ (ERM) cells, combining ATRi with Polθ depletion or inhibition induced synergistic lethality (Figure 7F, middle and right, and S16D). These results suggest that dual inhibition of Polθ and ATR selectively eradicates replication-stressed tumor cells while sparing normal cells.

## DISCUSSION

Polθ plays an important role for cells to cope with replication stress^14,22–25^. Our study demonstrated that fork-MMEJ, which depends on Polθ activity, can directly operate at broken forks to repair seDSBs upon fork breakage. This provides new insights into the mechanistic understanding of MMEJ and highlights its significance in replication restart for maintaining genome stability.

### The mechanism underlying fork-MMEJ differs from cMMEJ at deDSBs

Fork-MMEJ on broken forks differs from cMMEJ at deDSBs in that it does not require end resection but relies on RPA. We propose that instead of end resection, DNA unwinding, facilitated by Polθ helicase activity, exposes MH for fork-MMEJ (Figure 7G, right). *In vitro*, Polθ-HelD exhibits DNA unwinding activity, which is enhanced by RPA^59^. We further showed that ssDNA can anneal to MH unwound by Polθ-HelD from a duplex DNA, which then prime DNA synthesis by Polθ-PolD. Since we did not detect a direct interaction between RPA and Polθ-HelD or Polθ-PolD (Figure S8D), the requirement for RPA in fork-MMEJ may stem from its role in stabilizing ssDNA during Polθ-HelD-mediated DNA unwinding. As Polθ-HelD helicase activity is considered low^59^, it is also possible that Polθ-HelD acts with other helicases to promote DNA unwinding, with their activities requiring RPA. We examined the role of the MCM replicative helicase and found that it is not involved in fork-MMEJ (Figure S6F).

Another notable difference is that cMMEJ tolerates long NH tails (flaps) on both ends of a DSB, but fork-MMEJ is strongly suppressed by ≥2 bp NH tails outside of the MH at the 3’ broken end on the parental strand (Type III and IV: Figure 3B, blue circle of the 3’ end; Figure S10A), whereas the other 5’ end of the parental strand permits long NH tails (flaps) (Type I and II: Figure 3B, blue rectangle of the 5’ end; Figure S10A). Upon fork breakage, the two broken ends on the parental strands need to religate for restoring fork integrity (Figure S10B). This requires the 3’ end of the broken parental strand to anneal with the 3’ daughter strand of the other end (lead-seDSBs) or the other unbroken parental strand (lag-seDSBs) using an MH sequence (Figure S10B). We propose that the MH at the 3’ end of the broken parental strand is used to search for MH internal of the seDSB (designated as 3’ searching end, Figure S10B). This is supported by the observations of asymmetric deletion patterns observed after Cas9n cleavage (Figure 1E and 4F), which is different from the symmetric deletion pattern of cMMEJ at deDSBs resulting from bidirectional end resection after Cas9^WT^ cleavage (Figure 1G top). The tolerance for a 1-bp, but not a 2-bp, NH tail at the 3’ searching end for fork-MMEJ is consistent with the biochemical activity of Polθ, showing that only a single unpaired base, but not more than one, at the 3’ end permits Polθ-PolD-mediated DNA synthesis *in vitro*^60^ (Figure S9B).

We speculate that relying on DNA unwinding rather than end resection, and directly using the 3’ end of the broken parental strand for MH searching and DNA synthesis without end trimming, are driven by the urgent need for fork-MMEJ to rapidly repair broken forks. At deDSBs, KU-dependent c-NHEJ acts immediately, while c-MMEJ requires end resection and is engaged later. However, at seDSBs on broken forks, where c-NHEJ is unavailable, fork-MMEJ serves as the immediate repair pathway. Because end resection is time-consuming, directly using the 3’ end of broken ends for MH searching, together with DNA unwinding to expose MH, enables faster repair. In addition, since MH annealing is transient and prone to dissociation, initiating bridging DNA synthesis directly from the 3′ end without end trimming can quickly stabilize broken forks for efficient repair (Figure 7G, step 3). It has been shown that in c-MMEJ, Polδ uses its exonuclease activity to remove NH tails^74^. Because this process requires sequential handoffs—Polθ aligns MHs, then a switch to Polδ for NH removal with each round to remove one nucleotide, followed by a switch back to Polθ for DNA synthesis—skipping trimming could allow faster engagement of fork-MMEJ.

Using our reporter, we score a fixed event with designed MHs to produce green cells; in this context, 3′ NH tails on the broken parental strands inhibit fork-MMEJ (Figure S10C left). However, at genomic sites, the 3’ ends of broken parental strands can directly search for MHs within a relatively large region of the other broken ends (Figure S10C right). Given that there is more than a 93% chance of finding a ≥ 3bp MH within 15 bp of any given pair of DNA ends, a high chance is predicted for a fixed end to find ≥1-3 bp on the other side of the break to yield productive fork-MMEJ ^75^.

### A proposed working model of fork-MMEJ for seDSB repair

We propose a working model as illustrated in Figure 7G. Upon fork breakage, the two Polθ-HelDs tether the broken forks through dimerization^76,77^, with one binding to the seDSB and the other to the unreplicated DNA duplex with a 3’ broken end (Figure 7G right top). Both DNA duplex of the unreplicated DNA and the other seDSB end (lead-seDSB; lag-seDSB if ssDNA overhang is not sufficient) need to be unwound to expose MHs (Figure 7G, step 1 and S10B).

Subsequently, the 3’ end on the broken parental strands (3’ MH-searching strand) is directly used to search for an MH internal of the other end and anneals with it (Figure 7G, step 2 and S10B), followed by DNA synthesis from the 3’ end without end trimming (Figure 7G, step 3). After bridging DNA synthesis from one end, the broken forks are stably tethered together. In contrast to the 3’ MH-searching end that is directly used for priming DNA synthesis, the second 3’ end at lead-seDSB requires 3’-flap nucleases, such as XPF/ERCC1^78^, to remove the 3’ flap prior to initiating DNA synthesis (Figure 7G, step 4). In addition, strand displacement DNA synthesis may occur, generating 5’ flaps which need to be removed by nucleases at both lead-seDSB and lag-deDSB (Figure 7G, step 4). 5’-flap nucleases such as SLX1^78,79^, DNA2^80^, FEN1^81^ and FAN1^82^ could possibly contribute to 5’-flap removal, which requires further investigation. Polθ is not a processive polymerase, and a switch from Polθ to Polδ^74^ may occur during strand displacement DNA synthesis in fork-MMEJ. Since Polε disassociates with CMG from lead-seDSBs^36,38^, Polε needs to be reloaded after completion of fork-MMEJ to restore normal replication forks and promote replication restart (Figure 7G, step 6). Finally, fork-MMEJ is completed by end ligation on the parental strand and replication forks are restored (Figure 7G right, step 5).

### Fork-MMEJ directly acts on seDSBs, alongside BIR, to repair replication-associated fork breakage

While BIR is a well-established mechanism for repairing seDSBs, we found that fork-MMEJ can also directly repair seDSBs immediately upon fork breakage, before BIR engages for replication restart. Both Polθ and PIF1 are recruited by ubiquitinated PCNA upon replication stress, but pathway choice is not determined merely by their recruitment, rather by the extent of 3′ end resection, which governs the balance between fork-MMEJ and BIR. Using the MMEJ/BIR dual reporter, we showed that inhibiting end resection by depleting MRE11 or CtIP strongly stimulates fork-MMEJ while suppressing BIR, whereas depleting PIF1, which functions downstream of resection, blocks BIR but only modestly increases fork-MMEJ. Thus, while end resection controls the choice between fork-MMEJ and BIR, once a pathway is committed, the two cannot fully compensate for each other when one is deficient. Consistently, simultaneous inactivation of both pathways causes stronger defects in fork restart and hypersensitivity to replication stress than loss of either alone.

Recent studies have shown that in addition to BIR, HR is also employed at broken forks^83,84^, following the formation of deDSBs after fork convergence (both lead- and lag-seDSBs) and replication bypass (lag-seDSBs) (Figure S18). To assess HR involvement, we established a competition reporter to monitor fork-MMEJ and HR at broken forks (Figure S19A). While both cMMEJ and fork-MMEJ occurred prior to HR (Figure S19B), HR increased at later time points, with its usage eventually slightly exceeding cMMEJ (Cas9^WT^), but remaining lower than fork-MMEJ (Cas9^D10A^). This differs from BIR after Cas9^D10A^ cleavage, which increased to levels comparable to fork-MMEJ at later time points (Figure 5C). These findings support the model that seDSBs at broken forks are first repaired by fork-MMEJ, followed by BIR, with HR engaged when seDSBs are converted to deDSBs. They also indicate that BIR is used more frequently than HR at broken forks, consistent with our previous observations^63^.

Using the EGFP-MMEJ reporter, we observed that fork-MMEJ is more preferentially used on leading strands after Cas9n cleavage. This is consistent with the observation that following Cas9n cleavage, indels accumulate more frequently on leading strands at genomic sites exhibiting unidirectional replication over ∼100 kb (Figure S20). We anticipate that the preferential use of fork-MMEJ on broken leading strands is largely due to the presence of 3′ ssDNA overhangs (∼70 bp) at lag-seDSBs^38^, which could directly channel the repair toward BIR, whereas lead-seDSBs are blunt-ended or with short 5′ overhangs and thus require resection to initiate BIR (Figure S18). In addition, rapid conversion of lag-seDSBs to deDSBs by CMG bypass may more quickly activate HR on broken lagging strands than on leading strands, further contributing to the reduced use of fork-MMEJ on lagging strands.

MMEJ has been considered a backup pathway for DSB repair when HR is defective^13,23^. BIR also relies on key HR factors such as RAD51 and BRCA1^63^. End-seq analysis revealed that in RAD51-deficient cells, seDSBs on both broken leading and lagging strands are converted to deDSBs, which are also hyper end-resected^36^. As the resulting long ssDNA overhangs expose MHs, in HR/BIR-deficient cells, it is likely that cMMEJ, rather than fork-MMEJ, is involved in the repair of broken forks. Furthermore, we anticipate that Polθ-mediated fork-MMEJ is distinct from previously described Polθ activities involved in filling lagging-strand gaps when Okazaki fragment elongation is stalled^24^ or promoting microhomology-mediated gap skipping at post-replicative ssDNA gaps in BRCA-deficient cells^25^. In those contexts, Polθ-driven gap-filling occurs at stalled forks before DSB formation and is likely associated with Polθ DNA synthesis activity^85^. In contrast, fork-MMEJ, as defined in our study, functions after DSB formation at broken forks and is linked to the end-joining activity of Polθ in MMEJ.

### ATR promotes the transition from fork-MMEJ to BIR in conjunction with the extent of end resection

Fork-MMEJ provides rapid repair of seDSBs on broken forks, but need to be tightly regulated given its error-prone nature. Our study suggests that ATR controls the use of fork-MMEJ by phosphorylating RPA2, linking end resection to a switch from fork-MMEJ to BIR. At replication-associated DSBs, end resection-generated RPA-ssDNA drives ATR-mediated RPA2 phosphorylation, coupling resection extent to RPA2 modification^67^. Our model is that progressive resection at replication-associated DSBs activates ATR to phosphorylate RPA2, which in turn specifically suppresses fork-MMEJ without affecting cMMEJ at deDSBs (Figure 7G left). It remains possible that other ATR targets, in addition to RPA2, may also contribute to fork-MMEJ suppression.

The pronounced effect of ATR inhibition in enhancing fork-MMEJ while suppressing BIR, implies that ATR inhibition would result in a strong reliance on fork-MMEJ for repairing seDSBs. Indeed, simultaneous inhibition of ATR and Polθ synergistically kills cancer cells by blocking both repair pathways. In normal cells, replication stress is low, and even with ATR inhibition, ssDNA levels remain tolerable with minimal DSB formation (Figure S21), resulting in a low demand for fork repair. In contrast, cancer cells experience high replication stress^73^, and ATR inhibition therefore causes extensive ssDNA accumulation, fork breakage, and seDSBs, which require fork-MMEJ and BIR for repair. Consequently, combined ATR and Polθ inhibition effectively eradicates cancer cells with low toxicity to normal cells, providing a new strategy for targeted cancer therapy.

### Limitations of the study

Our EGFP-MMEJ reporter was designed to monitor MMEJ using 9 bp MH (Figure 1A). It can also be adapted to assess MMEJ using MH of 8 bp (g5), 7 bp (g4) (Figure 3C), and 5 bp (g6) (Figure S12) by designing gRNAs that cleave within the MHs. However, due to limited gRNA target sites in this reporter, we could not use the same EGFP-MMEJ to score MMEJ with MHs shorter than 5 bp. Instead, we generated separate reporters to assess repair efficiency when different MH lengths were used (Figure S9A). In addition, building on the mechanistic understanding from the studies using the EGFP-MMEJ reporter, we verified obtained general principles through direct deep sequencing analysis of indels generated at genomic loci following Cas9 or Cas9n cleavage. For indel analysis at each cleavage site, we need to analyze at least 50K reads for accurate measurements. This limits our study to examining sites one by one, rather than using whole-genome deep sequencing to assess many sites simultaneously.

## RESOURCE AVAILABILITY

### Lead contact

Further information and requests for resources and reagents should be directed to and will be fulfilled by the lead contact, Xiaohua Wu.

### Materials availability

Materials generated in this study are available through lead contact upon request with a standard Material Transfer Agreement.

### Data and code availability

- The raw data of site-specific PCR deep sequencing, nascent strand sequencing of U2OS cells, and Repli-seq of mouse Embryonic stem cells used in this manuscript are available in the NIH Sequence Read Archive (SRA) as BioProject: PRJNA1235770.
- Original images of uncropped gels are deposited at Mendeley Data: 10.17632/d79jttdxyc.1 and are publicly available as of the date of publication.
- This paper does not report original code.
- Any additional information required to reanalyze the data reported in this paper is available from the lead contact upon request.

## ACKNOWLEDGMENTS

Sumo3 PolQM1 (Addgene plasmid # 78462) and pCDH-EF1-FHC-POLQ (Addgene plasmid # 64875) are gifts from Dr. Richard D. Wood (The University of Texas MD, Anderson Cancer Center). pSpCas9(BB)-2A-Puro (PX459) V2.0 (Addgene plasmid # 62988) and lentiCRISPR v2 (Addgene plasmid # 52961) are gifts from Dr. Feng Zhang (Massachusetts Institute of Technology). pLKO.1-blast (Addgene plasmid # 26655) is a gift from Dr. Keith Mostov (University of California, San Francisco). pCDH-CMV (Addgene plasmid # 72265) is a gift from Dr. Kazuhiro Oka (Baylor College of Medicine). pCW-Cas9 (Addgene plasmid # 50661) is a gift from Dr. Eric Lander & Dr. David Sabatini (Massachusetts Institute of Technology). Tet-On 3×Flag Polθ is a gift from Dr. Dale A. Ramsden (University of North Carolina, Chapel Hill). We thank Dr. Peiqing Sun (Wake Forest University) for the BJ and BJ (EMR) cells. This work was supported by research grants from NIH CA187052, CA197995, GM080677 and CA294646 (X.W.), and in part by National Natural Science Foundation of China (32270579 to S.L.), the intramural program of the Center for Cancer Research, National Cancer Institute (ZIA BC010411 to M.I.A.) and NIH CA270335 (D.M.G).

## AUTHOR CONTRIBUTIONS

Conceptualization, S.L. and X.W.; Methodology, S.L., Y.Z., Y.L., S.B.S., Y.S., T.N., A.R. and T.S.; Software, Y.S., A.R., S.L. and T.S.; Validation, S.L., Y.Z., Y.L., S.B.S., Y.S., A.R. and T.S.; Formal Analysis, S.L., Y.Z., Y.L., S.B.S., Y.S. and T.S.; Investigation, S.L., Y.Z., Y.L., S.B.S., Y.S., T.N., Z.W., C.C., A.R., T.B., S.L., T.S. and J.H.S.; Writing – original draft, X.W. and S.L.; Writing – review & editing, S.L., D.M.G., M.I.A. and X.W.; Visualization, S.L., Y.Z., Y.L., S.B.S., Y.S.,T.N., A.R. and T.S.; Supervision, S.L., H.W., D.M.G., M.I.A. and X.W.; Project administration, S.L. and X.W.; Funding acquisition, X.W., S.L. and M.I.A.

### DECLARATION OF INTERESTS

The authors declare no competing interests.

## SATR METHODS

### KEY RESOURCES TABLE

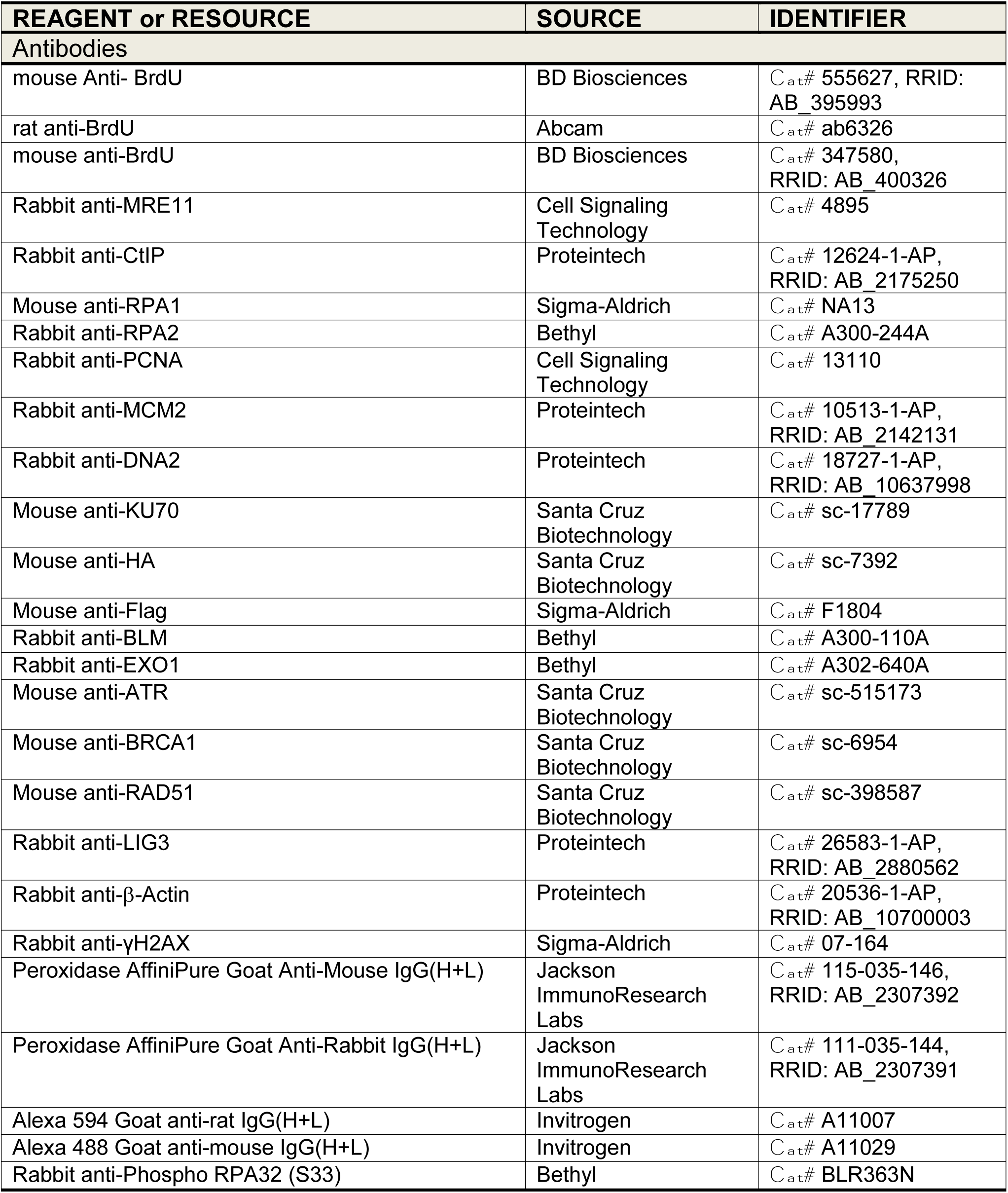
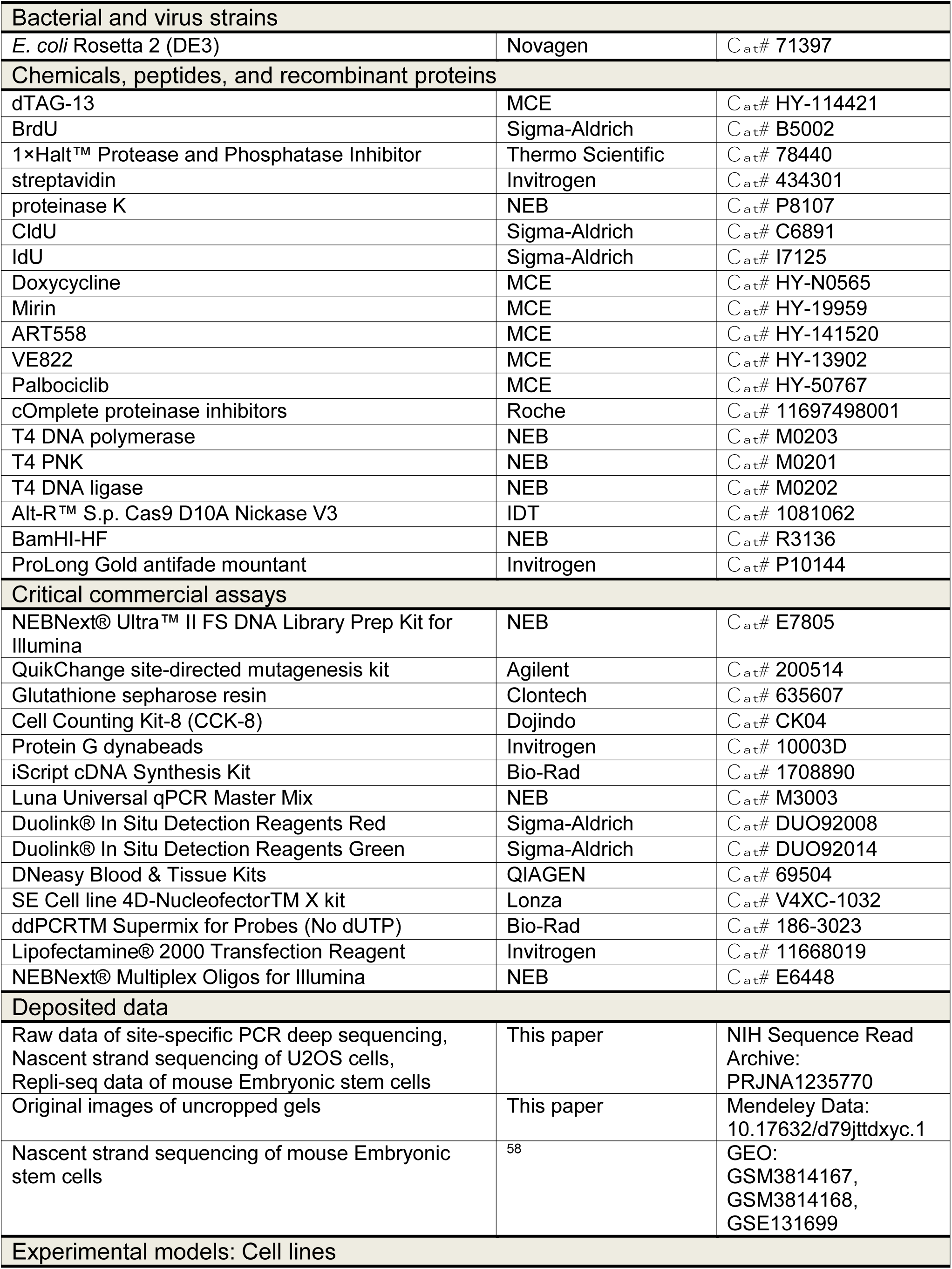
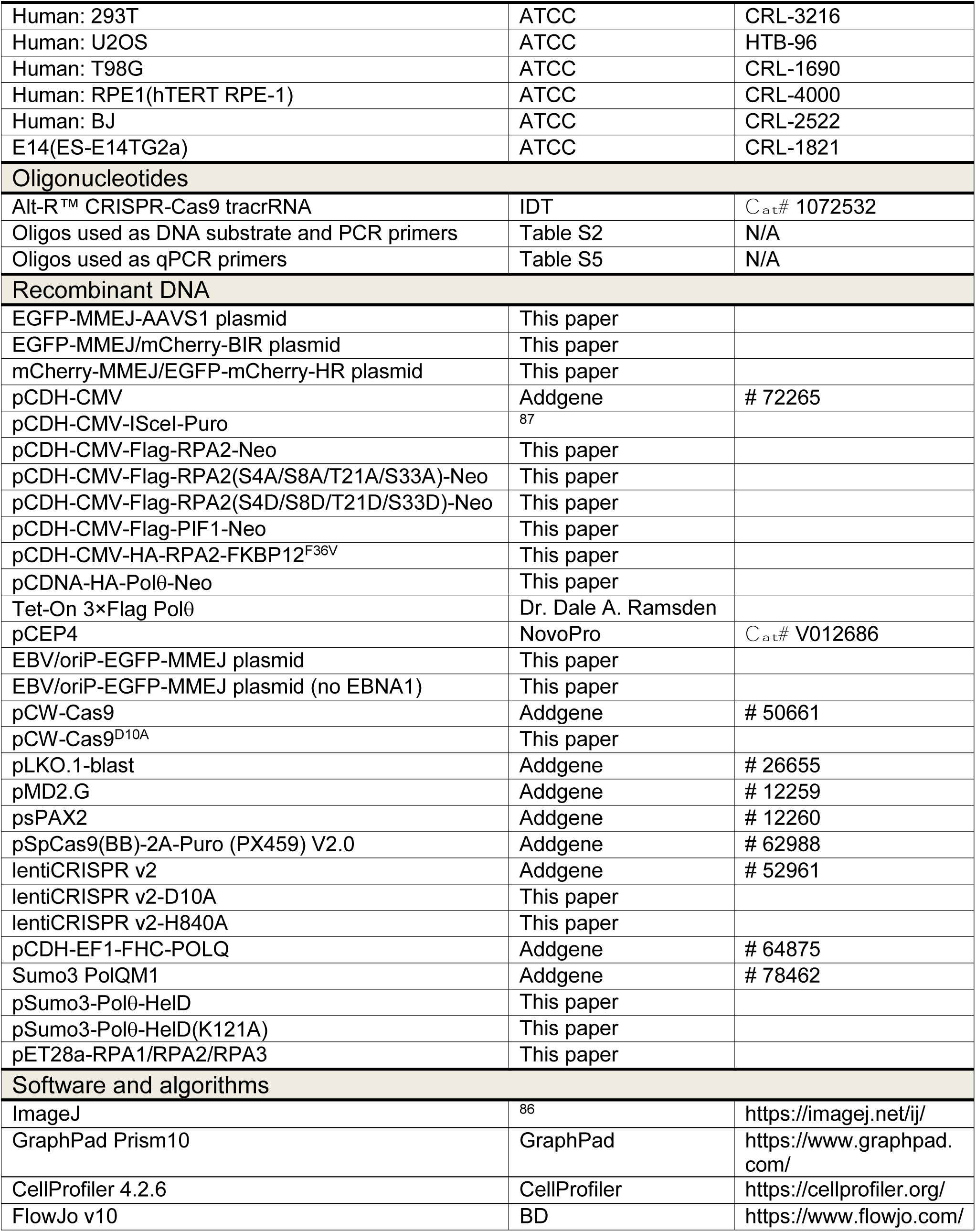

### EXPERIMENTAL MODEL AND STUDY PARTICIPANT DETAILS

#### Cell lines and cell culture

U2OS (human osteosarcoma), HEK293T (human embryonic kidney), and T98G (human glioblastoma) and RPE1 (human retina) were obtained from ATCC and tested negative for mycoplasma contamination. BJ (ERM) tumor cells were generated by infecting BJ cells with retroviruses encoding oncogenes E1A, H-RAS-V12 (RAS) and MDM2 (ERM).

U2OS, HEK293T, BJ cells are grown in Dulbecco’s modified Eagle’s medium (DMEM, Gibco) supplemented with 10% fetal bovine serum (FBS, GeminiBio) and 1% penicillin-streptomycin containing glutamine (Gibco). T98G cells are grown in Minimum essential medium (MEM, Gibco) supplemented with 10% fetal bovine serum (FBS, GeminiBio), 1% non-essential amino acids (NEAA, Gibco) and 1% penicillin-streptomycin containing glutamine (Gibco).

RPE1 cells are grown in DMEM/F-12 Medium supplemented with 10% fetal bovine serum (FBS, GeminiBio) and 1% non-essential amino acids (NEAA, Gibco). E14 mouse embryonic stem (mES) cells are grown in Glasgow modification of Eagle’s medium (GMEM, Sigma) supplemented with 15% ES-qualified FBS (Gibco), 1% non-essential amino acids (Gibco), 2 mM L-glutamine (Life Technologies), 0.1 mM β-mercaptoethanol (Sigma), 1mM sodium pyruvate (Gibco) and leukemia inhibitory factor (LIF) (2000U/ml, Amsbio). All cells are cultured at 37°C in 5% CO_2_ atmosphere.

### METHOD DETAILS

#### Plasmid construction and generation of the reporter cell lines

The CMV-EGFP-MMEJ reporter was generated by placing a disrupted full-length EGFP under the control of a CMV promoter, with insertion of an 18-bp sequence containing in-frame stop codons and a 9-bp microhomology sequence between E112 and V113 as described^87^.

The EGFP-MMEJ/mCherry-BIR reporter was generated by placing the sequence encoding the 1-108 amino acid of EGFP with an in-frame fusion of T2A, followed by a full-length mCherry, in front of the EGFP-MMEJ cassette as shown in Figure 5A.

To generate the mCherry-MMEJ/EGFP-mCherry-HR reporter, the MMEJ cassette was constructed by inserting into the EGFP open reading frame a gRNA cleavage site followed by three stop codons, flanked by a duplication of 9-bp microhomology sequence that replaced the chromophore of EGFP (T65/Y66/G67)^88^. An in-frame T2A-mCherry sequence was fused to the 3’ of the MMEJ cassette. A truncated EGFP fragment (1-167 aa) was placed in front of the MMEJ cassette to serve as the HR donor cassette, as shown in Figure S19A.

To integrate the reporters into the *AAVS1* locus in U2OS cells, the reporter cassettes were subcloned into an *AAVS1*-targeting vector containing hygromycin-resistance gene. Integration was achieved using CRISPR/Cas9 with the *AAVS1* gRNA (Table S1) and confirmed by PCR analysis. The EGFP-MMEJ reporter was also randomly integrated into the U2OS genome and clones with single integration was confirmed by ddPCR.

To generate the E14 mES (EGFP-MMEJ-ROSA) reporter cell line, the CAG-EGFP-MMEJ reporter, generated by replacing the CMV promoter of the CMV-EGFP-MMEJ reporter with the CAG promoter, was targeted to the *ROSA* locus by CRISPR/Cas9 (ROSA gRNA in Table S1). To generate *Igh*-MMEJ-For or *Igh*-MMEJ-Rev reporters, the CAG-EGFP-MMEJ cassette was cloned into a vector containing 1 kb homology arms to the *Igh* locus on both sides of the reporters in either forward or reverse directions. *Igh*-MMEJ-For or *Igh*-MMEJ-Rev reporters were targeted to the *Igh* locus in E14 mES cells by CRISPR/Cas9 (*Igh* gRNA in Table S1). The EBV/oriP-EGFP-MMEJ reporter was generated by inserting the CMV-EGFP-MMEJ reporter into the pCEP4 vector or the pCEP4-derived vector without the EBNA1 coding sequence.

#### *In vitro* assays of Polθ helicase and polymerase activities

Coding sequence of the human Polθ helicase domain (1-894 aa) was amplified from pCDH-EF1-FHC-POLQ (Addgene, #64875) and inserted into the Sumo3 PolQM1 plasmid (Addgene, #78462, encoding Polθ-PolD) to replace the Polθ polymerase domain. The helicase-dead mutation K121A was introduced into Polθ helicase domain by QuikChange site-directed mutagenesis (Agilent, 200514). The Polθ polymerase domain (Polθ-PolD), helicase domain (Polθ-HelD) and RPA1/RPA2/RPA3 complex were expressed in *E. coli* Rosetta 2 (DE3) and purified as described^89,90^.

Double-stranded DNA (dsDNA) substrates, with or without single-stranded DNA (ssDNA) overhangs, were generated by annealing of ssDNA oligos. Briefly, the indicated amount of ssDNA oligos were mixed in NEBuffer 2, heated in a water bath at 95°C for 5 minutes, and then allowed to cool slowly to room temperature.

DNA unwinding activity of Polθ-HelD was determined as described^91^. 5 nM ^32^P-5’-radiolabeled pre-annealed DNA substrate was mixed with Polθ-HelD WT or K121A mutant (20 nM) in reaction buffer (25 mM Hepes-NaOH, pH 7.0, 2 mM DTT, 0.01% NP-40, 40 mM KCl, 5% glycerol, 1 mM MgCl_2_), followed by the addition of ATP (2 mM), cold ssDNA trap (200 nM) and RPA complex (20 nM) as indicated for 15 minutes at 37°C.

To assay for DNA unwinding and strand displacement by Polθ-HelD, 10 nM ^32^P-5’-radiolabeled ssDNA was mixed with 5 nM pre-annealed dsDNA substrate, Polθ-HelD WT or K121A mutant (20 nM), and ATP (2 mM), and RPA complex (20 nM) were added to the reactions as indicated, followed by incubation at 37°C for 15 minutes. The reaction was stopped by addition of 10 mg/ml proteinase K (NEB, P8107), 10 mM EDTA, and 0.08% SDS at final concentration, and resolved on 12% non-denaturing polyacrylamide gels and visualized by phosphor imaging.

To access the activities of Polθ-HelD and Polθ-PolD in strand displacement DNA synthesis, the strand displacement reaction by Polθ-HelD was supplemented with 20 nM Polθ-PolD and 0.1 mM dNTPs and incubated for another 20 minutes at 37°C. The reactions were terminated by adding an equal volume of denaturing stop buffer (90% formamide, 50 mM EDTA), resolved on 10% denaturing polyacrylamide gels (8 M urea), and visualized by phosphor imaging.

To examine the ability of Polθ-PolD to extend DNA synthesis from 3’ unpaired bases, 100 nM ^32^P-5’-radiolabeled DNA substrates were incubated with Polθ-PolD (20 nM) and 0.5 mM dATP in reaction buffer (20 mM Tris-HCl, pH 8.0, 100 mM NaCl, 5 mM MgCl_2_) for indicated time at 37°C. The reactions were terminated by adding equal volume of denaturing stop buffer (90% formamide, 50 mM EDTA) and resolved on 15% denaturing polyacrylamide gels (8 M urea) and visualized by phosphor imaging.

#### GST pull-down experiments

GST-fused Polθ helicase (Polθ-HelD) and polymerase (Polθ-PolD) domains were expressed in *E. coli* Rosetta 2 (DE3). Bacteria pellets were lysed in NETN buffer (20 mM Tris-HCl, pH 8.0, 150 mM NaCl, 1 mM EDTA, 0.5% NP-40) and incubated with Glutathione sepharose resin (Clontech, 635607) for 3 hours at 4°C, followed by three washes with NETN buffer. Purified RPA1/RPA2/RPA3 (RPA) complex was incubated with the glutathione resin bound with GST-fused Polθ-HelD and Polθ-PolD or GST in NETN buffer for 3 hours at 4°C. After extensive washing, GST pull-down samples were subjected to 15% SDS-PAGE, followed by Western blot analysis using anti-RPA1 and anti-RPA2 antibodies.

#### Generation of *POLQ* knock-out and mutants by CRISPR

A pair of *POLQ* KO gRNAs (POLQ KO gRNA1 and gRNA2 in Table S1) targeting mouse *POLQ* exon 1 were subcloned into a CRISPR vector expressing pSpCas9 and mCherry marker. Transfections were carried out in mES cells containing the EGFP-MMEJ reporter with

Lipofectamine 2000 (Invitrogen) according to the manufacturer’s recommendations. 72 hours after transfection, mCherry-positive cells were sorted by flow cytometry and single cells were placed into individual wells of a 96-well plate pre-coated with 0.1% gelatin. *POLQ* knock-out clones were screened by PCR amplification using a pair of primers (Mouse *POLQ* KO screening-F and Mouse *POLQ* KO screening-R in Table S2) and were confirmed by Sanger sequencing.

*POLQ* helicase and polymerase mutants in E14 cells were generated following the described strategy^89^. For making the Polθ helicase mutant (K120G) in E14 cells harboring the EGFP-MMEJ reporter, a pair of gRNAs (K120G gRNA1 and gRNA2 in Table S1) were subcloned into the pX459 V2.0-Cas9^D10A^ vector, and the resulting gRNA plasmids were co-transfected together with a donor cassette using Lipofectamine 2000 (Invitrogen). The donor plasmid contains 690 bp and 565 bp homology arms on the upstream and downstream of the mutation K120G. Similarly, for Polθ polymerase mutant (D2494P/E2495R) in E14 cells, a pair of gRNAs (D2494P/E2495R gRNA1 and gRNA2 in Table S1) were subcloned in pX459 V2.0-Cas9^D10A^ and were co-transfected together with a donor template containing 700 bp and 544 bp homology arms on the upstream and downstream of the mutation D2494P/E2495R. 24 hours after transfection, cells were selected for puromycin (1 μg/ml, 2 days) to enrich transfected cells, and then maintained in regular media for 3-4 days. Single clones were expanded and screened for the knock-in mutations by genomic DNA PCR sequencing (Primers for K120G: K120G screening-F and K120G screening-R; Primers for D2494P/E2495R: D2494P/E2495R screening-F and D2494P/E2495R screening-R in Table S2).

#### EGFP-MMEJ, EGFP-MMEJ/mCherry-BIR and mCherry-MMEJ/EGFP-mCherry-HR reporter assays

For protein knockdown in U2OS cells carrying the EGFP-MMEJ, EGFP-MMEJ/mCherry-BIR or mCherry-MMEJ/EGFP-mCherry-HR reporter, cells were infected with concentrated lentiviruses expressing shRNAs for indicated genes with the empty vector as a control. 24 hours after infection, cells were replaced with fresh media and subjected to blasticidin selection (10

μg/ml, 2 days). To perform reporter assays, U2OS cells were infected by lentiviruses, encoding the indicated gRNAs cloned in the lentiCRISPRv2 vector or its derived vectors, expressing either Cas9^D10A^ or Cas9^H840A^, followed by puromycin selection (2 μg/ml, 2 days). EGFP or mCherry-positive events were analyzed four days after viral infection of Cas9 or Cas9n using a BD Accuri C6 flow cytometer. The gRNA cleavage sites in the EGFP-MMEJ, EGFP-MMEJ/mCherry-BIR, and mCherry-MMEJ/EGFP-mCherry-HR reporters are identical; all reporters use the same gRNAs for the repair assays.

To perform reporter assays in E14 cells, MMEJ gRNA2 and gRNA6 in Table S1, targeting the EGFP-MMEJ reporter, were cloned in the pSpCas9(BB)-2A-Puro (PX459) V2.0 vector (Addgene plasmid # 62988) or its derived vectors that express Cas9^D10A^ or Cas9^H840A^, followed by transfection to E14 cells carrying the EGFP-MMEJ reporter at the *ROSA* or *Igh* locus.

Transfection was carried out with a total of 3×10^5^ cells per well in a 24-well plate with Lipofectamine 2000 (Invitrogen) according to the manufacturer’s instructions (1 μg plasmid DNA per well). 4 hours later, transfection mixes were replaced with fresh medium, and after another 24 hours, 1 μg/ml puromycin was added for 48 hours to eliminate untransfected cells. Cells were analyzed for EGFP-positive events by flow cytometry 4 days post infection.

#### Droplet digital PCR (ddPCR) assay for quantification of double-strand breaks

To determine the percentage of cells with DSB formation at the site of gRNA cleavage, ddPCR was performed as described^92^. Two amplicons were designed, of which, one includes the gRNA/Cas9 target site, and the other is a proximal uncut site as a control. A dual-quenched probe was designed for each amplicon, which was labeled using FAM or HEX at the 5’ of the probe for target site and control site. 50 ng genomic DNA was used as template in a 20 μl reaction containing 10 μl 2×ddPCR Supermix for Probes (No dUTP) (Bio-Rad, 186-3023), 900 nM of each primers and 250 nM probes for both target and control sites. BamHI-HF (NEB, R3136) was added to the reaction at a ratio of 1:100 for better separation of signals. Droplets were generated using QX200 Droplet generator (Bio-Rad). The amplification was performed on a C1000 Touch Thermal Cycler with the following conditions: 95°C for 10 min, 40 cycles of (94°C for 30 s, 60°C for 30 s, 72°C for 1 min), 98°C for 10 min, 12°C hold. Droplets were scanned using the QX200 Droplet Digital PCR system (Bio-Rad). Droplets in each fluorescent channel (FAM/HEX) were plotted and distinguished in clusters using a global threshold to bin droplets to positive and negative labels. The DSB frequency was calculated as [target-, control+]/([target-, control+] + [target+, control+]).

#### Light-inducible CRISPR

Caged gRNA for EGFP-MMEJ reporter was designed as described^48^, and synthesized by Bio-Synthesis Inc. 200 pmol caged gRNA was mixed with 200 pmol Alt-R™ CRISPR-Cas9 tracrRNA (IDT, 1072532), heated to 95°C for 5 minutes, and allowed to cool to room temperature for 5 minutes. 3 μl of 10 μg/μl Alt-R™ S.p. Cas9 D10A Nickase V3 (IDT, 1081062) was added to the annealed mixture of caged gRNA and tracrRNA, and was incubated at room temperature for 20 minutes to form RNP complex. 1×10^7^ U2OS (EGFP-MMEJ) cells were resuspended in 90 μl SE solution and supplied with 20 μl Supplement solution from SE Cell line 4D-NucleofectorTM X kit (Lonza, V4XC-1032). Electroporation was performed according to the manufacturer’s instructions on the 4D-Nucleofector Core Unit (Lonza) with code DN-100. Cells were plated in complete DMEM and incubated at 37°C for 16 hours for recovery.

#### Synchronization in G1 phase and cell cycle analysis of U2OS cells

To synchronize U2OS cells in G1 phase, Palbociclib was added to 70% confluent cells at a final concentration of 0.25 μM for 20 hours. Cells remaining in G1 arrest or released from G1 by replacing with fresh medium were trypsinized, and vigorously resuspended in 1 ml PBS, followed the dropwise addition of 4 ml cold ethanol. After fixation for 1 hour at 4°C, cells were stained with 10 µg/mL Propidium Iodide in the presence of 0.2 mg/mL RNase A. Cell cycle profile was analyzed by flow cytometry and the percentage of cells in different cell cycle phases was analyzed by FlowJo.

#### Analysis of Cas9^WT^, Cas9^D10A^ or Cas9^H840A^-induced indels at genomic loci by deep sequencing

gRNAs targeting human *LBR* locus (LBRg1 and LBRg2 in Table S1) and mouse *Igh* locus (Igh-g2 in Table S1) were cloned into pSpCas9(BB)-2A-Puro (PX459) V2.0 vector (Addgene plasmid # 62988) and LentiCRISPR v2 vector (Addgene plasmid # 52961) or their derived vectors expressing Cas9^D10A^ or Cas9^H840A^. 72 hours after transfection of plasmids encoding gRNAs with Cas9^WT^, Cas9^D10A^ or Cas9^H840A^, genomic DNA was extracted using DNeasy Blood & Tissue Kits (QIAGEN, 69504). Libraries of repair products were generated by PCR amplification using primers for *LBR* and *Igh* listed in Table S2, producing ∼250 bp DNA fragments surrounding the gRNA target sites. A second round of PCR was performed with P5 and P7 primers containing i5 and i7 index sequences, and subjected to GENEWIZ for paired-end 150 bp deep sequencing on illumina NovaSeq platform. Quality control of raw reads was performed using FastQC. Paired-end reads were merged using FLASH, followed by trimming and demultiplexing with Cutadapt. High-quality reads were aligned to the reference genome using BBMap. Repair patterns were analyzed based on CIGAR values using Python. Data visualization and statistical analyses were conducted in R and GraphPad Prism.

#### DSB end capture

To directly demonstrate DSBs formation at the EGFP-MMEJ reporter after Cas9 and Cas9n cleavage, we adapted DSBcapture^47^ for DSB end capture at a specific genomic locus (Fig. S4A). 24 hours after infection by lentiviruses expressing gRNA and Cas9^WT^ or Cas9^D10A^, 1×10^7^ cells were fixed with 2% formaldehyde at room temperature for 30 minutes, followed by addition of glycine to a final concentration of 125 mM to quench formaldehyde. Cells were collected, washed with ice cold PBS and lysed in lysis buffer (10 mM Tris-HCl, pH 8.0, 10 mM NaCl, 1 mM EDTA, 1 mM EGTA, 0.2% NP-40, 1 mM DTT, cOmplete proteinase inhibitors) by gently rotating at 4°C for 90 minutes. After centrifugation at 1200 rpm for 5 minutes, nuclei were resuspended in nucleus break buffer (10 mM Tris-HCl, pH 8.0, 10 mM NaCl, 1 mM EDTA, 1 mM EGTA, 0.3% SDS, 1 mM DTT) and incubated at 37°C for 45 minutes. Nuclei pellets were then resuspended in NEBuffer 2 (10 mM Tris-HCl, pH8.0, 50 mM NaCl, 10 mM MgCl_2_, 1 mM DTT) with 0.1% Triton X-100, and proteinase K was added to a final concentration of 100 µg/mL. After incubation at 37°C for 8 minutes, an equal volume of NEBuffer 2 containing 0.1% Triton X-100 and 2 mM PMSF was added to inactivate proteinase K. Nuclei were then washed twice by resuspending in NEBuffer 2 with 0.1% Triton X-100 followed by centrifugation at 1200g for 10 minutes. Nuclei were washed with Blunting Buffer (100 mM Tris-HCl, pH 7.5, 50 mM NaCl, 10 mM MgCl_2_, 0.025 % Triton X-100, 5 mM DTT) with 100 µg/mL BSA and were blunt-ended by the addition of 2 µL T4 DNA polymerase (NEB, M0203), 0.5 µL T4 PNK (NEB, M0201) and 4 µL 2.5mM dNTPs in a 50 µl volume, followed by incubation at 25°C for 45 minutes. Nuclei were then washed three times with NEBuffer 2 with 0.1% Triton X-100, resuspended in NEBuffer 2 and A-tailed using 3 µL Klenow Fragment 3’-5’ exo- (NEB, M0212L) and 1 µL 5 mM dATP in a final volume of 50 µL at 37°C for 45 minutes. Following A-tailing, nuclei were washed three times with NEBuffer 2 with 0.1 % Triton X-100, once with T4 DNA ligase reaction buffer containing 0.1% Triton X-100 at 4°C and resuspended in T4 DNA ligase reaction buffer. Annealed adaptor containing UMI (UMI-For and UMI-Rev in Table S2), was then ligated to DNA ends using T4 DNA ligase (NEB, M0202M) for 15 hours at 16°C in a final reaction volume of 50 µL. For genomic DNA extraction, the ligation mixture was treated with Proteinase K (200 µg/ml) for 30 minutes at 55°C, followed by incubation at 65°C for 30 minutes, and DNA was precipitated using isopropanol. The pelleted DNA was resuspended in 50 µL of water. The DSB capture library was then amplified by nested PCR, followed by an additional PCR to add illumina i5 and i7 sequences to the ends of the amplicons. The final PCR products were subjected to paired-end 150 bp deep sequencing on the illumina NovaSeq platform. The PCR primers for DSB end capture after Cas9 and Cas9n cleavage of the EGFP-MMEJ reporter are listed in Table S2.

#### Lentiviral infection for protein expression and shRNA interference

The coding sequences of PCNA, Flag-RPA2, RPA2-FKBP12^F36V^ were subcloned into the pCDH-CMV-EF1-Puro vector (NovoPro, V006738). RPA2 S4A/S8A/T21A/S33A and S4D/S8D/T21D/S33D mutants, as well as the PCNA-K164R mutant, were generated using QuikChange site-directed mutagenesis kit (Agilent, 200514). Short hairpin RNAs (shRNAs) targeting the indicated genes were cloned into the pLKO.1-blast vector (Addgene #26655). shRNA target sequences were listed in Table S4.

For lentivirus production, 293T cells were co-transfected with lentiviral expression plasmids, along with pMD2.G (Addgene, #12259) and psPAX2 (Addgene, #12260) using standard calcium chloride protocol. Lentivirus particles released into the cell culture medium were collected 72 hours after 293T transfection. The cell culture medium was filtered and concentrated using PEG-800 (40% W/V) containing NaCl (1.2 M). After lentiviral infection, cells were selected with blasticidin (10 μg/ml, 2 days) or puromycin (2 μg/ml, 2 days). Protein expression levels of the targeted genes were verified by Western blot and RT-qPCR using primers listed in Table S5.

#### Cell viability assay

Cells were seeded in 96-well plates at a density of 2×10^3^ cells per well and treated with indicated concentration of drugs for 72 hours. Subsequently, 100 μl of cell medium from each well was mixed with 20 μl Cell Counting Kit-8 (Dojindo, CK04) and incubated at 37°C for 2 hours. Relative viability was determined by measuring the emission at 490 nm using 800TS Microplate Reader (BioTek), and normalized to control cells treated with DMSO.

#### Immunoblotting

Cells from a confluent 6 cm plate were lysed in NETN buffer (20 mM Tris-HCl, pH 8.0, 100 mM NaCl, 0.5 mM EDTA, 0.5% NP-40) at 4°C for 30 minutes, followed by centrifugation (12500 rpm for 5 minutes) to remove cell debris. After adding 2×loading buffer (100 mM Tris-HCl, pH 6.8, 4% SDS, 0.2% bromophenol blue, 20% glycerol, 200 mM DTT) to the cell lysates, the samples were heated at 95°C for 5 minutes and separated on 6∼15% SDS-PAGE. Antibodies used in immunoblotting are: MRE11 (Cell Signaling Technology, 4895), CtIP (Proteintech, 12624-1-AP), RPA1 (Sigma-Aldrich, NA13), RPA2 (Bethyl, A300-244A), PCNA (Cell Signaling Technology, 13110), MCM2 (Proteintech, 10513-1-AP), DNA2 (Proteintech, 18727-1-AP), KU70 (Santa Cruz Biotechnology, sc-17789), HA (Santa Cruz Biotechnology, sc-7392), Flag (Sigma-Aldrich, F1804), BLM (Bethyl, A300-110A), EXO1 (Bethyl, A302-640A), ATR (Santa Cruz Biotechnology, sc-515173), BRCA1 (Santa Cruz Biotechnology, sc-6954), RAD51 (Santa Cruz Biotechnology, sc-398587), LIG3 (Proteintech, 26583-1-AP), γH2AX (Upstate, 07-164), Peroxidase AffiniPure Goat Anti-Mouse IgG (H+L) (Jackson ImmunoResearch Labs, 115-035-146), and Peroxidase AffiniPure Goat Anti-Rabbit IgG (H+L) (Jackson ImmunoResearch Labs, 111-035-144).

#### Reverse transcription-quantitative PCR (RT-qPCR)

Total RNA was extracted from cell lines using TRIzol reagent (Invitrogen) according to manufacturer’s instructions. cDNA was synthesized from 1 μg of total RNA as the template by reverse transcription using the iScript cDNA Synthesis Kit (Bio-Rad, 1708890) according to the manufacturer’s protocol. The synthesized cDNA was quantified by qPCR using Luna Universal qPCR Master Mix (NEB, M3003) in a C1000 Thermal Cycler (Bio-Rad) using HPRT as an internal control. Primers used for RT-qPCR are listed in Table S4.

#### DNA fiber assay

Cells were first pulse labeled with 40 µM CldU (Sigma-Aldrich, C6891) for 30 minutes, followed by treatment with 2 mM HU for 2 hours, and then labeled with 200 µM IdU (Sigma-Aldrich, I7125) for 30 minutes. After trypsinization, cells were resuspended in PBS and then labeled and unlabeled cells were mixed in a ratio of 1:6.5 μl of cell suspension was placed on a glass slide, then mixed with 7.5 μl lysis solution (200 mM Tris-HCl, pH 7.5, 50 mM EDTA, 0.5% SDS) for 3 minutes. Tilt the slide to a suitable angle to spread the DNA. The fibers were fixed in a 3:1 (vol/vol) methanol: acetic acid solution. DNA was denatured with HCl and blocked with 5% bovine serum albumin (BSA) in PBS for 1 hour after washing. CldU and IdU detection were performed using rat anti-BrdU (Abcam, ab6326) and mouse anti-BrdU (BD Biosciences, 347580) primary antibodies for 2 hours at 37°C, followed by incubation with Alexa 594 anti-rat (Invitrogen, A11007) and Alexa 488 anti-mouse (Invitrogen, A11029) for 1 hour at room temperature. Coverslips were washed with PBS with 0.1% Tween-20 and mounted with Prolong Gold antifade reagent (Invitrogen, P10144). DNA fibers were imaged with a LSM 780 confocal laser scanning microscope and analyzed using ImageJ software (NIH, USA). At least 100 replication forks were analyzed per experimental condition, with the results representing the mean of three independent experiments.

#### *In situ* Proximity Ligation Assay (PLA)

Proximity ligation assay was performed following the manufacturer’s recommendation (Duolink, Sigma-Aldrich, DUO92101). Cells treated with HU (2 mM) for 24 hours were fixed in 2% paraformaldehyde for 20 minutes. Fixed cells were treated with 0.5% Triton X-100 for 10 minutes for permeabilization. Cells were then blocked with 3% BSA for 30 minutes at 37°C in a humidity chamber, followed by incubation with primary antibodies diluted in Duolink Antibody Diluent overnight at 4°C. Coverslips were fixed onto glass slides using ProLong Gold antifade mountant (Invitrogen, P10144) with DAPI. The PLA signals were visualized as distinct fluorescence spots, and images were captured with an Olympus confocal microscope using a 60X objective. The number of PLA signals was quantified using CellProfiler 4.2.6 software.

#### Chromatin immunoprecipitation (ChIP) assay

The recruitment of Flag-Polθ and γH2AX to the Cas9 cleavage site on the EGFP-MMEJ reporter was performed by ChIP assay. U2OS (EGFP-MMEJ) cells expression Flag-Polθ or RPA2-FKBP12^F36V^ was infected with lentivirus expressing gRNA2/Cas9^WT^ or gRNA2/Cas9^D10A^. 24 hours after removal of the virus, cells were fixed with 1% formaldehyde for 10 minutes at room temperature. Glycine was added to the final concentration of 125 mM for 15 minutes at room temperature. After washing twice with cold PBS, cells were lysed and subjected to sonication. The supernatant was collected and incubated with Protein G dynabeads (Invitrogen, 10003D) which have been pre-loaded with indicated anti-Flag antibody or anti-γH2AX antibody for 4 hours at 4 °C, followed by washing with 1 ml TSE I (20 mM Tris-HCl, pH 8.1, 150 mM NaCl, 2 mM EDTA, 0.1% SDS, 1% Triton X-100), TSE II (20 mM Tris-HCl, pH 8.1, 500 mM NaCl, 2 mM EDTA, 0.1% SDS, 1% Triton X-100), buffer III (10 mM Tris-HCl, pH 8.1, 0.25 M LiCl, 1 mM EDTA, 1% NP-40, 1% deoxycholate), and TE. The protein-DNA complex was eluted from beads by elution buffer (1% SDS, 0.1M NaHCO_3_), and cross-linking was reversed by adding in 4 μl of 5 M NaCl and incubating at 65 °C for 6 hours, followed by proteinase K digestion (NEB, P8107) for 2 hours at 42°C. DNA was recovered by column purification, and analyzed by RT-qPCR using primers MMEJ-ChIP-F and MMEJ-ChIP-R.

#### Immunofluorescence

The procedure for visualizing single-stranded DNA was described previously^93^. In short, cells were cultured in 10 µM BrdU (Sigma-Aldrich, B5002) for 24 hours. After PBS washing, cells were incubated in pre-extraction buffer (10 mM PIPES, pH 6.8, 100 mM NaCl, 300 mM sucrose, 3 mM MgCl_2_, and 0.2% Triton X-100) for 5 minutes, fixed with 3% paraformaldehyde in PBS for 20 minutes. Subsequently, fixed cells were washed with PBS, treated with cold acetone for 30 seconds, washed again with PBS, and blocked in 2% BSA for 1 hour. Then blocked cells were incubated with the primary antibody at 4°C overnight. After washing three times with blocking buffer, the cells were incubated with appropriate secondary antibodies for 1 hour in the dark, then incubated in DAPI solution for 2 minutes. Cells were subjected to immunofluorescence analyses using an Olympus IX-81 fluorescence microscope.

#### Nuclear extract-based assay to analyze RPA2 phosphorylation

Nuclear extracts were prepared from U2OS cells as previously described^71^. Nuclear extracts were incubated with 1 mM Mirin at 37°C for 10 minutes in reaction buffer containing 10 mM Hepes-KOH, pH 7.9, 50 mM KCl, 0.5 mM DTT, 4 µM Okadaic acid, 1 mM ATP, 0.1 mM MgCl_2_, 5 mM creatine phosphate and 1×Halt™ Protease and Phosphatase Inhibitor Cocktail (Thermo Fisher Scientific, 78440). 1 µM 3’-biotinylated ssDNA was annealed to its complementary strand at a ratio of 1:1 to generate dsDNA substrate with 3’ overhang of indicated length. 200 nM annealed substrate was incubated with 1.5-fold streptavidin (Invitrogen, 434301) in reaction buffer at 25°C for 30 minutes. The reaction was initiated by mixing equal volume of DNA substrates with nuclear extracts, and incubated at 37°C for 30 minutes. The reaction was stopped with 2×SDS loading buffer (100 mM Tris-HCl, pH 6.8, 4% SDS, 0.2% bromophenol blue, 20% glycerol, 200 mM DTT) and resolved on 12% SDS-PAGE. RPA2 phosphorylation was determined by immunoblotting using phospho-RPA32 (S33) antibody (Bethyl, BLR363N). DNA sequences are listed in Table S2.

#### Replication timing analysis by Repli-seq

Repli-seq was performed as previously described^94^. Briefly, cells were labeled with 100 µM BrdU (Sigma-Aldrich, B5002) for 2 hours and then fixed in 75% ethanol. Fixed cells were then sorted by FACS into early-S and late-S fractions based on propidium iodide staining of DNA. DNA was then purified from each fraction, sheared and adaptor-ligated with NEBNext Ultra II FS Library DNA Prep Kit for Illumina (NEB, E7805). BrdU-labeled nascent DNA library fragments were then enriched by immunoprecipitation with anti-BrdU antibody (BD, 555627). IP products are then PCR amplified and indexed with NEBNext Multiplex Oligos for Illumina (NEB, E6448). The libraries were sequenced on NovaSeq X Plus for 150 bp paired-end run to obtain at least 10 million clusters per library. The sequencing data were analyzed as previously described^94^.

### QUANTIFICATION AND STATISTICAL ANALYSIS

Statistical analysis was performed using GraphPad Prism 10 and Microsoft Excel. The data are presented as mean values ±SD from at least three independent experiments. Significant differences were determined by unpaired Student’s t-test between the two groups or two-way ANOVA test between multiple groups. The *p* value is labeled as \**p* < 0.05, \*\**p* < 0.01, \*\*\**p* < 0.001, \*\*\*\**p* < 0.0001, and ns (not significant) *p* > 0.05. The statistical significance of the replication timing difference was analyzed using SwitchRT (https://github.com/GilbertLab-dnadave/SwitchRT)

## Notes

### Competing Interest Statement

The authors have declared no competing interest.

