## supplemental for "Microhomology-mediated end joining acts directly on replication forks to repair single-ended double strand breaks"

Figure S1

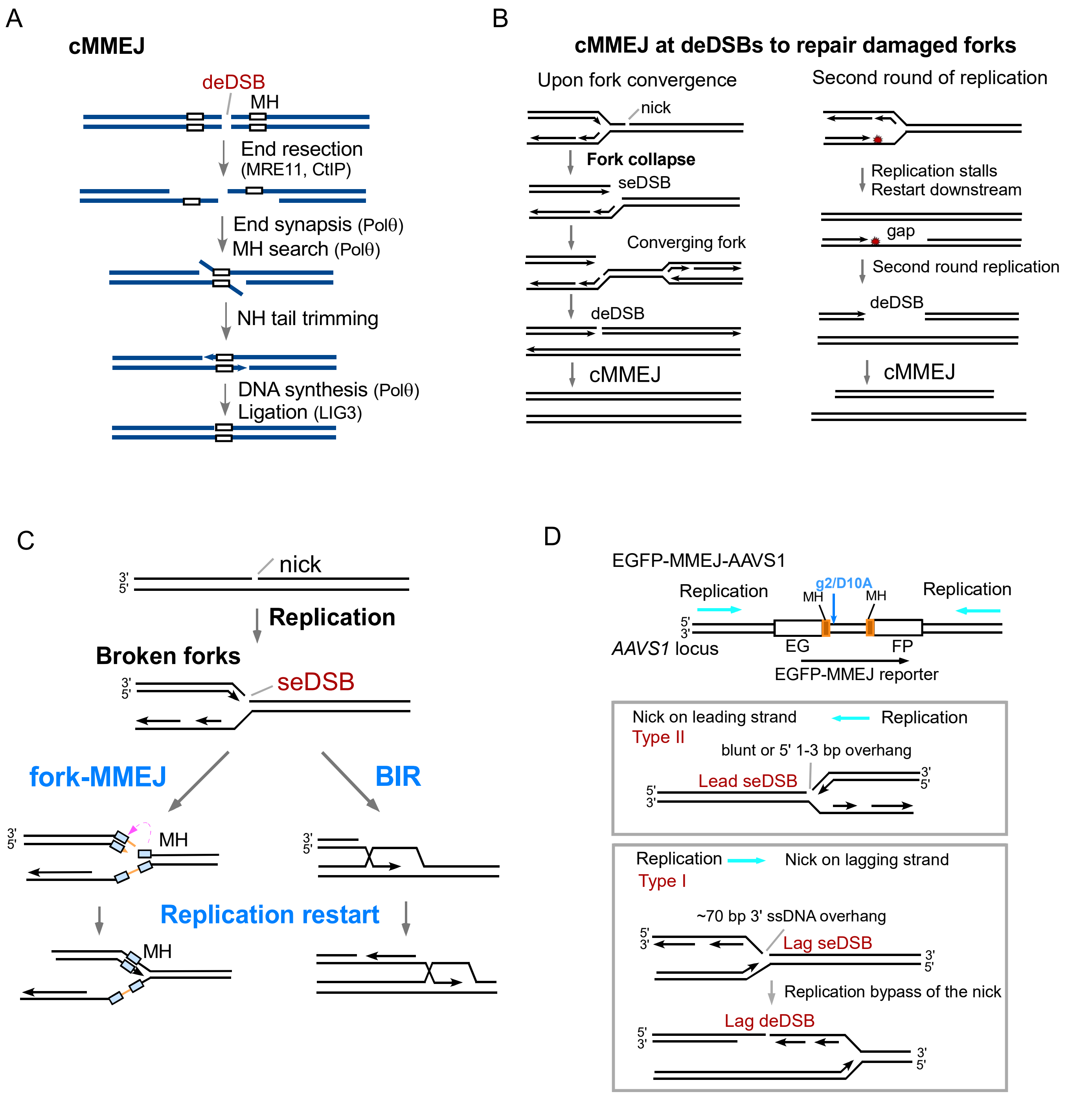


**Figure S1. Models for cMMEJ and fork-MMEJ.**
(A) Schematic representation of cMMEJ in the repair of deDSBs. MH: microhomology; NH tail: non-homologous tail. (B) Models for using cMMEJ to repair broken forks after deDSBs are generated by converging forks (left) or after second round of replication (right). (C) Schematic drawings illustrating the repair of seDSBs induced by nicks on replication forks through the fork-MMEJ and BIR pathways. (D) Schematic drawings depicting the EGFP-MMEJ reporter inserted in the *AAVS1* locus with indication of replication directions (top). When replication encounters the nicks generated by g2/Cas9$^{D10A}$, which would either create lead seDSBs or lag seDSBs/deDSBs, depending on replication directions.

Figure S2

A

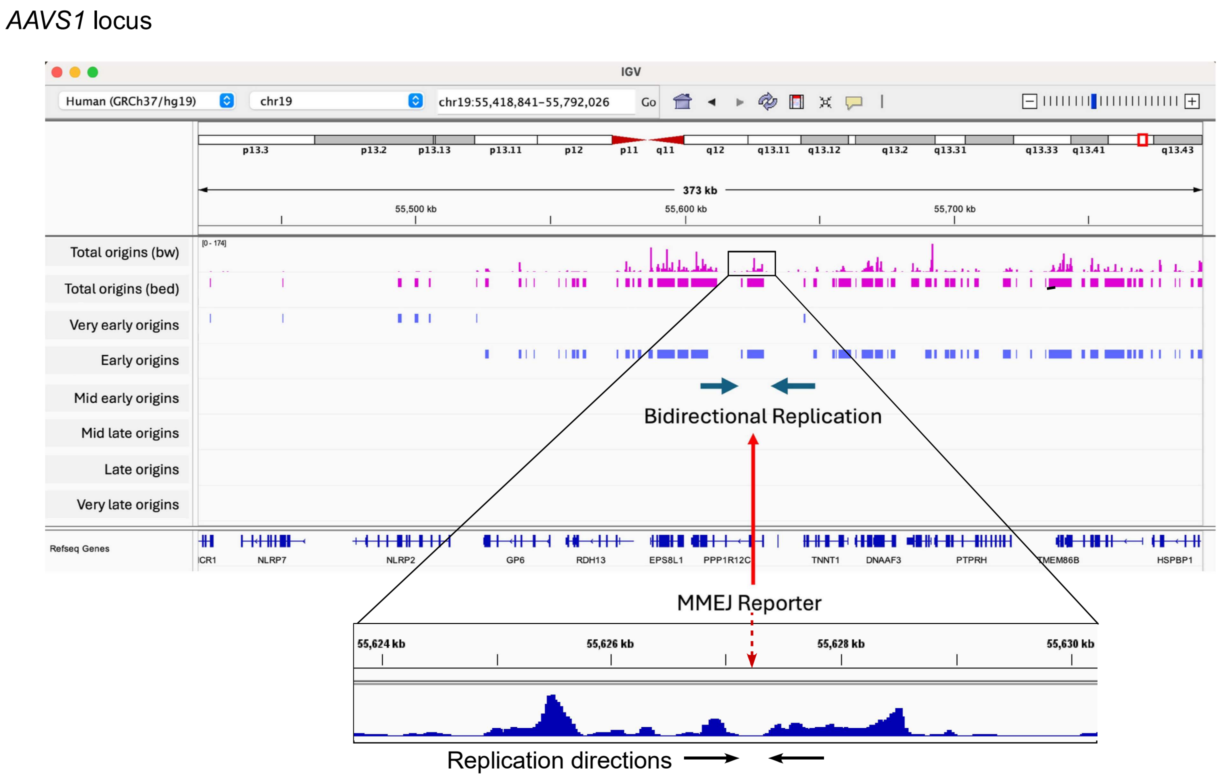


B

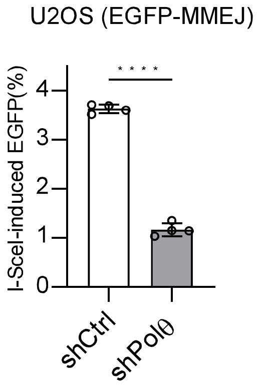


**Figure S2. MMEJ is dependent on Polθ.**

(A) The EGFP-MMEJ reporter was inserted into the *AAVS1* locus in U2OS cells and the insertion site is indicated by red arrows. Nascent strand sequencing analysis[42] was utilized here and revealed that the *AAVS1* locus in U2OS cells replicates early and bidirectionally. (B) The EGFP-MMEJ reporter cells expressing Polθ shRNA (shPolθ) or vector (shCtrl) were infected by lentiviruses encoding I-SceI. The percentage of EGFP positive cells was analyzed by FACS 4 days post infection.

Figure S3

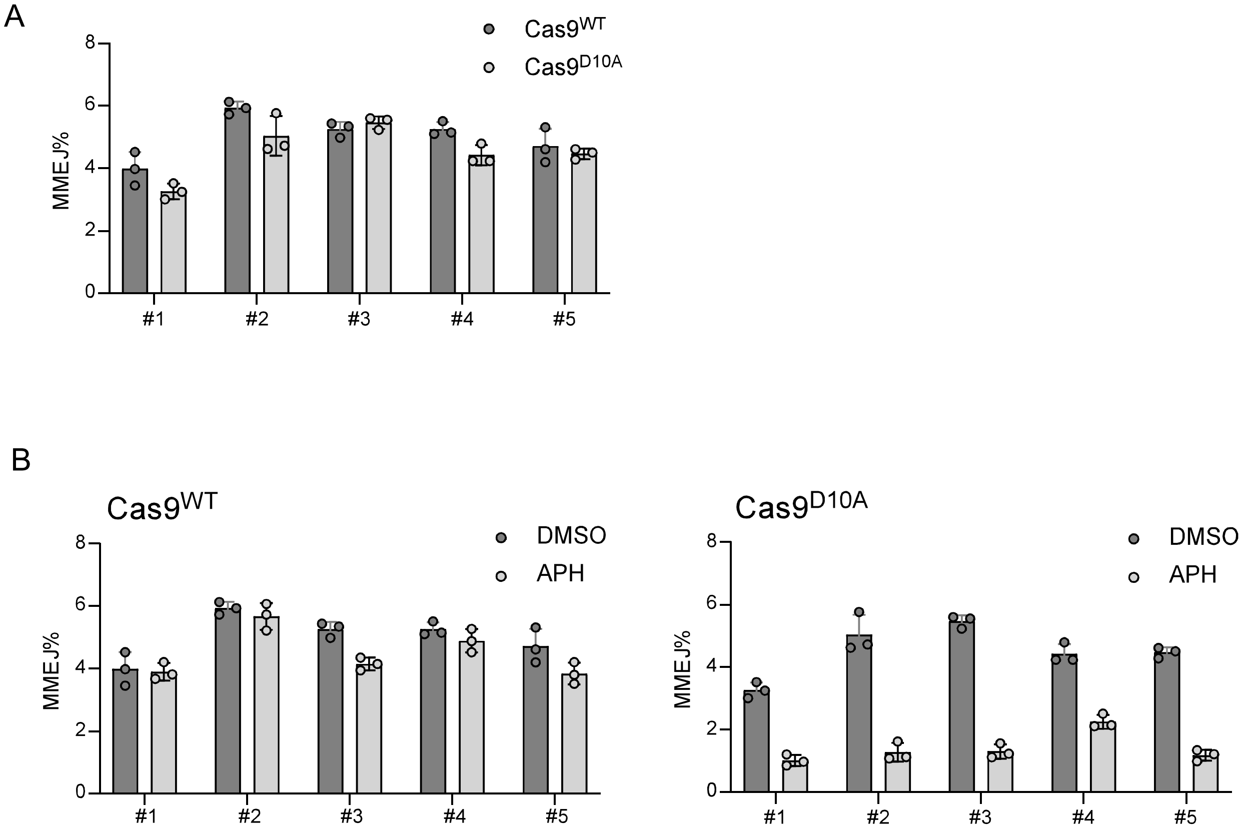


**Figure S3. The MMEJ assay was performed using the EGFP-MMEJ reporter randomly integrated into the U2OS genome.**

(A) Multiple single clones of the EGFP-MMEJ reporter randomly integrated into the U2OS genome were infected with lentiviruses encoding gRNA2/Cas9$^{WT}$ or gRNA2/Cas9$^{D10A}$, and MMEJ was assayed 4 days post infection. (B) U2OS (EGFP-MMEJ) single clones were infected with lentiviruses encoding gRNA2/Cas9$^{WT}$ (left) or gRNA2/Cas9$^{D10A}$ (right) with or without treatment of aphidicolin (APH, 0.4 uM), and assayed for MMEJ as described in (A).

### Figure S4

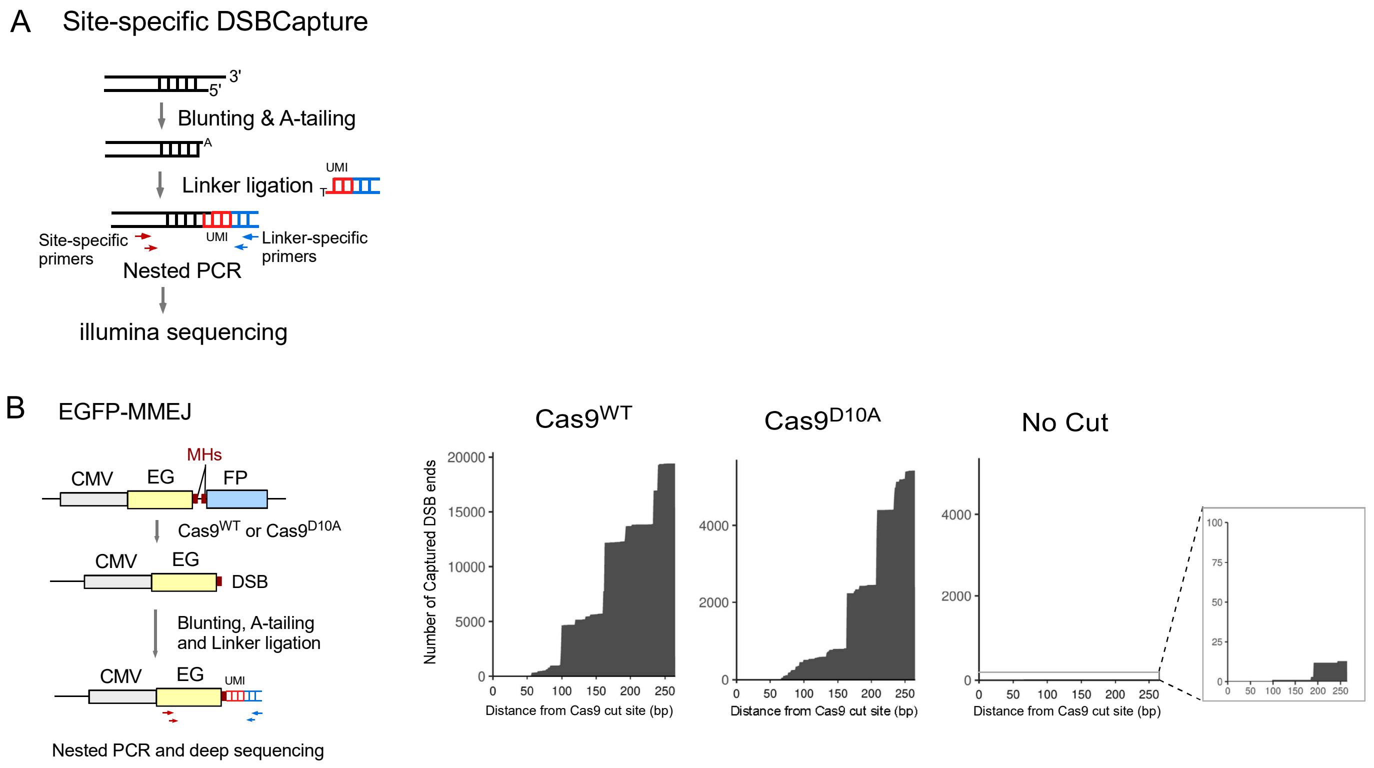


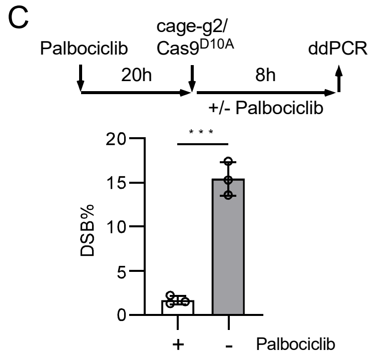


| | G1 | S | G2 |
|---|---|---|---|
| No release: | 54.6 | 6.5 | 28.3 |
| Release 8h: | 35.4 | 52.7 | 7.0 |

**Figure S4. Detection of DSB formation following Cas9 and Cas9n cleavage at the EGFP-MMEJ reporter.**
(A) Schematic drawing of modified DSBCapture to detect DSB formation at a specific genomic site. DSB ends were blunted and A-tailed, followed by ligation to a double-stranded adaptor containing unique molecular identifier (UMI). Nested PCR was then performed using adaptor-specific primers (marked in blue) and internal site-specific primers (marked in red), followed by illumina deep sequencing as described in the STAR Methods. PCR bias was eliminated by using UMI. (B) Site-specific DSBCapture was performed 24 hours after transfection of plasmids encoding g2/Cas9$^{WT}$ and g2/Cas9$^{D10A}$ to the U2OS (EGFP-MMEJ) reporter cells using no cut as a control. The positions of nested PCR primers on the EGFP-MMEJ reporter are indicated (left), and the sequences of captured DSB reads are aligned to the target sequence (right). The number of captured DSBs (y-axis) are plotted as a function of distance from the Cas9 cut site (x-axis). (C) EGFP-MMEJ reporter cell line transfected with caged gRNA2 (caged-g2) and Cas9$^{D10A}$ was treated with Palbociclib (0.25 uM, 20 hours) to arrest U2OS cells in G1 phase, followed by 365 nm light activation. After another 8-hour incubation with or without Palbociclib (arrested in G1 or released from G1, as shown by cell cycle analysis), genomic DNA was extracted and analyzed by ddPCR for DSB quantification.

Figure S5

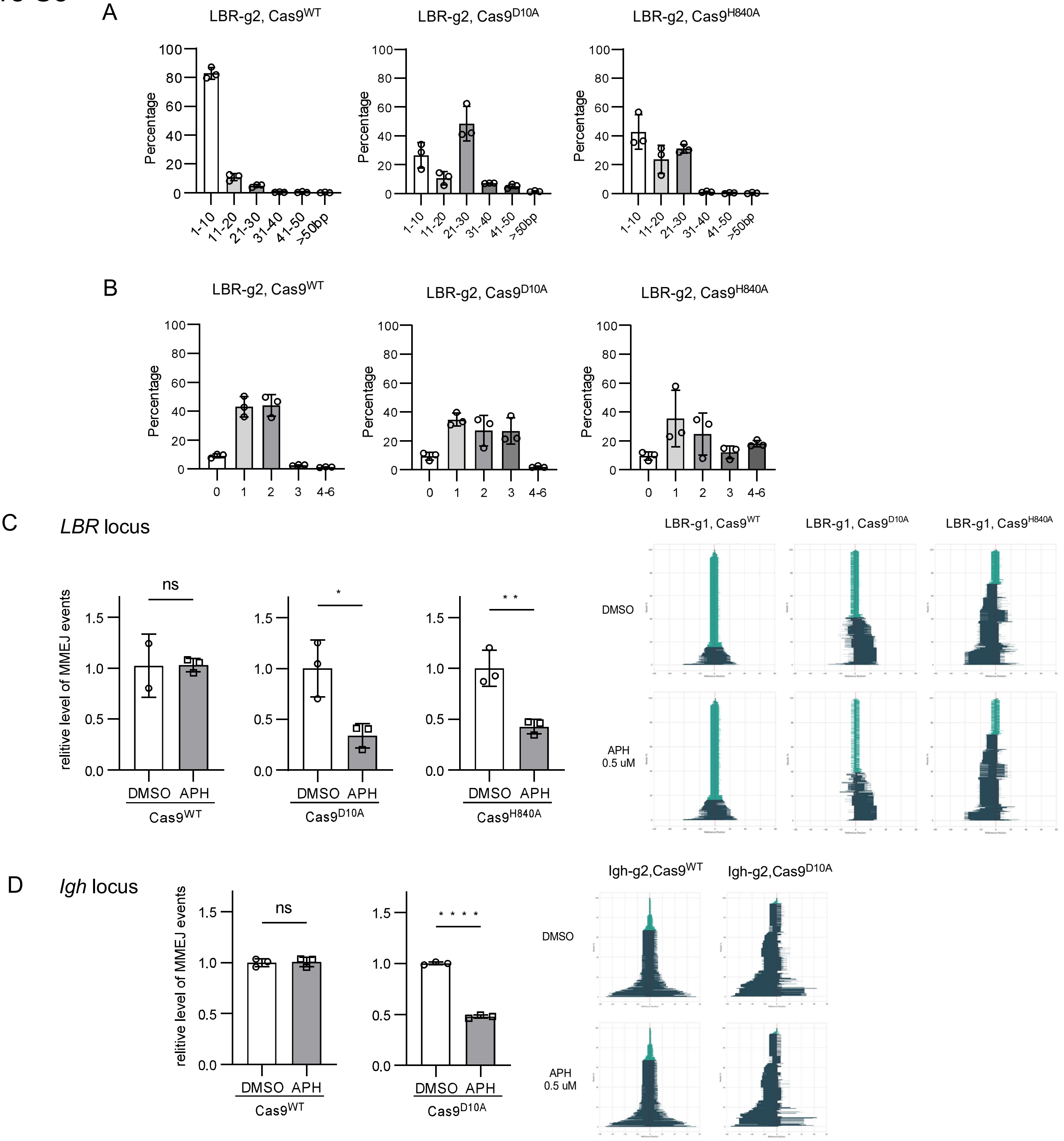


**Figure S5. Features of MMEJ repair at genomic loci after Cas9 and Cas9n cleavage.**
(A, B) Distributions of deletion sizes (A) and microhomology sizes (B) were analyzed by deep sequencing after cleavage of the human *LBR* locus by LBR-g2 with $Cas9^{WT}$, $Cas9^{D10A}$ or $Cas9^{H840A}$. Related to Fig.1D. (C, D) U2OS cells (C) or mES cells (D), treated with APH (0.4 uM) or DMSO, were infected with lentiviruses expressing LBR-g1 (C) or transfected with plasmids encoding Igh-g2 (D), along with $Cas9^{WT}$, $Cas9^{D10A}$ or $Cas9^{H840A}$, followed by deep sequencing to analyze repair products at the cleavage site. Related to Fig. 1E (C) and Fig. 4F (D). The percentage of MMEJ events were normalized to control cells treated with DMSO (left). The indel patterns was compared between samples with and without APH treatment (right), with deletion sizes color-coded as described in Fig. 1E.

Figure S6

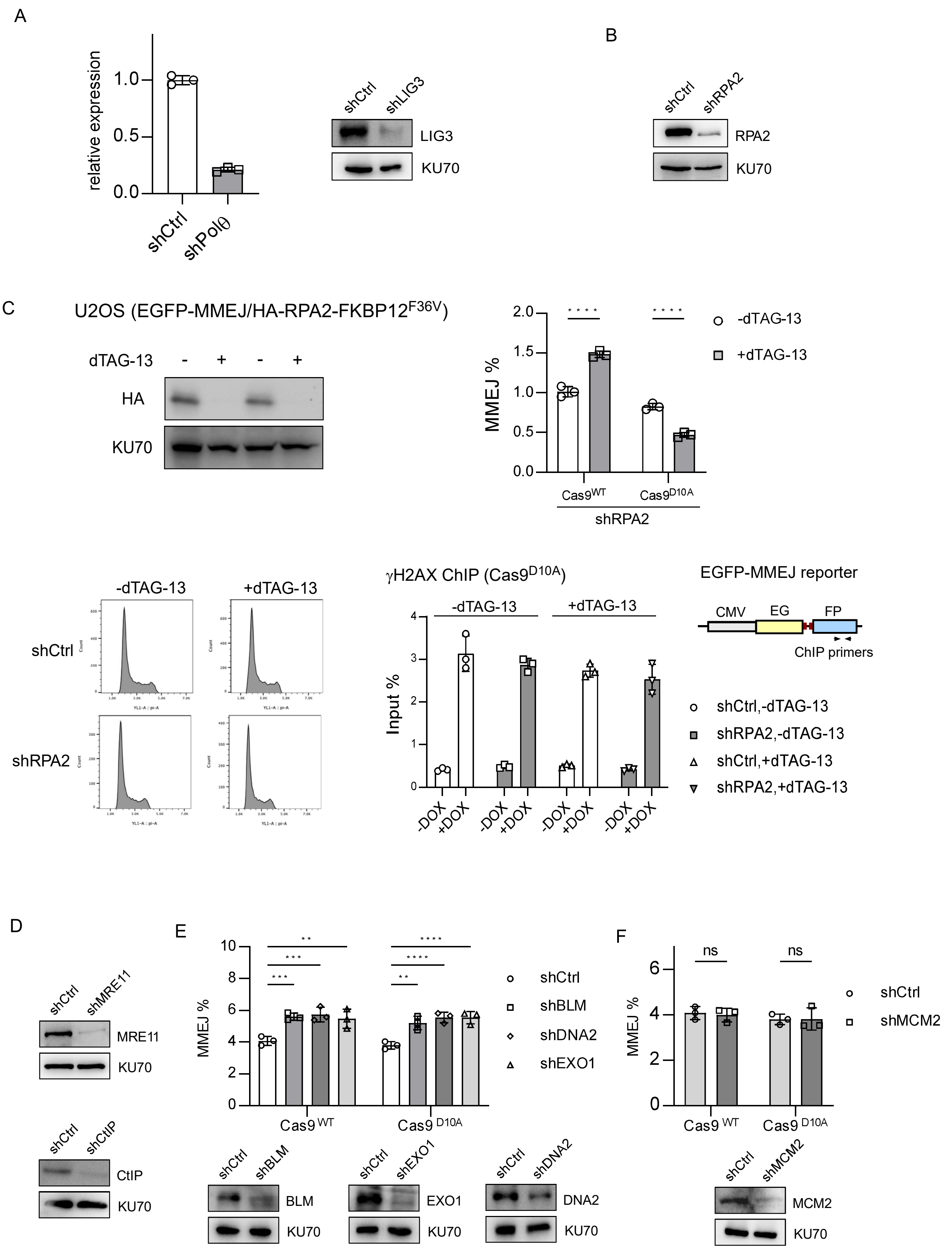
A
relative expression
1.0
0.5
0.0
shCtrl
shPolθ
shCtrl
shLIG3
LIG3
KU70
B
shCtrl
shRPA2
RPA2
KU70
C
U2OS (EGFP-MMEJ/HA-RPA2-FKBP12F36V)
dTAG-13
-
+
-
+
HA
KU70
MMEJ %
2.0
1.5
1.0
0.5
0.0
****
****
-dTAG-13
+dTAG-13
Cas9WT
Cas9D10A
shRPA2
-dTAG-13
+dTAG-13
shCtrl
shRPA2
γH2AX ChIP (Cas9D10A)
-dTAG-13
+dTAG-13
Input %
4
3
2
1
0
-DOX
+DOX
-DOX
+DOX
-DOX
+DOX
-DOX
+DOX
EGFP-MMEJ reporter
CMV
EG
FP
ChIP primers
shCtrl,-dTAG-13
shRPA2,-dTAG-13
shCtrl,+dTAG-13
shRPA2,+dTAG-13
D
shCtrl
shMRE11
MRE11
KU70
shCtrl
shCtIP
CtIP
KU70
E
MMEJ %
10
8
6
4
2
0
**
***
***
****
****
**
Cas9 WT
Cas9 D10A
shCtrl
shBLM
shDNA2
shEXO1
shCtrl
shBLM
BLM
KU70
shCtrl
shEXO1
EXO1
KU70
shCtrl
shDNA2
DNA2
KU70
F
MMEJ %
6
4
2
0
ns
ns
Cas9 WT
Cas9 D10A
shCtrl
shMCM2
shCtrl
shMCM2
MCM2
KU70

**Figure S6. Investigating the mechanism underlying fork-MMEJ.**

(A) Knock-down efficiency of Pol$\theta$ was assessed by RT-qPCR (left), and LIG3 by Western blot analysis (right) in U2OS (EGFP-MMEJ) cells. Related to Fig. 2A. (B) RPA2 knock-down efficiency in U2OS (EGFP-MMEJ) cells was determined by Western blot analysis using KU70 as the loading control. Related to Fig. 2B. (C). U2OS (EGFP-MMEJ/HA-RPA2-FKBP12$^{F36V}$) cells were treated with dTAG-13 (500 nM) and Western blot analysis was performed 6 hours later (top left). Cas9$^{D10A}$ expression was induced by doxycycline (DOX, 2 mM) 6 hours after dTAG-13 addition. 48 hours after RPA2 degradation induced by dTAG-13, FACS was performed (top right) and the cell cycle profile was determined by propidium iodide staining (bottom left). $\gamma$H2AX ChIP analysis using primers 0.1 kb away from the Cas9 cleavage site on the EGFP-MMEJ reporter was conducted 24 hours after DOX addition to induce Cas9$^{D10A}$ expression (bottom right). (D) Knock-down efficiency of MRE11 and CtIP in U2OS (EGFP-MMEJ) cells was determined by Western blot analysis using KU70 as the loading control. Related to Fig.2C. (E, F) U2OS (EGFP-MMEJ) cells expressing shRNAs for BLM, DNA2, EXO1 (E) or MCM2 (F) were infected with lentiviruses coding gRNA2 along with Cas9$^{WT}$ or Cas9$^{D10A}$, and assayed by FACS analysis 4 days post-infection. Expression levels of indicated proteins after depletion by shRNAs were shown by Western blot analysis.

### Figure S7

A

mES *ROSA* locus

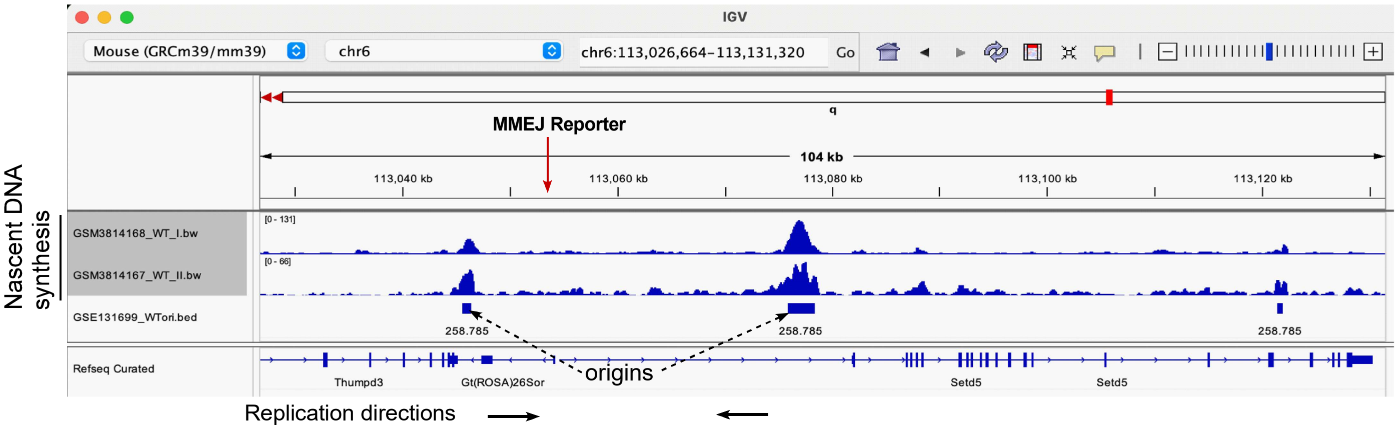


B

*POLQ*-KO-A9 and C4 clones

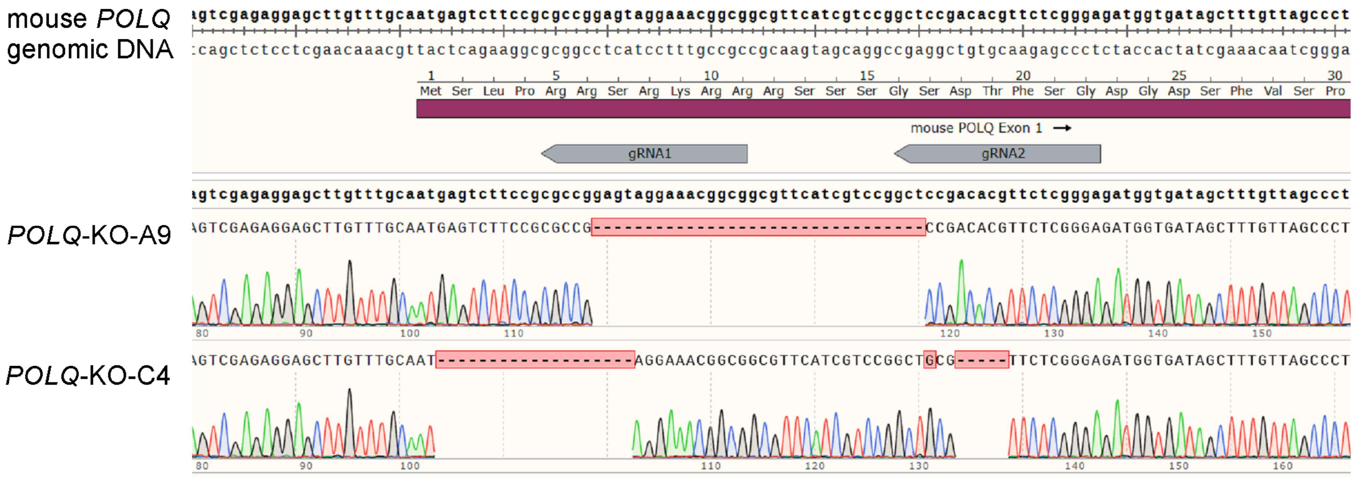


**Figure S7. Polθ-mediated fork-MMEJ is involved in the repair of seDSB.**
(A) The EGFP-MMEJ reporter was integrated into the *ROSA* locus in mES cells. The *ROSA* locus replicates bidirectionally as revealed by analysis of reported nascent DNA sequencing data (GEO accession number: GSM3814167, GSM3814168, and GSE131699)[58]. (B) *POLQ* knock-out (KO) in mES cells was verified by genomic DNA PCR sequencing. Two gRNAs targeting Exon 1 of *POLQ* are indicated, and the sequencing results from the *POLQ*-KO clones are aligned with the original *POLQ* genomic DNA sequence. Related to Fig. 2D.

Figure S8

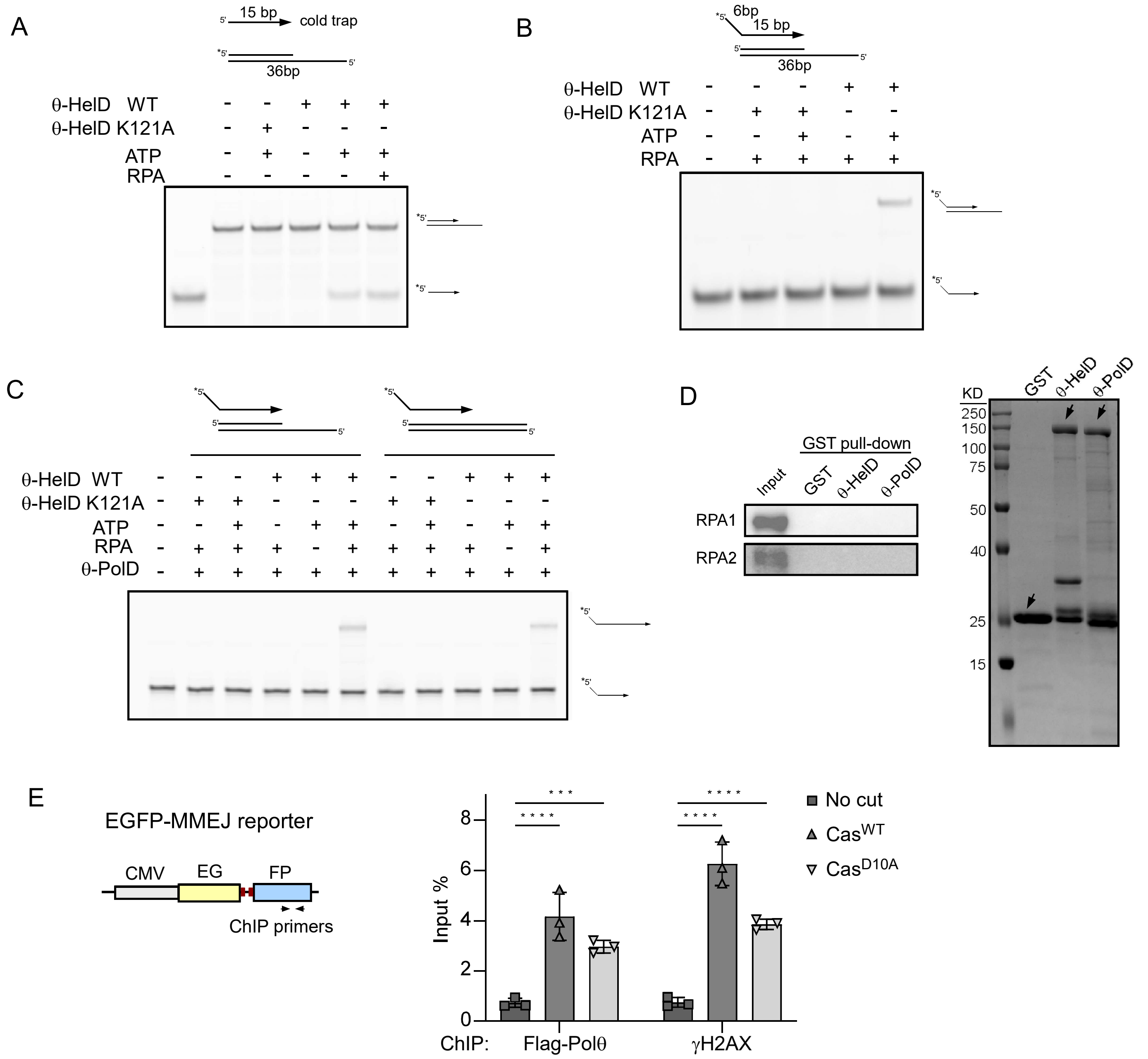


**Figure S8. Polθ-PolD, Polθ-HelD and RPA are required for fork-MMEJ.**
(A) 5 nM Pre-annealed $^{32}$P-5'-labeled 15-nt ssDNA and a complementary 36-nt DNA strand were mixed with 200 nM cold trap (15-nt), and incubated with Polθ-HelD-WT or K121A, ATP, and RPA complex as indicated. Reaction products were resolved on a non-denaturing gel. (B) 10 nM $^{32}$P-5'-labeled ssDNA mixed with 5 nM pre-annealed DNA substrate was incubated with Polθ-HelD WT or K121A, ATP, and RPA complex as indicated, and the products were resolved on a non-denaturing gel. (C) 10 nM $^{32}$P-5'-labeled ssDNA mixed with 5 nM pre-annealed DNA substrate was incubated with Polθ-HelD$^{WT}$ or K121A, ATP and RPA complex as indicated, followed by extension with Polθ-PolD and dNTPs. The reaction products were then resolved on a denaturing gel. (D) Purified GST-fused Polθ-PolD and Polθ-HelD, along with GST, were pre-bound to glutathione resin and incubated with purified RPA complex. Western blot analyses of RPA1 and RPA2 were performed on the GST pull-down samples. (E) U2OS (EGFP-MMEJ) cells expressing Flag-Polθ were infected with lentiviruses encoding gRNA2 and Cas9$^{WT}$ or Cas9$^{D10A}$. The recruitment of Flag-Polθ and γH2AX to the cleavage site was assayed by ChIP using primers 0.1 kb away from the cleavage site 24 hours after gRNA2/Cas9$^{WT}$ or gRNA2/Cas9$^{D10A}$ infection.

Figure S9

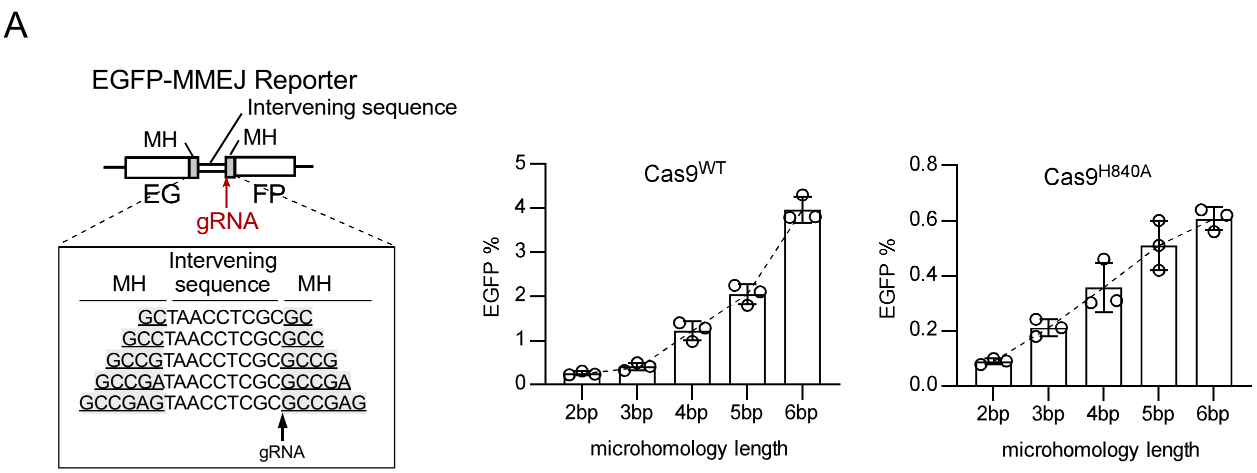


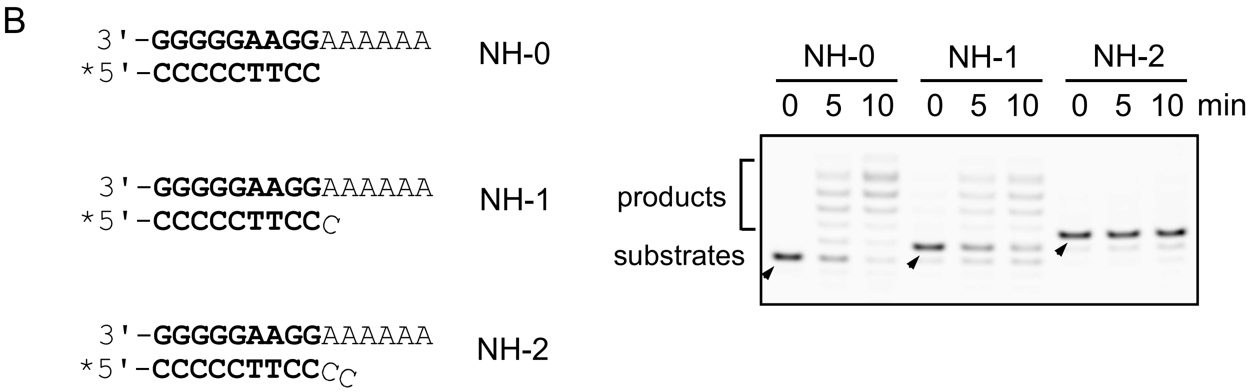


**Figure S9. The effect of microhomology (MH) and non-homologous (NH) tail length on MMEJ repair efficiency.**
(A) U2OS cells carrying EGFP-MMEJ reporters with different MH sizes as indicated (left) were assayed for MMEJ following infection with lentiviruses encoding gRNA/Cas9$^{WT}$ (middle) or gRNA/Cas9$^{H840A}$ (Right). (B) $^{32}$P-5'-labeled pre-annealed DNA with 3'-non-homologous tails of different length was incubated with Polθ-PolD and dNTPs for the indicated time and the repair products were resolved on a denaturing gel.

Figure S10

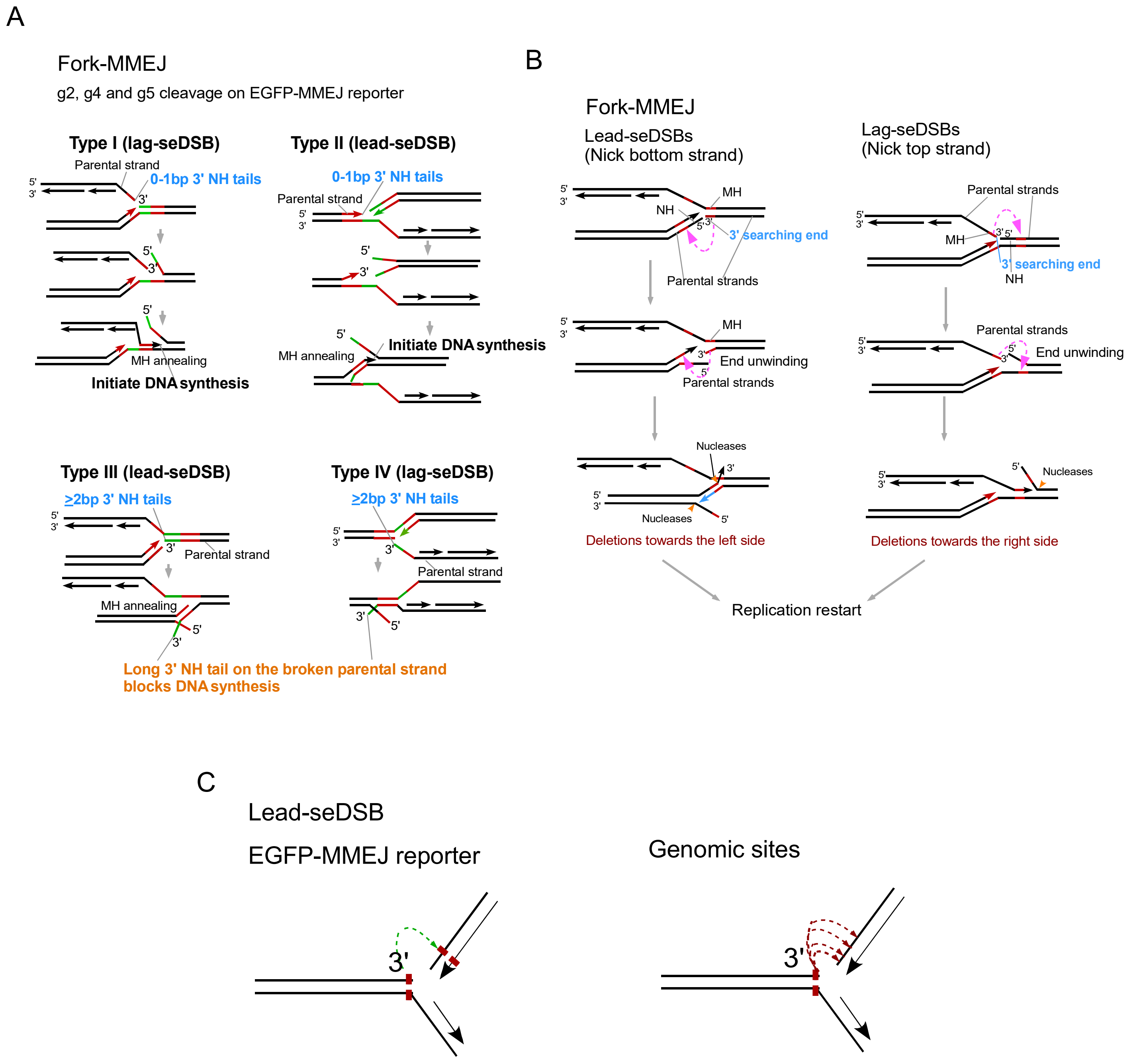


**Figure S10. Fork-MMEJ utilizes the 3' end of broken parental strands to directly search for microhomology (MH) internal of the other broken end.**
(A) Schematic drawings to illustrate that fork-MMEJ accommodates no or 1 bp (0-1 bp) non-homologous (NH) tails (top, Type I/II seDSBs), but not >2 bp NH tails (bottom, Type III/IV seDSBs), on the 3' end of broken parental strands in our EGFP-MMEJ reporter. Related to Fig. 3B. (B) At genomic loci, fork-MMEJ uses the 3' broken end (designated as 3' searching end) on the broken parental strands to search for the microhomology (MH) internal of the other broken end, leading to asymmetric deletion patterns. (C) While the EGFP-MMEJ reporter only scores a fixed event using designed MHs, at genomic sites, the 3' end of the broken parental strands can search for MHs within a relatively big region of the other broken end.

Figure S11

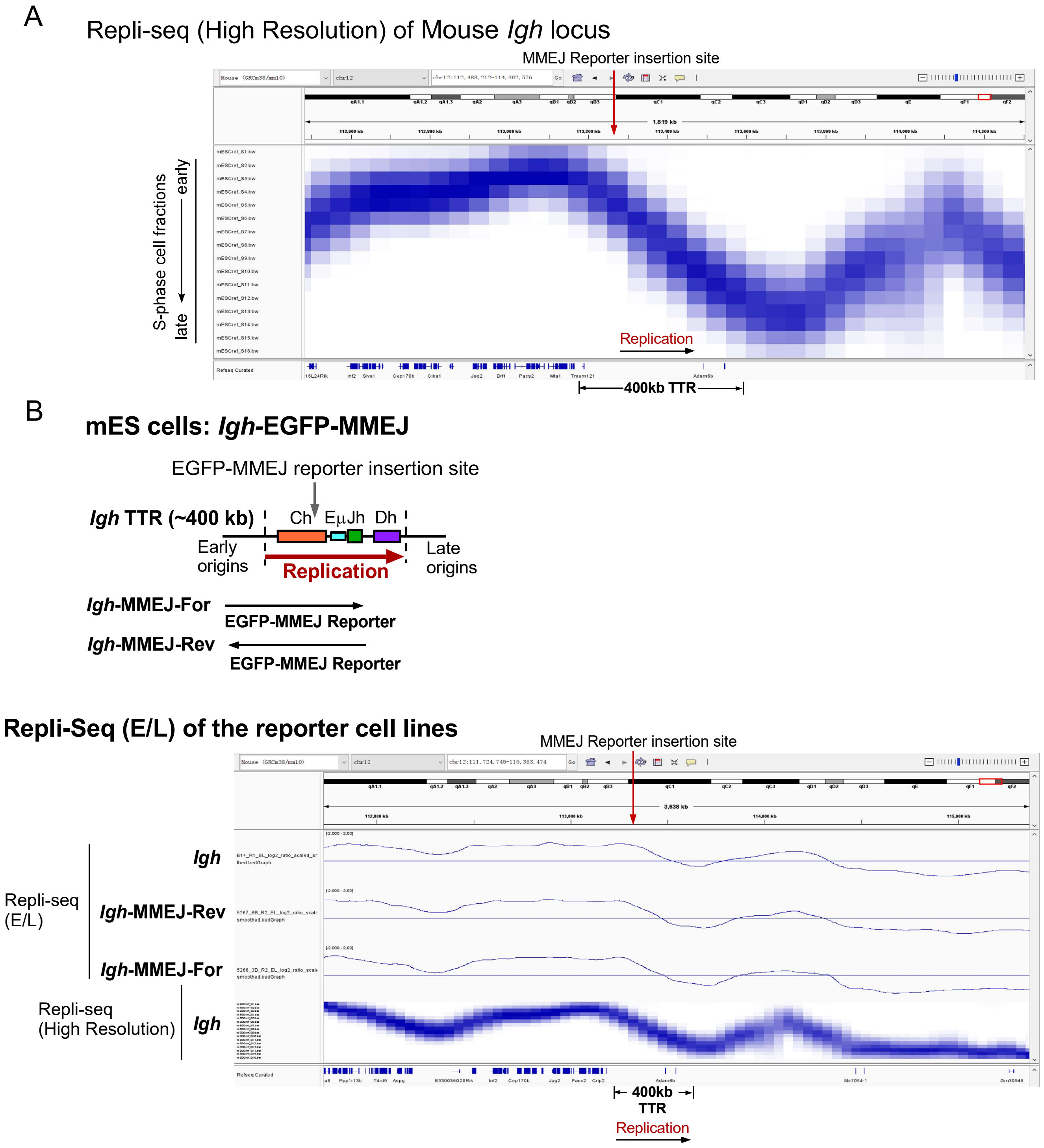


**Figure S11. Repli-seq analysis of mES cells with and without insertion of EGFP-MMEJ reporters at the Igh locus.**
(A) High-resolution Repli-Seq analysis was performed in mES cells, and the results at the mouse Igh locus are shown. The 400 kb TTR displays a uniform slope from early to late replication, consistent with the replication fork progression rate in mammalian cells, indicating unidirectional replication in this region. (B) Schematic illustration showing the insertion site and orientations of the EGFP-MMEJ reporter in the mouse *Igh* locus (top). The replication direction of the TTR (400kb) region in *Igh* is indicated. Repli-Seq (E/L) analysis was performed in mES cells with or without integration of the EGFP-MMEJ reporter in two different orientations (For and Rev) at the *Igh* locus. Representative results at the mouse *Igh* locus are shown (bottom).

Figure S12

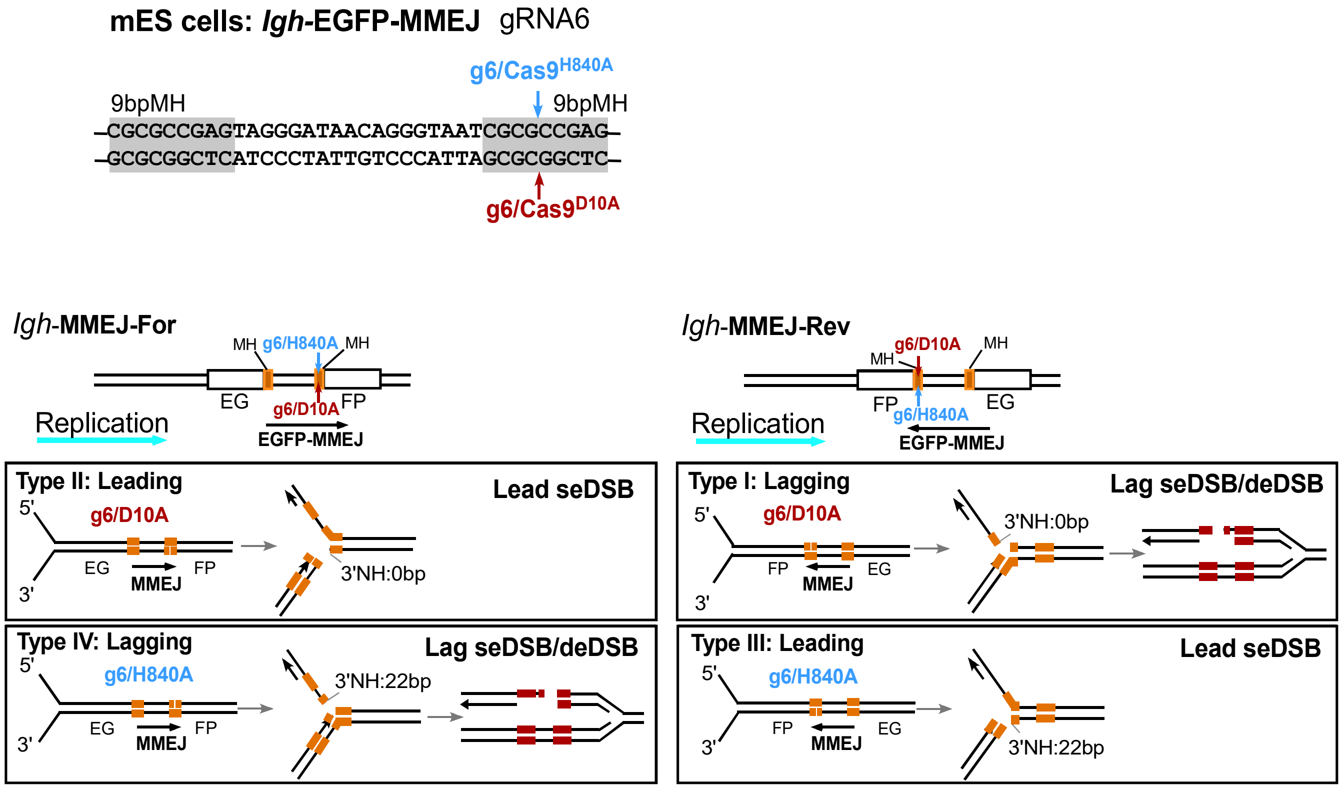


**Figure S12. Analysis of fork-MMEJ using the EGFP-MMEJ reporter inserted at the *Igh* locus of mES cells, where replication is unidirectional.**

The EGFP-MMEJ reporter was integrated into the *Igh* TTR locus in mES cells in two different orientations (*Igh*-MMEJ-For and *Igh*-MMEJ-Rev). The four types of seDSB generated after cleavage by gRNA6 along with Cas9$^{D10A}$ or Cas9$^{H840A}$ are illustrated. Unidirectional replication at the *Igh* locus in reference to the orientation of the MMEJ reporter is indicated.

Figure S13

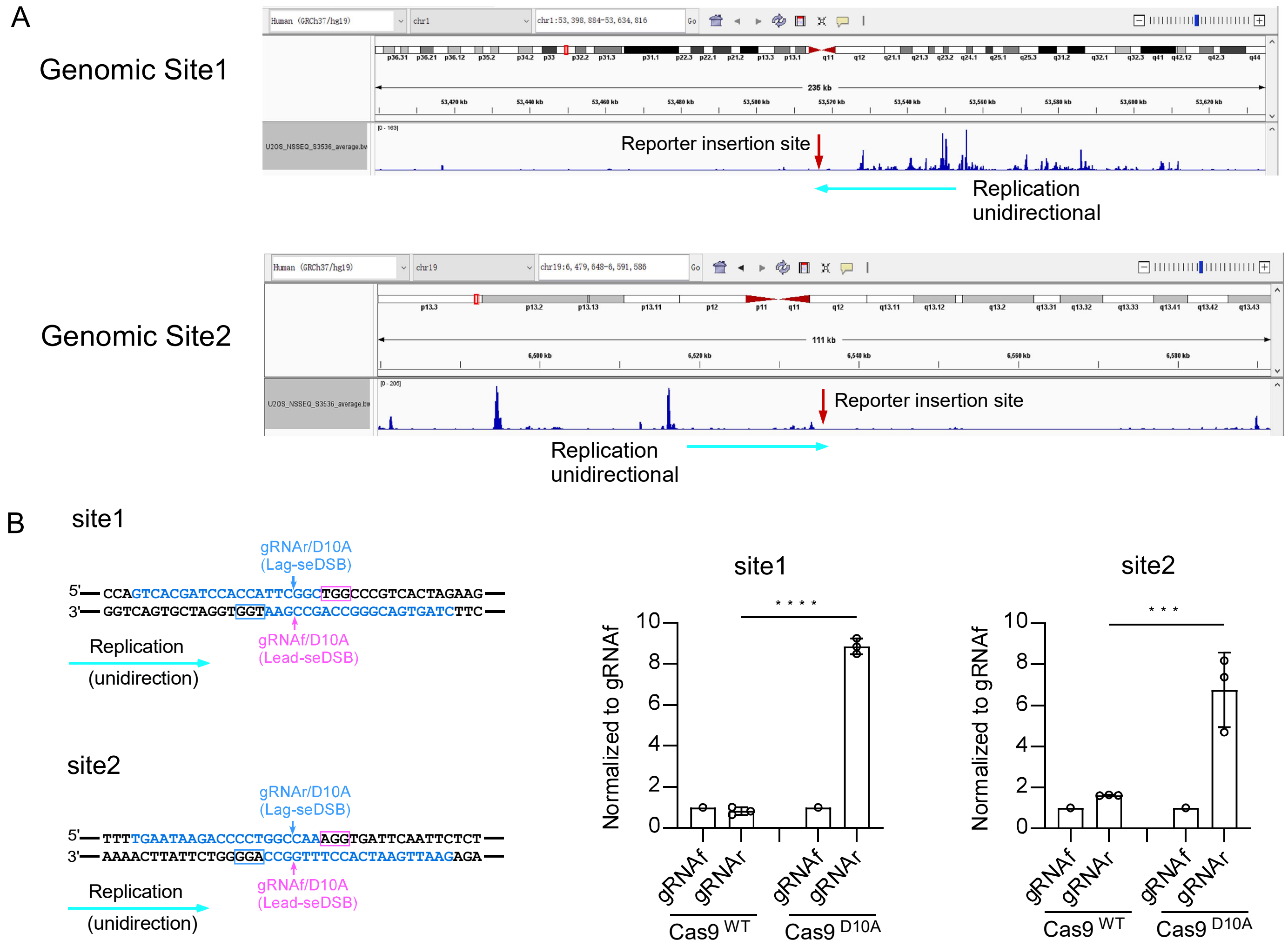


**Figure S13. Repair indels with MMEJ features are accumulated more on the leading strands upon Cas9n cleavage.**
(A) Nascent strand sequencing data from U2OS cells are shown for two genomic sites (GRCh37, site 1: chr1, 53519413-53519629, site 2: chr19, 6536158-6536368) with unidirectional replication. The gRNA cleavage sites are indicates. (B) The genomic site 1 and site 2 described in (A) were cleaved in U2OS cells by indicated gRNAs with Cas9$^{WT}$ and Cas9$^{D10A}$, followed by deep sequencing analysis of the cleavage sites. The gRNA cleavage sites in relationship with replication directions for the two sites are shown (top). The normalized frequency of indels with MMEJ features (>4 bp deletion and >1 bp MH) was calculated by setting the frequency of gRNAf/Cas9$^{WT}$ and gRNAf/Cas9$^{D10A}$ to 1, respectively.

Figure S14

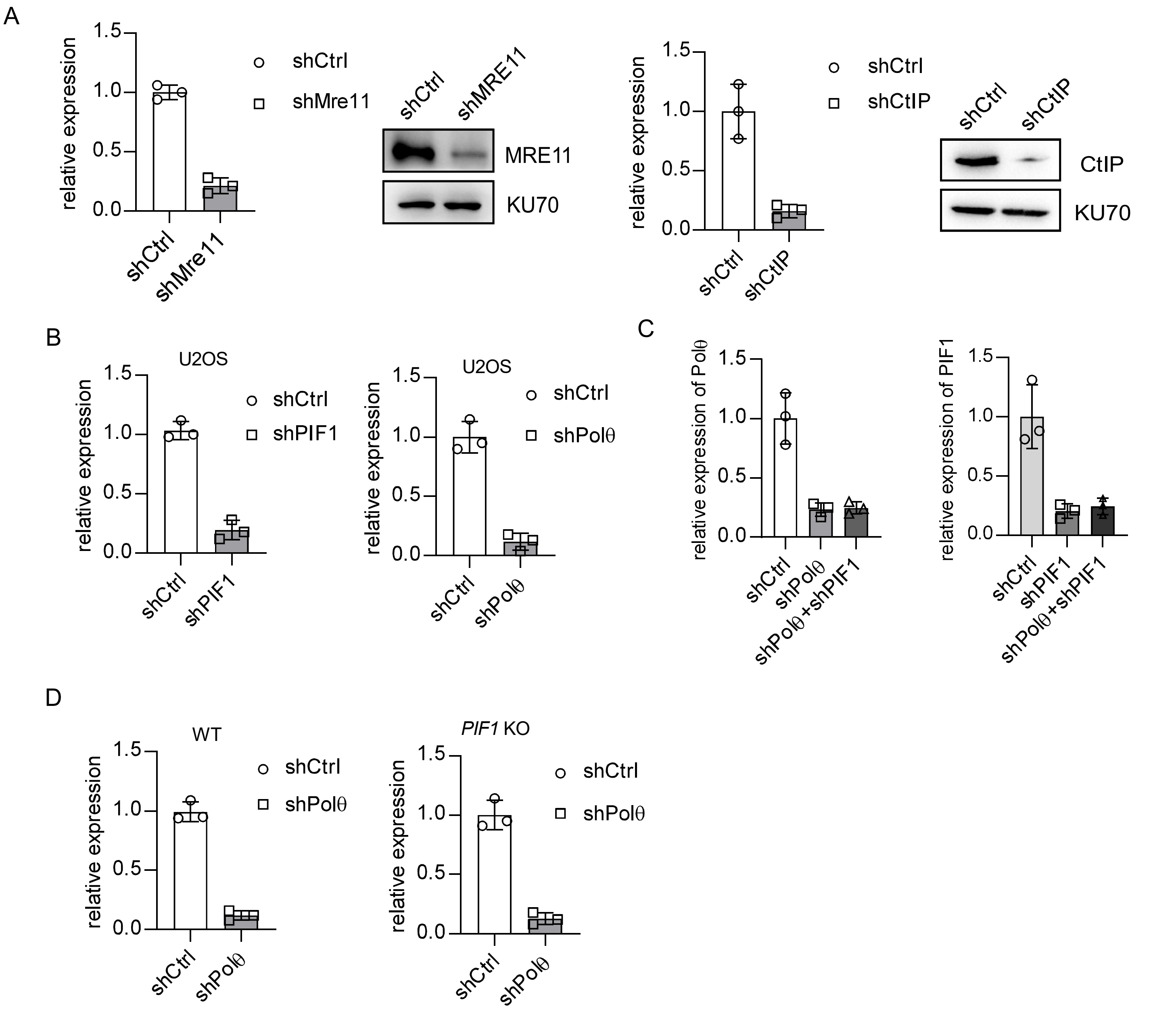


**Figure S14. Examining the knock-down efficiency induced by shRNAs.**
(A) Knock-down efficiency of MRE11 and CtIP in U2OS (EGFP-MMEJ/mCherry-BIR) cells was determined by RT-qPCR and Western blot analysis. Related to Fig. 5D. (B) Knock-down efficiency of PIF1 and Polθ in U2OS (EGFP-MMEJ/mCherry-BIR) cells was determined by RT-qPCR. Related to Fig.5E and 5F. (C) Polθ knock-down efficiency in WT and *PIF1*-KO U2OS cells was determined by RT-qPCR. Related to Fig. 6A. (D) Knock-down efficiency of PIF1 and Polθ in U2OS cells was determined by RT-qPCR. Related to Fig. 6B.

Figure S15

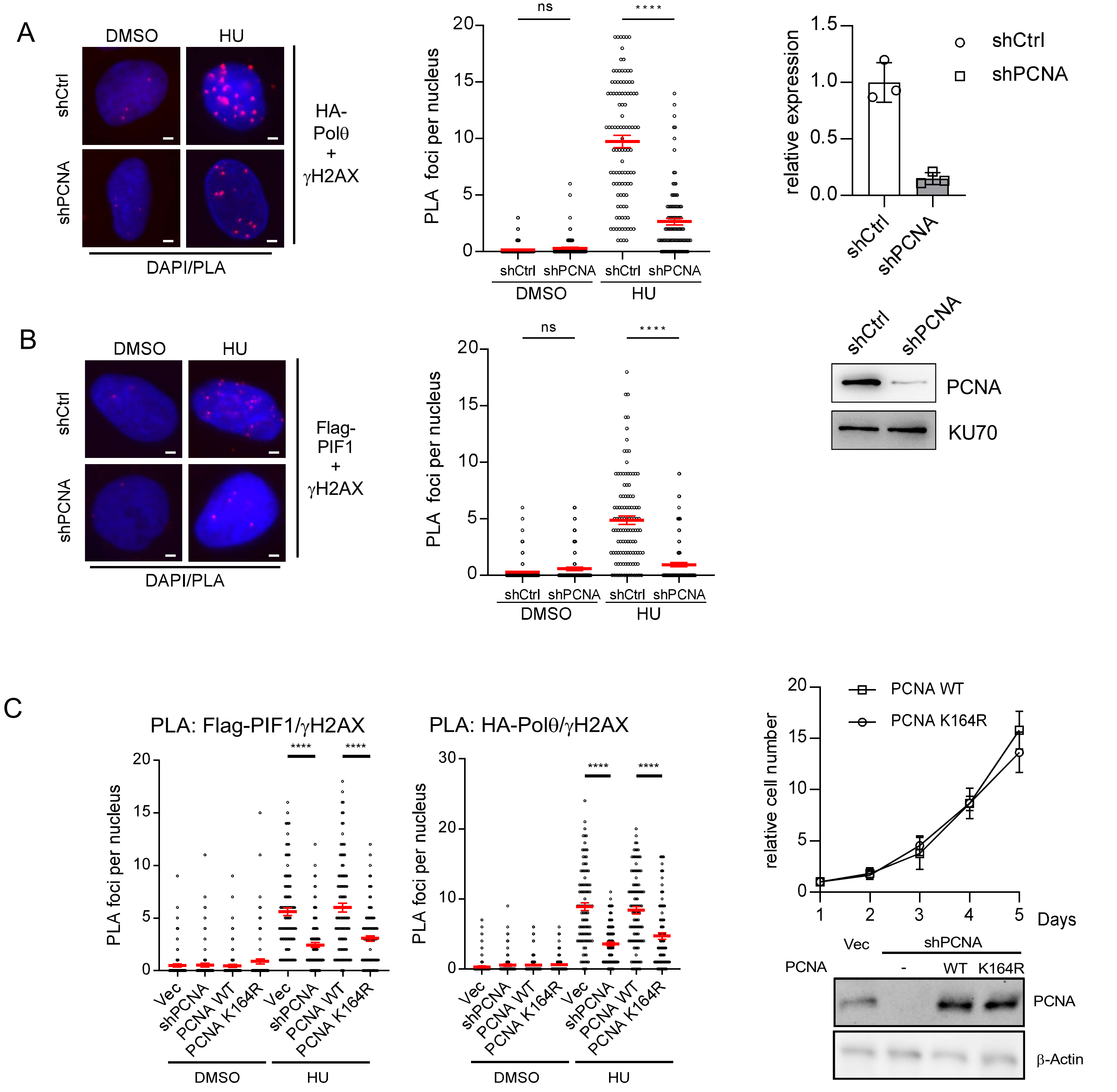


**Figure S15. PCNA ubiquitination is required for recruitment of Polθ and PIF1 to DSBs upon replication stress.**

(A) PLA was performed to show co-localization of HA-Polθ and γH2AX in U2OS cells expressing PCNA shRNA or shCtrl with or without treatment of HU (2 mM, 24 hours) (left). PCNA knock-down was determined by RT-qPCR and western blot (right). (B) PLA was performed to show co-localization of Flag-PIF1 and γH2AX in U2OS cells expressing PCNA shRNA or shCtrl with or without treatment of HU (2 mM, 24 hours). (C) PLA was performed to show co-localization of Flag-PIF1 (left) and HA-Polθ (middle) with γH2AX in U2OS cells expressing PCNA-WT, K164R mutant or the vector with endogenous PCNA silenced by shRNA, with or without treatment of HU (2 mM, 24 hours). Growth curves of U2OS cells expressing PCNA-WT or K164R mutant with endogenous PCNA silenced by shRNA are shown (right top) and the expression of PCNA WT and K164R along with endogenous PCNA was examined by western blot (right bottom). Scale bar, 2 um.

Figure S16

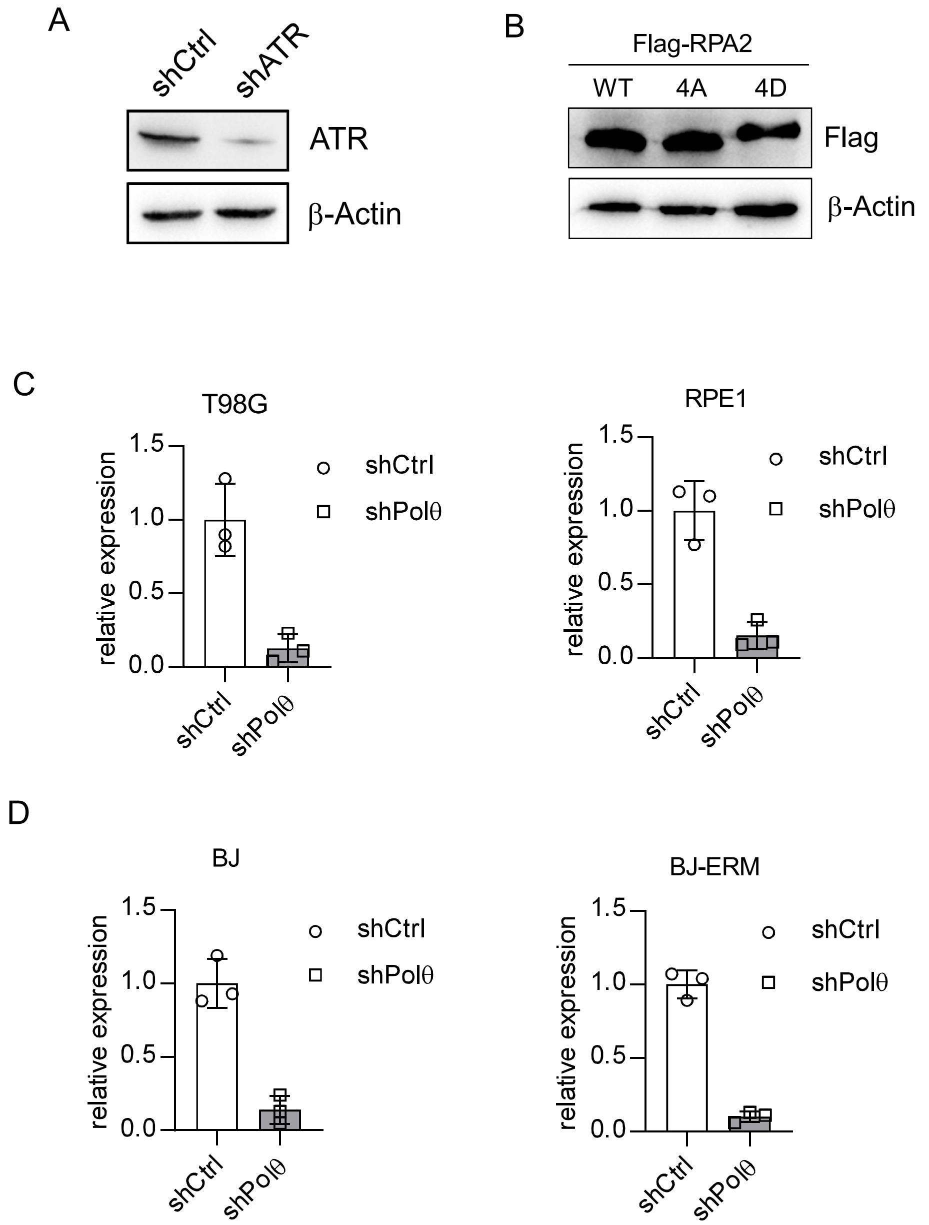


**Figure S16. shRNA Knock-down efficiency and protein expression levels.**
(A) ATR knock-down efficiency in U2OS (EGFP-MMEJ) cells was determined by RT-qPCR and western blot. Related to Fig. 7C. (B) Expression of Flag-RPA2 WT as well as RPA2-4A and 4D mutants in U2OS (EGFP-MMEJ) cells with endogenous RPA2 depleted by shRNA was assayed by western blot. Related to Fig. 7D. (C) Polθ knock-down efficiency in T98G and RPE1 cell lines was determined by RT-qPCR. Related to Fig. 7E. (D) Polθ knock-down efficiency in BJ and BJ-ERM cell lines was determined by RT-qPCR. Related to Fig. 7F.

Figure S17

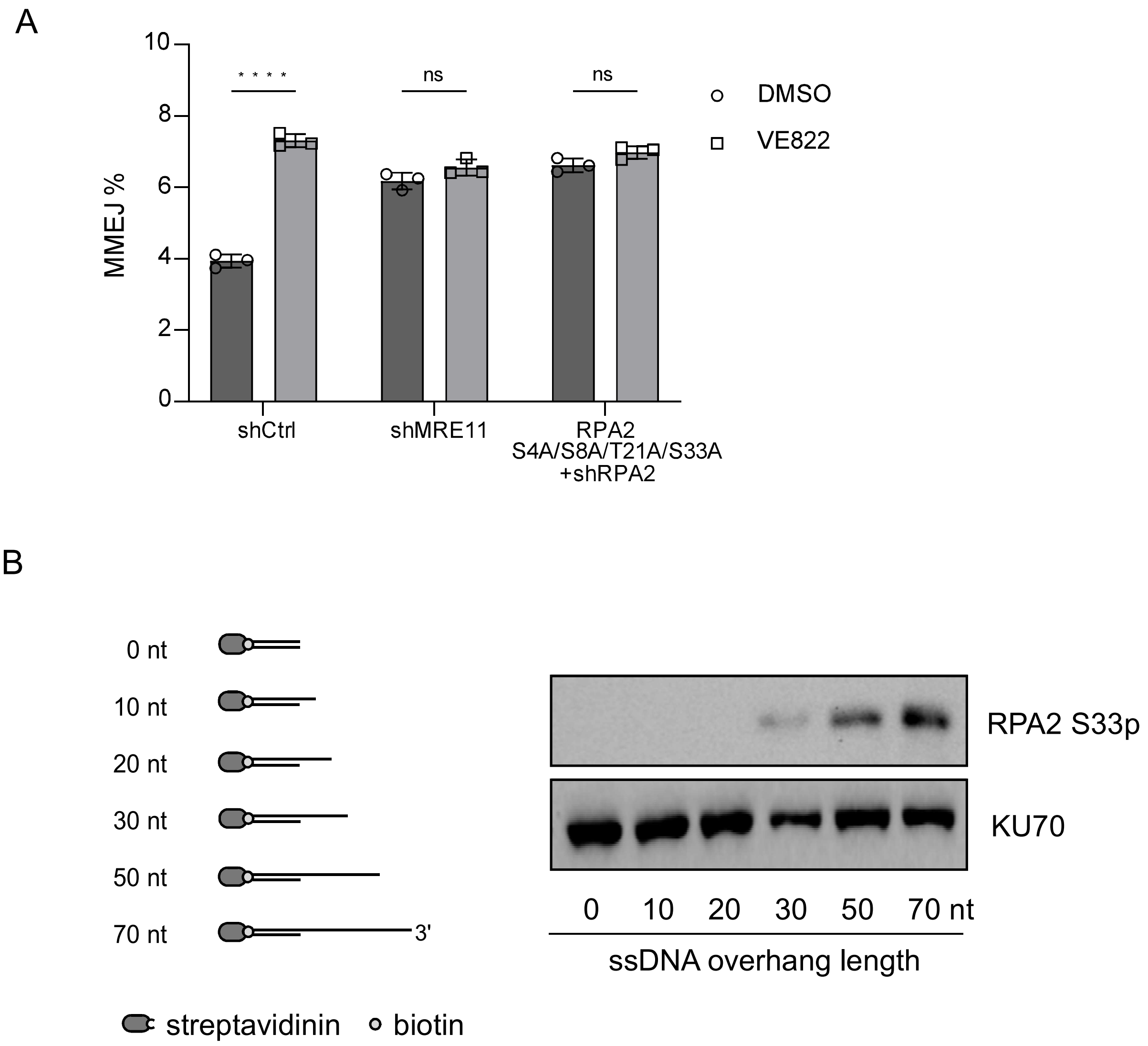


**Figure S17. RPA2 was phosphorylated in DSB end resection, which is required for fork-MMEJ.**

(A) The EGFP-MMEJ reporter cells expressing shRNA vector, MRE11 shRNA or RPA-S4A/S8A/T21A/S33A (RPA-4A) with endogenous RPA2 silenced by shRNA were treated with DMSO or VE822 (0.5 uM) and assayed for MMEJ following infection by lentiviruses expressing gRNA2/Cas9$^{D10A}$. (B) Nuclear extracts prepared from U2OS cells were incubated with one-end biotinylated DNA substrates containing 30 bp double stranded DNA with varying lengths of single-stranded overhangs as indicated, in the presence of streptavidin, as well as MRE11 inhibitor Mirin (1 mM) to prevent end resection. RPA2 phosphorylation at S33 was analyzed by western blot analysis with KU70 as a loading control.

Figure S18

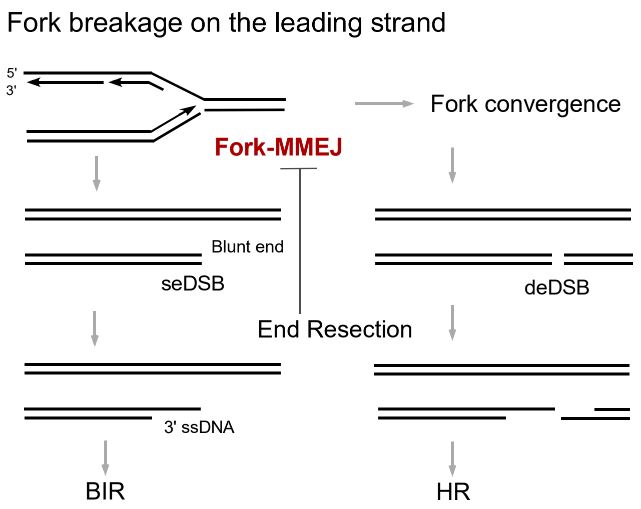


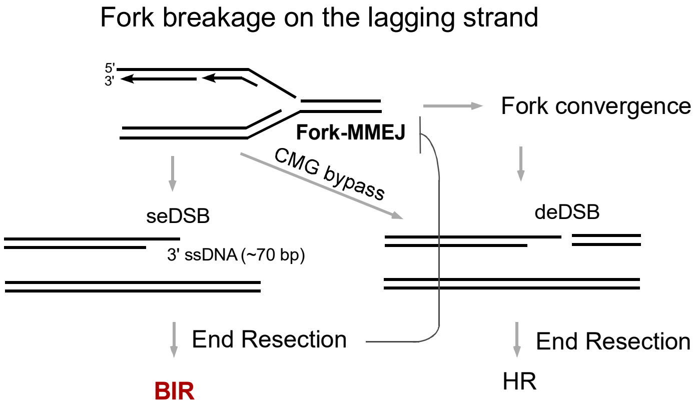


**Figure S18. Schematic drawings of the regulation of fork-MMEJ and BIR on broken leading and lagging strands.**

When forks are broken on the leading strands, seDSBs are blunt ended or contain up to 3 bp 5' overhangs, fork-MMEJ would immediately engage to repair seDSBs as BIR requires time-consuming end resection (top). When forks are broken on the lagging strands, seDSBs contain ~70 bp ssDNA overhangs, which channels the repair towards BIR, thereby reducing the use of fork-MMEJ (bottom). As end resection proceeds, repair of seDSBs is shifted towards BIR, and fork-MMEJ is suppressed. Both CMG-medited replication bypass on the broken lagging strands and fork convergence on both broken leading and lagging strands lead to the formation of deDSBs, which would be repaired by HR.

Figure S19

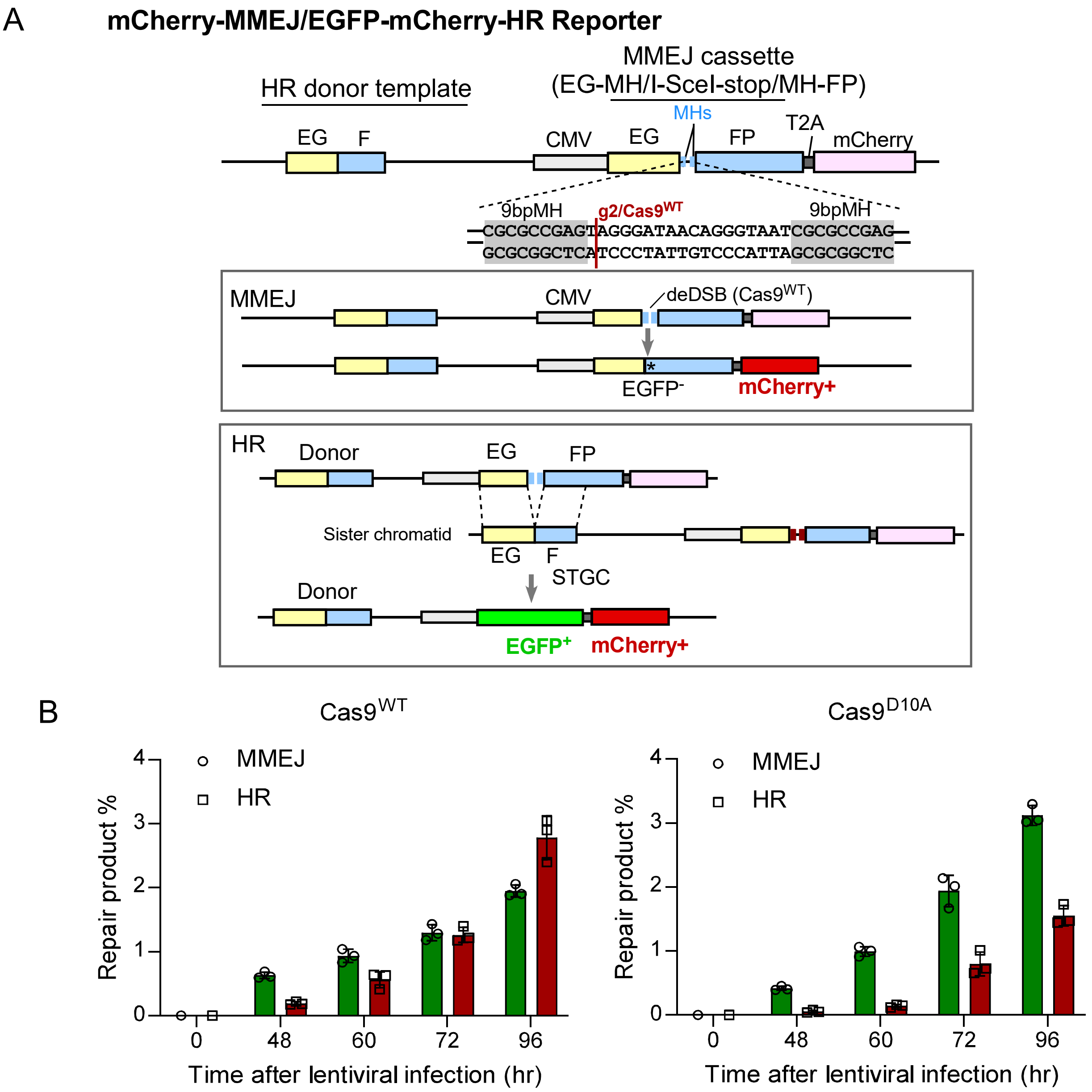


**Figure S19. Establishing a MMEJ and HR competition reporter to analyze the use of fork-MMEJ and HR to repair DSBs generated by Cas9$^{WT}$ and Cas9$^{D10A}$.**
(A) Schematic drawings of the mCherry-MMEJ/EGFP-mCherry-HR reporter (top) and the repair products by cMMEJ and HR after Cas9$^{WT}$ cleavage (bottom). In the MMEJ cassette (EG-MH/I-SceI-stop/MH-FP) of this reporter, EGFP was disrupted by replacing the chromophore residues T65/Y66/G67 with 9 bp (encoding R/A/E), followed by an intervening sequence containing the Cas9 cleavage site and stop codons, and then a duplication of the 9 bp serving as MHs. The HR donor template, the promoter-less EGF cassette, retains the original T65/Y66/G67 residues. If MMEJ is used, the open reading frame of EGFP-T2A-mCherry is restored by deletion of the intervening sequence between the two MHs. In the resulting MMEJ repair product, EGFP carries the mutated chromophore residues T65R/Y66A/G67E, remaining EGFP-negative. However, mCherry cDNA, fused to the EG/I-SceI-stop/FP recipient cassette via a T2A sequence, restores the correct open reading frame, producing red fluorescence. Consequently, MMEJ repair generates mCherry+/EGFP- cells. In contrast, HR repair copies the original EGFP chromophore residues T65/Y66/G67 from the donor template, restoring green fluorescence, thereby producing cells with both green and red fluorescence (EGFP+/mCherry+). (B) Time course experiments in U2OS (mCherry-MMEJ/EGFP-mCherry-HR) cells were performed to analyze MMEJ and HR frequency after infection with lentiviruses expressing gRNA2 along with Cas9$^{WT}$ or Cas9$^{D10A}$.

Figure S20

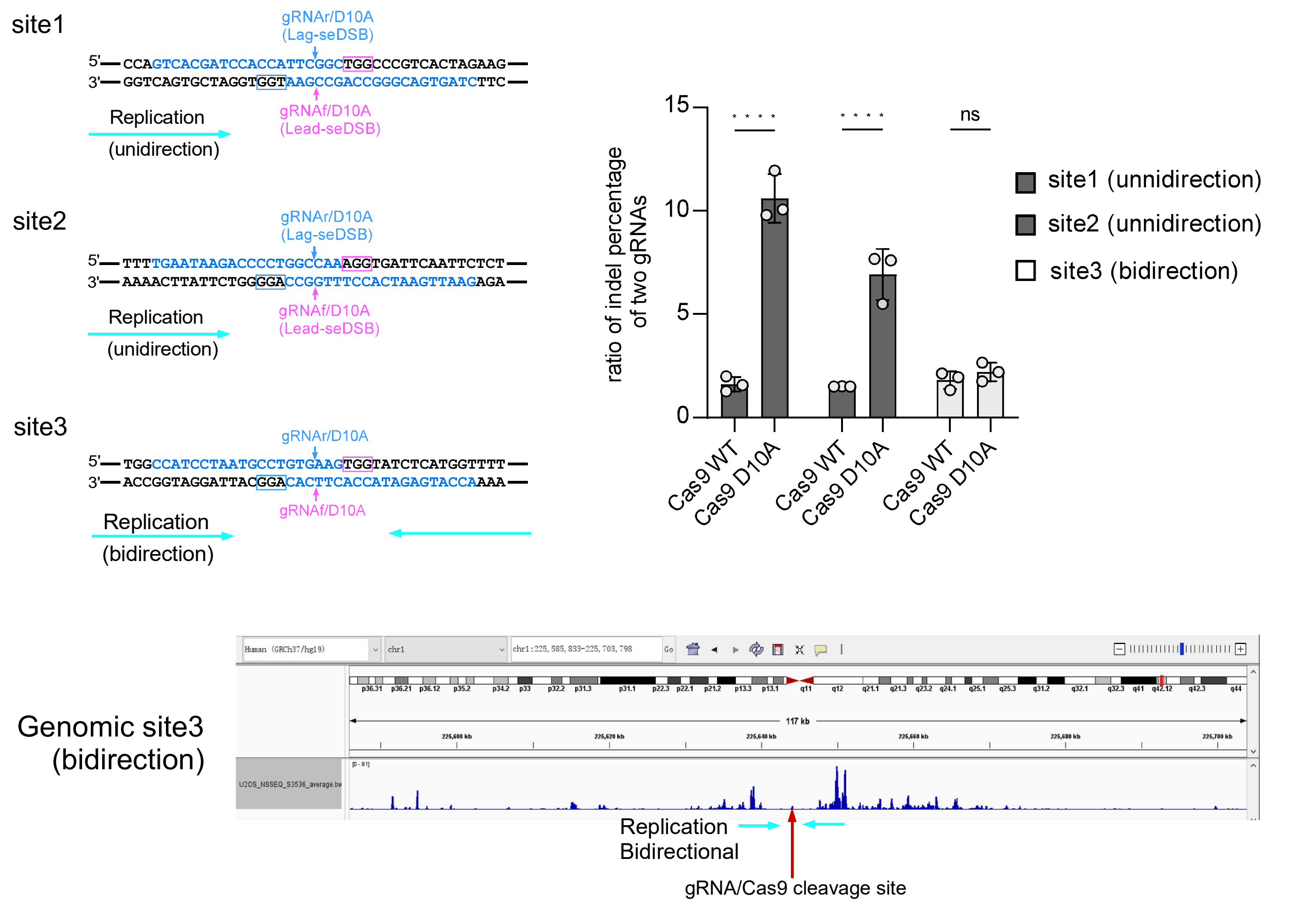


**Figure S20. Indels are accumulated more frequently on leading strands after Cas9n cleavage.**
The genomic sites, site 1 and site 2, with unidirectional replication described in Fig. S13, as well as site 3 (GRCh37, chr1, 225645808-225646057) with bidirectional replication were cleaved in U2OS cells by indicated gRNAs with Cas9$^{WT}$ and Cas9$^{D10A}$, followed by deep sequencing analysis of the cleavage sites. The gRNA cleavage sites in relationship with replication directions are shown (top left). The ratio of indel percentage between the two gRNAs (gRNAf over gRNAr) at each site after Cas9$^{WT}$ and Cas9$^{D10A}$ cleavage is shown (top right). For Cas9$^{D10A}$ cleavage, the ratio indicates the indel percentage of lead-seDSBs over lag-seDSBs. Nascent strand sequencing data from U2OS cells are shown for the genomic site 3 with bidirectional replication (bottom).

Figure S21

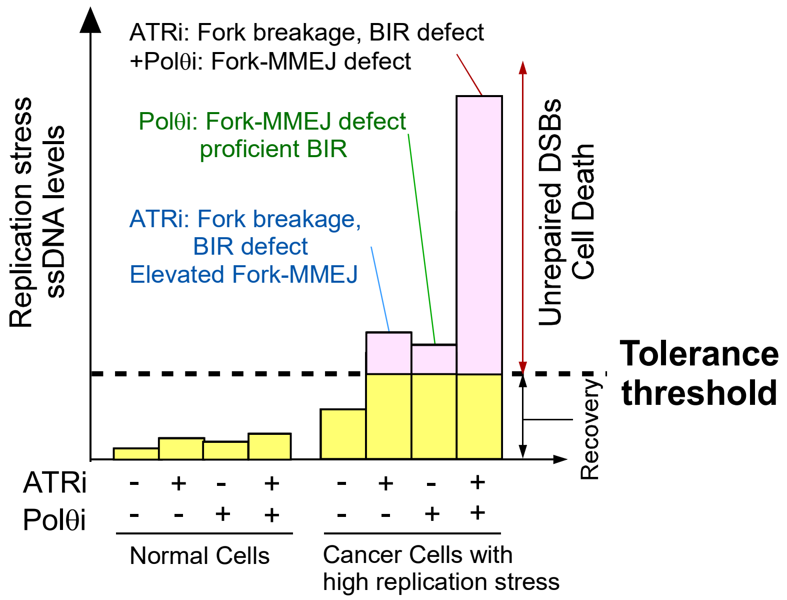


**Figure S21. Combined treatment using ATR inhibitor (ATRi) and Polθ inhibitor (Polθi) effectively eradicates cancer cells under high replication stress with low toxicity to normal cells.** Graphic depiction of the principle behind the different tolerance of normal cells and replication-stressed cancer cells to the treatment using ATRi and Polθi.

**Table S1. gRNA sequences**

| name | sequence |
|---|---|
| *AAVS1* gRNA | GTCCCTAGTGGCCCCACTGT |
| *ROSA* gRNA | ACTCCAGTCTTTCTAGAAGA |
| *Igh* gRNA | GATTCCACCATTGGCGCCCC |
| *POLQ* KO gRNA1 | CGCCGTTTCCTACTCCGGCG |
| *POLQ* KO gRNA2 | TCCCGAGAACGTGTCGGAGC |
| K120G gRNA1 | GAGTCTTCCCAGCACTTGT |
| K120G gRNA2 | CTTATTCTGAAGCGGGTTT |
| D2494P/E2495R gRNA1 | TGAAGAATCCTCCTCTCAT |
| D2494P/E2495R gRNA2 | GACGAGCTTTTATATGAAG |
| MMEJ gRNA1 | CGCGCCGAGTAGGGATAACA |
| MMEJ gRNA2 | TTACCCTGTTATCCCTACT |
| MMEJ gRNA4 | ACAAGACGCGCGCCGAGTA |
| MMEJ gRNA5 | TACAAGACGCGCGCCGAGT |
| MMEJ gRNA6 | ATAACAGGGTAATCGCGCCG |
| LBR-g1 | GCCGATGGTGAAGTGGTAAG |
| LBR-g2 | ACCTCTTACCACTTCACCAT |
| Igh-g2 | GCCACCTGGGCCCTACTCTG |
| Igh-g4 | GCTTCTACAGGCCTCAGAGT |
| Site1-gRNAf | CTAGTGACGGGCCAGCCGAA |
| Site1-gRNAr | GTCACGATCCACCATTCGGC |
| Site2-gRNAf | TGAATAAGACCCCTGGCCAA |
| Site2-gRNAr | GAATTGAATCACCTTTGGCC |
| Site3-gRNAf | CCATCCTAATGCCTGTGAAG |
| Site3-gRNAr | ACCATGAGATACCACTTCAC |

**Table S2. Sequences of DNA substrate and PCR**

| name | sequence |
|---|---|
| ssDNA1 | GCAAAGACCGCGGAAAGATCT |
| ssDNA2 | ACCGCGGAAAGATCT |
| ssDNA3 | GAGAGAAACTGAAGTATGACTAGATCTTTCCGCGGT |
| ssDNA4 | ACCGCGGAAAGATCTAGTCATACTTCAGTTTCTCTC |
| Mouse *POLQ* KO screening-F | ACTAGGTTGGGGTTCTCCCG |
| Mouse *POLQ* KO screening-R | CAGGATCTGGCCAACTTGGAA |
| K120G screening-F | TGCTGTACATACAGTGTGCACAGAG |
| K120G screening-R | CCTGTACCACAATACCCAACTGTCC |
| D2494P/E2495R screening-F | CTTAGGCACCTGTACCCACATAC |
| D2494P/E2495R screening-R | TTCAGAACTTGGAAACCTCAGACAAACG |
| *LBR* amplification-F | ACGACGCTCTTCCGATCTAT GCTGTCGTGGCTCAGAATTT |
| *LBR* amplification-R | AGTTCAGACGTGTGCTCTTCCGATCTCGCTATTCAGGTTAATT ATAGAAAATGCC |
| *Igh* amplification-F | ACGACGCTCTTCCGATCTAT GGCTTGGGACACATCCTGAG |

| | |
|---|---|
| *Igh* amplification-R | AGTTCAGACGTGTGCTCTTCCGATCTCGCACAAGGAGGTCCAC ACTGG |
| ddPCR-target-F | GAGCGCACCATCTTCTTCA |
| ddPCR-target-R | TGTGGCTGTTGTAGTTGTACTC |
| ddPCR-target-probe | /56-FAM/CATCGAGCT/ZEN/GAAGGGCATCGACTT/3 IABkFQ/ |
| ddPCR-control-F | ACAACAGCCACAACGTCTAT |
| ddPCR-control-R | GGGTGTTCTGCTGGTAGTG |
| ddPCR-control-probe | /5HEX/AGCAGAAGA/ZEN/ACGGCATCAAGGTGA/3 IABkFQ/ |
| UMI-For | p-GCATCGGAAGAGCGTCGNNNNNNNNNNACCAGCGCGA TAGGTTCGTCTTCTGCCGTATGC |
| UMI-Rev | CGACGCTCTTCCGATGC |
| Nest-left-1st | GCCCATATATGGAGTTCCGC |
| Nest-adaptor-1st | GCATACGGCAGAAGACGA |
| Nest-left-2nd | ACGACGCTCTTCCGATCTATGGCTACGCTGGACGGCGACGTAA AC |
| Nest-adaptor-2nd | AGTTCAGACGTGTGCTCTTCCGATCTCGGTTTCGCGAACCTAT CGCGCTGGT |
| Nest-right-1st | GCTGTCCATCTGCACGAGAC |
| Nest-right-2nd | ACGACGCTCTTCCGATCTATCTTGTAGTGCTCAGGTAGTGGTT GTC |
| P5 | AATGATACGGCGACCACCGAGATCTACAC[i5]ACACTCTTTCCC TACACGACGCTCTTCCGATCT |
| P7 | CAAGCAGAAGACGGCATACGAGAT[i7]GTGACTGGAGTTCAGA CGTGTGCTCTTC |
| 3’overhang-bottom | CCGAGCTCGAATTCACTGGCCGTCGTTTTA-3’ biotin |
| 3’overhang-top-0 nt | TAAAACGACGGCCAGTGAATTCGAGCTCGG |
| 3’overhang-top-10 nt | TAAAACGACGGCCAGTGAATTCGAGCTCGGTACCCGGGGA |
| 3’overhang-top-20 nt | TAAAACGACGGCCAGTGAATTCGAGCTCGGTACCCGGGGATC CTCTAGAG |
| 3’overhang-top-30 nt | TAAAACGACGGCCAGTGAATTCGAGCTCGGTACCCGGGGATC CTCTAGAGTCGACCTGCA |
| 3’overhang-top-50 nt | TAAAACGACGGCCAGTGAATTCGAGCTCGGTACCCGGGGATC CTCTAGAGTCGACCTGCAGGCATGCAAGCTTGGCGTAA |
| 3’overhang-top-70 nt | TAAAACGACGGCCAGTGAATTCGAGCTCGGTACCCGGGGATC CTCTAGAGTCGACCTGCAGGCATGCAAGCTTGGCGTAATCATG GTCATCTAGAAGCTG |
| MMEJ-ChIP-F | ACAACAGCCACAACGTCTAT |
| MMEJ-ChIP-R | GGGTGTTCTGCTGGTAGTG |
| LBR-PCR-F | ACGACGCTCTTCCGATCTATGGCTACGCTGTCGTGGCTCAGAA TTT |

| | |
|---|---|
| LBR-PCR-R | AGTTCAGACGTGTGCTCTTCCGATCTCGGTTTCGCTATTCAGGTTAATTATAGAAAATGCC |
| igh-PCR-F | ACGACGCTCTTCCGATCTATGGCTACGGCTTGGGACACATCCTGAG |
| igh-PCR-R | AGTTCAGACGTGTGCTCTTCCGATCTCGGTTTCGCACAAGGAGGTCCACACTGG |

**Table S3. Sequence of RNA**

| name | sequence |
|---|---|
| Caged gRNA2 for EGFP-MMEJ | rA[NPOM-dT][NPOM-dT]rArCrCrC[NPOM-dT]rGrUrUrArUrCrCrCrUrArCrUrGrUrUrUrUrArGrArGrCrUrArUrGrCrUrGrUrUrUrUrG |

**Table S4. Targeting sequence of shRNAs**

| name | sequence |
|---|---|
| Polθ | ACAACAACCCTTATCGTAAAG |
| LIG3 | AACCCTCCGTGCACTCAGCCG |
| MRE11 | GATGAGAACTCTTGGTTTATT |
| ATR | CGAGACTTCTGCGGATTGCAG |
| CtIP | GAGCAGACCTTTCTCAGTATA |
| RPA2 | CCTAGTTTCACAATCTGTTGT |
| PIF1 | GAAGACAGGTGCTCCGGAAGC |
| PCNA | GTGGAGAACTTGGAAATGGAA |
| MCM2 | GCTCTTCATACTGAAGCAGTT |
| BLM | GAGCACATCTGTAAATTAATT |
| BRCA1 | TGATAAAGCTCCAGCAGGAAA |
| RAD51 | CTAATCAGGTGGTAGCTCAAG |
| EXO1 | ACTCGGATCTCCTAGCTTTTG |
| DNA2 | GTGCTAAACCGTGAAGCAAGA |
| RHINO | GAAGCTGAGCAGAAGCCAATT, GCACTTCAGAGCTTACCTTAT |

**Table S5. Primer sequence for RT-qPCR**

| name | sequence |
|---|---|
| POLQ | CACACTGCTACAGGACGAATAA and AGGTGGGCTTTCTCCTACTA |
| MRE11 | GTAACCCAAGCCATACAAAGC and ACCTCCACTATAGTCCACTCG |
| LIG3 | CCACTAACCTTCCCCAACTC and CCCTTATTCTCTTCACTGCTACC |
| CtIP | GAAATTGGCTTCCTGCTCAAG and TTTTGGACGAGGACAAGGATC |
| ATR | CCTTGAACATGAAAGCCTTGG and CCTGAGTGATAACAGTAGACAGC |
| PIF1 | CGAGCCTAGCACAGAAGCC and CCCAGGATTCGCTTTAGCAG |
| PCNA | CTAGCCTGACAAATGCTTGC and AGGAAAGTCTAGCTGGTTTCG |
| RHINO | CTTGTTCCCAGCCTACCAGA and TGTGGAAACTTGGAGGTGGT |
| HPRT | CTGGCGTCGTGATTAGTGAT and CTCGAGCAAGACGTTCAGTC |
